# Non-genetic control of cell plasticity and phenotypic cell states is mediated by cytoplasmic IncRNA nucleated signalling pathways

**DOI:** 10.1101/2024.11.08.622637

**Authors:** Y Shlyakhtina, B Bloechl, KL Moran, A Maslakova, RB Johnson, AF Munro, NO Carragher, MM Portal

**Affiliations:** Cell Plasticity & Epigenetics Lab, Cancer Research UK – Manchester Institute, The University of Manchester, M20 4BX, Manchester, UK; Cell Plasticity & Epigenetics Lab, Cancer Research UK – Scotland Institute, University of Glasgow, G61 1BD, UK; Edinburgh Cancer Research, Institute of Genetics and Cancer, University of Edinburgh, EH4 2XR, UK; Cancer Research UK Scotland Centre, UK

**Author notes:** Contributed equally to this work (Y Shlyakhtina, B Bloechl, KL Moran).

**Keywords:** cell plasticity, epigenetics, RNA sequencing, transcriptomics

## Abstract

Cell plasticity, the ability that cells display to rapidly adapt to environmental cues, is thought to be encoded in non-genetic information reservoirs. Although the notion is widely acknowledged, the molecular details underlying this phenomenon remain largely concealed. Herein, we show that clonal cell populations inherently display multiple co-existing metastable gene expression states that co-segregate with various phenotypic outputs. Moreover, we provide primer evidence suggesting that transcriptome states are inherited, dynamically interconvert and determine phenotypic output upon a variety of biological cues, not as a result of transcriptional shifts, but rather through yet unidentified post-transcriptional mechanisms. Remarkably, among phenotypically divergent clonal cell populations enriched in subsets of transcriptome states, we identified a peri-nuclear cytoplasmic structure (Signal Integration Portal – SIP) where state-specific lncRNAs, proteins harbouring intrinsically disordered regions and various active signalling pathways converge. Herein, we propose that SIP-condensates act as nucleating reservoir of non-genetic information at the crossroads of cell plasticity and non-genetic heterogeneity where they integrate intra- and extracellular inputs thereby moulding phenotypic output.

## Introduction

The wide field of genetics has revolutionised the manner in which we approach biological problems and, even though its remarkable relevance cannot be contested, hundreds of observations over the years suggest that the non-genetic compartment is of equal importance to define life as we know it and, therefore, should not be neglected^1-3^. Along those lines, following the development and refinement in next generation sequencing (NGS) technology and its application to single cell analysis, we can finally begin to query the non-genetic compartment with unprecedented detail in a robust and quantitative manner. It is in that context that cell plasticity – the ability of a single genotype to produce a variety of phenotypes – has emerged as a central player in various and diverse fields of study ranging from microbiology to cancer research^4-7^. Notably, within the plethora of non-genetic information carriers present in cells, the study of the transcriptome has transformed basic and biomedical research. In particular, the development of single-cell RNA-Sequencing (scRNA-Seq) unravelled that even genetically homogeneous cell populations inherently display heterogeneous transcriptome profiles^8^.

Strikingly, though transcriptome heterogeneity is now considered to be prevalent in most biological settings, our understanding of how transcriptome states are established, maintained and inherited, and the molecular mechanisms underlying their potential to reshape into novel transcriptome arrangements remains fragmentary. Although attempts to unravel the molecular details underlying this phenomenon have been met with modest success^9-13^, the field of system dynamics brought forward a theoretical framework which anchored on Waddington’s epigenetic landscape^14-16^. It suggests that each metastable – attractor – state embedded in a gene regulatory network can be represented as low-energy wells in a rugged quasi-potential landscape towards which unstable transient states are attracted to^14,17,18^. Along those lines, and in partial agreement with the aforementioned theoretical framework, our reported data showed that non-genetic cell heterogeneity in terms of meta-stable transcriptome states is not randomly established but follows precise phylo-(epi-)genetic linked inheritance patterns, generating traceable cell-ancestry trees within isogenic and genetically stable experimental models (Barcode Decay Lineage tracing, BdLT-Seq^8^). Moreover, intrapopulation lineage-linked transcriptome states are perpetuated for several generations fuelling population heterogeneity and thus suggesting the existence of a “molecular memory” retained through clonal propagation/divergence.

Herein, we show that by subcloning a genetically stable clonal founder cell population we isolate ancestry trees that are enriched in specific subsets of transcriptome states which correlate with the fate of individual cells in response to intracellular and extracellular cues. Moreover, we provide primer evidence suggesting that for a given cellular population, the degree of phenotypic plasticity is anchored to the transcriptome state of each individual cell and its associated lineage-linked restricted plasticity. Finally, we describe the existence of a cytoplasmic compartment (Signal Integration Portal - SIP) that nucleates long non-coding RNAs, proteins containing intrinsically disordered regions (IDR) and active components of multiple signalling pathways that co-segregate with subsets of transcriptome states and whose variable composition correlates with phenotypic output and transcriptome-linked non-genetic heterogeneity.

Collectively, our data supports the notion that the perpetuation of non-genetically heterogeneous cell populations in physiological and pathological conditions, as well as in response to biological and/or therapeutic cues, is not only dictated by the overall cellular architecture (i.e. epithelial, mesenchymal, tissue of origin) and molecular characteristics (i.e. genetic landscape, bulk transcriptome profiles) of the population but also strongly relies on the non-genetic molecular identity (SIP complexity and composition) and plasticity of its intrapopulation phylo-epigenetic lineages.

## Results

### Clonal cell systems retain a molecular memory of past cell states

We have recently shown that by applying Barcode Decay Lineage tracing (BdLT-Seq) to a given cellular system of choice we can unravel metastable gene expression state dynamics in a lineage-linked manner^8^. BdLT-Seq data also suggested that the gene expression states observed within a cellular population are not randomly generated but rather constrained by unknown molecular mechanisms and could potentially drive phenotypic output.

Therefore, to unravel the molecular mechanisms underlying gene expression state transitions and its link with phenotypic output, we first evaluated a clonal founder population of HA1ER cells (F12 - HEK cells immortalised with hTERT/SV40ER and transformed by HRAS^G12V 19^) and two stable subclones derived from it (1F8 and 1C9) previously reported as diverging at their transcriptome level^8^. Even though the three clonal populations (F12, 1F8 and 1C9) were essentially indistinguishable for a plethora of cellular parameters (cell proliferation, ploidy, cell cycle profile, morphology. Supplementary Fig. 1), scRNA-Seq (Chromium 10x) revealed that, after data processing to mitigate clustering artefacts due to cell cycle progression and abnormal mitochondrial counts (Proxy for cell death. Seurat v4.0), gene expression states dramatically differed between the founder population and its subclones (subclones 1C9 and 1F8; Figs. 1a, 2a, Supplementary Figs. 2a, 3a). Interestingly, those transcriptomic differences are further propagated down the lineage when the first subclones obtained from the founder clone are further subcloned, indicating a certain degree of inheritance based on the lineage-linked plastic capacity of the transcriptome states (Subclones 1F8, 3C5 and 1E8; Supplementary Fig. 4, left. Subclones 2A6, 2B12 and 1F5; Supplementary Fig. 4, right). Remarkably, whole exome (WES, 100x coverage) and whole genome sequencing (WGS, 30x coverage) for clones F12, 1F8 and 1C9 did not reveal any genetic difference between them (variance per site (nucleotide position) <0.02, Supplementary Figs. 5, 6) thus suggesting that the observed gene expression state divergence is encoded and propagated by non-genetic networks.

**Fig. 1.**
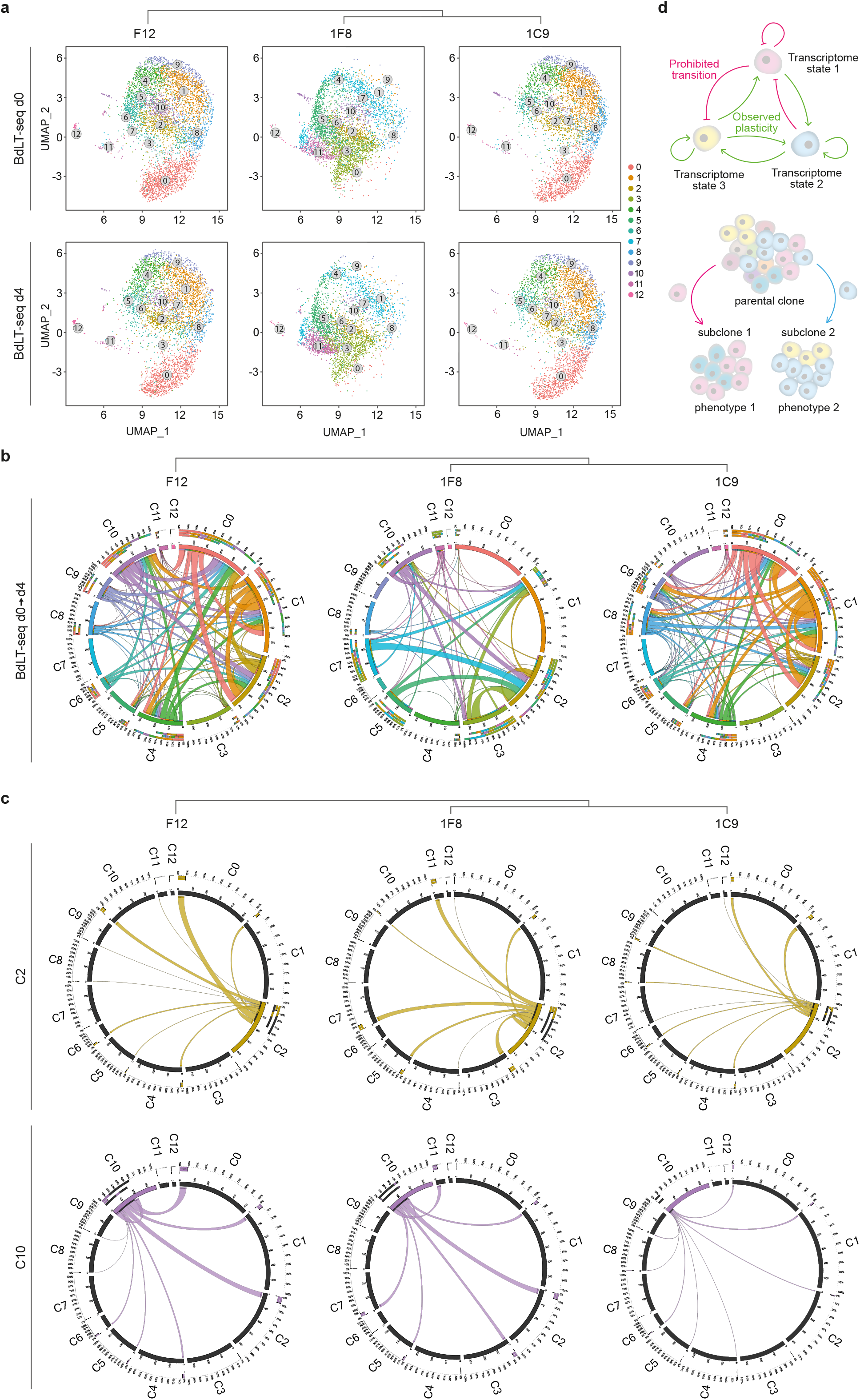
Non-genetic clonal divergence dynamics predicts transcriptomic state bottlenecks. **a**. HA1ER genetic anchor (F12) and two diverging subclones (1F8 and 1C9) were subjected to BdLT-Seq coupled to scRNA-Seq and traced for 4 days. UMAP plots depict transcriptome states at day 0 and 4 of tracing. Individual states are depicted in colour and numbered. **b**. Chord diagrams representing transcriptome state dynamics for cells belonging to a particular state/cluster at the beginning of tracing and their divergence after 4 days for clone F12 and subclones 1F8 and 1C9. Detected clusters are depicted (C0 to C12) and integrate the collapsed behaviour of all cells that belong to each gene expression state. Origin clusters are shown in different colours (Day 0) and chords represent the endpoint cluster association (Day 4). **c**. Chord diagrams representing individual examples of transcriptome divergence. C2 and C10 divergence is depicted. Lineage relationship of the clones is depicted for **a, b** and **c. d**. Schemes representing transcriptome divergence conceptual framework.

**Fig. 2:**
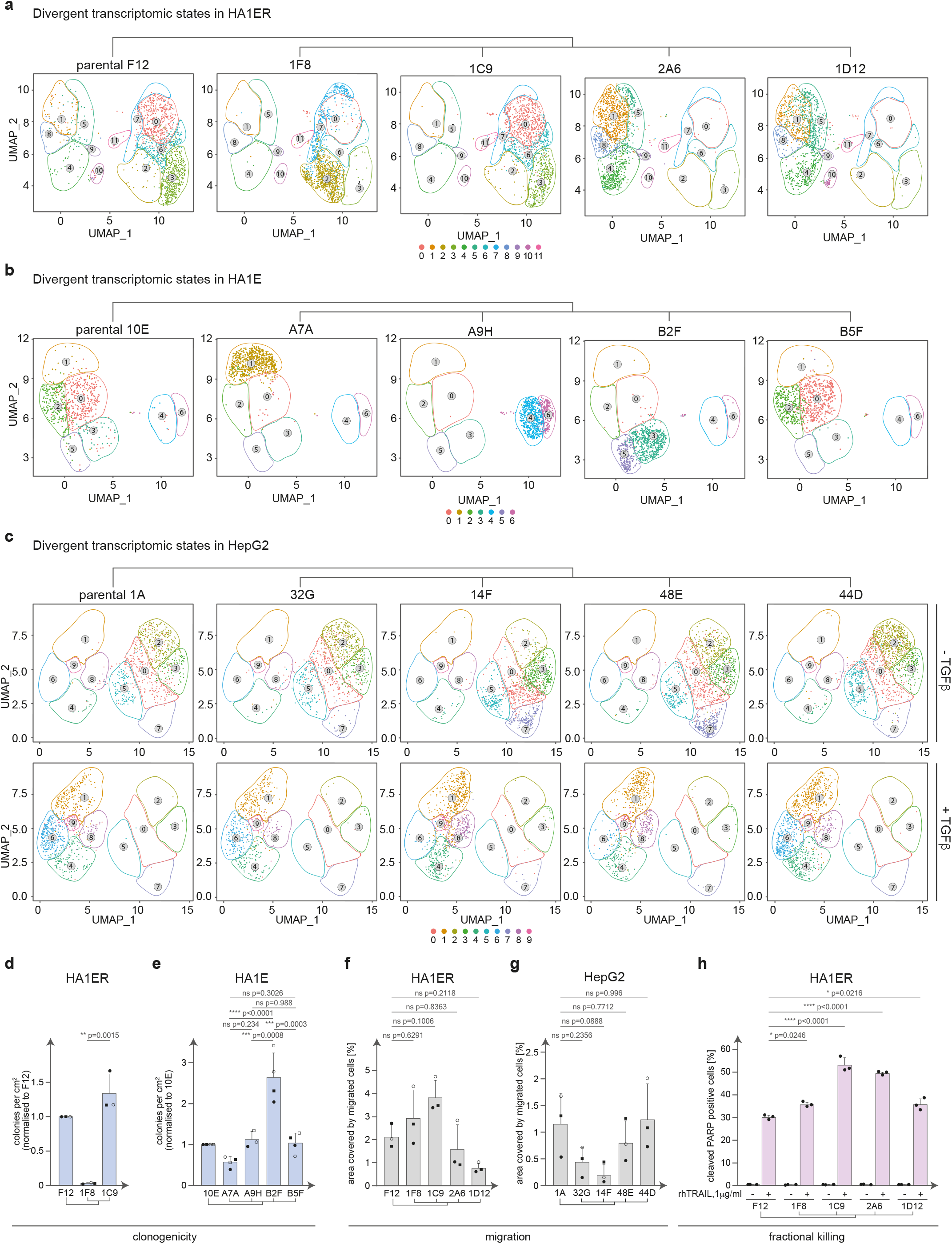
Clonal cell systems retain a molecular memory of past cells states. **a, b** and **c**. Genetic anchors from the HA1ER (F12), HA1E (10E) and HepG2 (1A) and divergent subclones from each cell system (HA1ER – 1F8, 1C9, 2A6 and 1D12; HA1E – A7A, A9H, B2F and B5F; HepG2 – 32G, 14F, 48E and 44D) were subjected to scRNA-Seq. UMAP plots represent transcriptome states and their divergence among analysed subclones. Individual states are depicted in colour and numbered. c. HepG2 clonal system was subjected to either vehicle (-TGFβ) or TGFβ treatment (+TGFβ) and processed for scRNA-Seq. UMAP plots represents clonal divergence among clones and between conditions. Lineage relationship is depicted for **a, b** and **c. d**. Clonogenicity assay in HA1ER clonal system (F12, 1F8 and 1C9). Histogram represents the number of colonies per cm^2^ normalised to F12 (genetic anchor). Histograms represent the mean value ± standard deviation (SD) of 3 independent experiments performed in 4-6 replicates. Statistical significance between 1F8 and 1C9 was assessed by two-tailed unpaired t-test with Welch correction. Individual data points are shown. **e**. Clonogenicity assay in HA1E clonal system (10E, A7A, A9H, B2F and B5F) upon RAS variants transduction. Histogram represents the number of colonies per cm^2^ normalised to 10E (genetic anchor). Histograms represent the mean value ± standard deviation (SD) of 4 (3 for A9H) independent experiments performed in triplicate. Statistical significance between all subclones was assessed by ordinary one-way ANOVA followed by Tukey’s multiple comparisons test. Individual data points are shown. **f**. HA1ER clonal system (F12, 1F8, 1C9, 2A6 and 1D12) was subjected to transwell migration assay. Histograms depict the mean area covered by migrated cells ± standard deviation (SD) of 3 independent experiments performed in triplicate/duplicate. Statistical significance of the subclones compared to F12 was assessed by ordinary one-way ANOVA followed by Dunnett’s multiple comparisons test. Individual data points are shown. **g**. HepG2 clonal system (1A, 32G, 14F, 48E and 44D) was subjected to transwell (migration) assay. Histograms depict the mean area covered by migrated cells ± standard deviation (SD) of 3 independent experiments performed in triplicate. Statistical significance of the subclones compared to 1A was assessed by ordinary one-way ANOVA followed by Dunnett’s multiple comparisons test. Individual data points are shown. Clonal relationship is depicted for all panels. **h**. Histograms depicting TRAIL-induced apoptosis (1 μg/ml rhTRAIL for 2h) in HA1ER clonal system (F12, 1F8, 1C9, 2A6 and 1D12) as determined by cleaved PARP staining analysed by flow cytometry. Histograms show the mean value⍰± ⍰standard deviation (SD) of 3 biological replicates (∼20,000 cells per condition). Statistical significance of the subclones compared to F12 was assessed by one-way ANOVA followed by Dunnett’s multiple comparisons test. Individual data points are shown.

To further verify this possibility, we performed BdLT-Seq to determine if transcriptome plasticity was differential between the founder clone F12 and subclones 1F8 and 1C9 and revealed that indeed, the founder clone and its derived subclones display not only a divergent state composition but also differential basal transcriptome plasticity (Fig. 1b, c and Supplementary Fig. 2b). For example, after four days, the progeny of cells in cluster 2 of clone F12 could be traced to every cluster except clusters 3, 7 and 12, with the majority transitioning to clusters 0 and 10. On the other hand, cells from cluster 2 of subclone 1F8 mainly diverged to clusters 3, 7 and 11, whereas cells from cluster 2 of subclone 1C9 could be traced primarily to clusters 0 and 1 but not to clusters 3, 10 and 11 (Fig. 1c, Supplementary Fig. 2b).

Prompted by these results, we hypothesised that any isogenic and genetically stable cellular system amenable to cloning could display similar divergent transcriptome behaviour when analysing subclonal populations at the single-cell level and concordantly could be exploited to shed light onto the molecular mechanism underlying the genesis/establishment and maintenance of metastable transcriptome state dynamics. To pursue this hypothesis, we built three different clonal cellular systems (HA1ER, described above; HA1E, HEK cells immortalised with hTERT/SV40ER; HepG2, Hepatocellular carcinoma cells) by generating in the first step a clonal founder population and then subcloning each founder population to obtain transcriptome state diverging subclones whilst maintaining isogeneity and enabling lineage-linked comparisons (Fig. 1d).

After initial phenotypic characterisation of each clone (cell proliferation, ploidy, cell cycle profile and morphology. Supplementary Figs. 1) and determination of their genetic makeup which did not show any significant differences (WES for all clones and WGS for HA1ER clones F12, 1F8 and 1C9 and HepG2 clones 1A, 14F and 44D, variance per site <0.02 for all, Supplementary Figs. 5, 6), the transcriptome states of the founder clone and the four subclones from each system were analysed by scRNA-Seq (10x Genomics Chromium 3’, Fig. 2a-c). Strikingly, scRNA-Seq data analysis determined that for the HA1ER system, 3 of the 4 subclones (1D12, 2A6 and 1F8) displayed transcriptome state re-arrangements when contrasted to the founder population F12, whilst the remaining subclone (1C9) exhibited similar transcriptome states as the founder albeit displaying altered state representation (Fig. 2a, Supplementary Fig. 3a). Similarly, when populations from the HA1E lineage were analysed, 3 out of the 4 subclones (A7A, A9H, B2F) also displayed vast transcriptome re-arrangements when compared to the founder clone 10E whereas subclone B5F was the most comparable to the parental clone (Fig. 2b, Supplementary Fig. 3b).

To further explore whether transcriptome complexity (state representation) would affect phenotypic output throughout dynamic processes, we took advantage of the capability of HepG2 cells to activate a TGFβ-dependent transcriptional programme driving the epithelial-to-mesenchymal transition (EMT). Briefly, we treated the founder HepG2 population (1A) and the four established subclones (32G, 14F, 48E and 44D) with TGFβ to promote EMT (Supplementary Fig. 7a) and evaluated the representation of gene expression states by scRNA-Seq in vehicle-treated (-TGFβ) or 4 days TGFβ-treated populations (+TGFβ). As observed in HA1E and HA1ER cells, when TGFβ–naïve HepG2 cells were assessed, significant differences in the transcriptomic states between subclones could be observed when compared to the founder clone 1A (Fig. 2c, Supplementary Figs. 3c, 8). For instance, whereas in the naïve founder clone (1A) virtually no cells could be detected belonging to cluster 7, subclone 14F and 48E displayed a pronounced increase in the cellular representation in this cluster (Fig. 2c, Supplementary Fig. 3c). As expected, upon EMT induction, a considerable gene expression shift was observed when either the founder clone or the subclones were subjected to TGFβ treatment. Strikingly, the initial differences observed in the TGFβ–naïve subclones propagated into the states identified upon TGFβ treatment. Notably, the subclones displaying an enrichment of cells in cluster 7 in naïve conditions (14F and 48E) displayed a reduced representation of cells occupying cluster 6 following EMT compared to the other clones (Fig. 2c, Supplementary Fig. 3c).

To take this further and given that the HepG2-based TGFβ-induced EMT model system also enables us to assess the mesenchymal-to-epithelial transition (MET) by the removal of TGFβ from the cultures, we took advantage of this possibility and explored transcriptomic state re-arrangement upon EMT reversion using scRNA-seq (Supplementary Fig. 8). We observed that the founder clone 1A and its most similar subclone at the transcriptome level, 44D, reverted to occupy similar transcriptome states as the initial TGFβ–naïve population; however, their state representation was significantly altered with cluster 6 becoming the most predominant (Supplementary Fig. 8). Notably, clone 14F which was the most dissimilar from the founder clone 1A, took a completely different path upon EMT reversion, as cluster 1 and 8 were overpopulated with only a minority of cells occupying other states (Supplementary Fig. 8). Together, this data supports the notion that even when a transcription factor mediated gene expression programme is enforced by external cues, the initial cell states are largely inherited and may impact the phenotypic fate of a clonal population.

### Isogenic subpopulations enriched in different transcriptomic states are associated with distinct biological phenotypes

Following up on these observations and our previously reported data^8^, we set out to evaluate several phenotypic outputs relevant to each particular cellular system to compare the phenotypic responses between subclonal populations enriched in distinct transcriptomic states. First, we took advantage of the well-characterised HA1ER cell system that bears the capability to grow in 3D soft agar matrices, where its growth in anchorage-independent conditions serves as *in-vitro proxy for c*ellular transformation^19-23^.

Interestingly, though no major differences could be observed in the proliferation rates of the analysed clones (1D12, 1C9, 2A6 and 1F8) and their founder clone (F12) in 2D growth (Supplementary Fig. 1a), challenging these clonal populations to grow in anchorage-independent conditions resulted in variable phenotypic outputs as subclone 1C9 (most similar to the founder population F12 at the transcriptome state level), grew more efficiently under these conditions (Fig. 2d). Notably, HA1ER subclones 2A6 and 1F8 (most dissimilar at the transcriptome state level to founder clone F12), have almost lost the capability to grow in 3D settings albeit being genetically identical to the founder cell population (Fig. 2d, Supplementary Fig. 7c). Strikingly, subclone 1D12 grew to a similar extent to the parental clone F12 in 3D conditions despite being the most similar at the transcriptome level to subclone 2A6 which failed to grow efficiently under the same conditions (Fig. 2a, Supplementary Fig. 7c), thus suggesting that transcriptome organisation plays a role in modulating phenotypic output but is not the sole determinant. Interestingly, when the second round of cloning from the F12 lineage was analysed, the subclones derived from 1F8 and 2A6 (1E8, 3C5; 1F5, 2B12, respectively) did follow the phenotype displayed by their respective founder clone and failed to grow in 3D soft agar (Supplementary Fig. 7c), thus indicating that phenotypic divergence is propagated across several generations and is potentially restricted by the plasticity of the parental clone.

To delve further into our findings, we took advantage of the fact that immortalised HA1E cells (hTERT + SV40ER), are susceptible to transformation by overexpressing a RAS oncogene^19,23^. To evaluate whether transcriptomic divergence can result in differences in transformation potential, we prepared a multiplexed lentiviral library containing 26 RAS variants from the three main families H-RAS, N-RAS and K-RAS (wild type and 7-8 mutants for each family; Supplementary Fig. 7f), transduced each of the HA1E subclones (A7A, A9H, B2F and B5F) and their founder (10E) and then subjected each population to anchorage-independent growth conditions in 3D soft agar to evaluate their susceptibility to malignant transformation (Fig. 2e). Notably, subclones A9H and B5F grew to a similar extent in 3D soft agar cultures as the founder clone (subclone B5F most similar at the transcriptome level to the founder clone 10E. Fig. 2b, e). However, two subclones that dramatically diverge at the level of gene expression states from the founder clone, A7A and B2F, either grew only to half the extent of the founder or grew two times more efficiently (Fig. 2b, e) thus suggesting that the transformation capacity of a given cell is somehow modulated by intrinsic molecular mechanisms encoded within state-specific non-genetic networks.

Concordantly, when HepG2 clonal populations were analysed for their capability to grow in soft agar in naïve conditions or upon TGFβ treatment, once again a divergence in their individual growth capabilities could be observed as all the subclones analysed grew less efficiently than the founder clone in naïve conditions, yet displaying variability among them (Supplementary Fig. 7d). Interestingly, whereas most clones almost lost the ability to form colonies in soft agar upon TGFβ treatment (1A, 32G, 44D), subclones 14F and 48E, which both display a divergent transcriptome profile compared to the other three clones grew more efficiently in 3D soft agar compared to the other subclones (Supplementary Fig. 7e).

Interestingly, when we subjected clonal populations of HA1ER and HepG2 and their corresponding subclones to a transwell migration assay, once again we observed profound phenotypic variability that did not segregate with the observed growth in 3D soft agar but was albeit different between the subclones analysed and their respective founder clone (Fig. 2f, g, Supplementary Fig. 7b). Moreover, as HA1ER cells are susceptible to trigger apoptosis upon treatment with Tumour necrosis factor-related apoptosis-inducing ligand (TRAIL), we also tested our clonal HA1ER models for their capability to withstand TRAIL challenge. Notably, although TRAIL treatment induced fractional killing in all clones, the extent to which apoptosis was triggered differed between all subclones (1F8, 1C9, 2A6 and 1D12) and the founder clone (F12) (Fig. 2h). Collectively, our data reveals notable phenotypic differences between genetically identical subclonal populations that appear to be profoundly impinged by their underlying (state-specific) non-genetic makeup.

### Gene expression states are post-transcriptionally established and maintained

Our data supports the notion that gene expression states are highly dynamic and shape phenotypic output. These observations encouraged us to explore the underlying molecular mechanisms contributing to transcriptome state dynamics, maintenance and divergence.

To address these, we set out to investigate whether changes in active transcription were driving transcriptome divergence by initially analysing clones from the HA1ER systems (F12 lineage, subclones 1F8 and 1C9). To do so, we performed SLAM-Seq^24,25^ (shown to provide high-resolution transcriptome dynamics data either at the transcriptional or RNA stability levels) to determine the rate of active transcription and its contribution to steady-state RNA levels (defined as snapshot RNA pool) in both clones. After sequencing, obtained reads were mapped to hg38 using a T>C aware aligner (NextGenMap^26^) and further processed through the SLAM-DUNK pipeline^27^. Next, following the determination of T-to-C conversion rates in each sequencing library as quality control (0.52-0.75 for all samples, robustly above background T-to-G conversion rate of 0.07, Supplementary Fig. 9a), we proceeded to perform a pairwise comparison between both clones analysed (1F8 and 1C9) where we applied the paired Beta binomial distribution (active transcription vs steady-state levels) to determine whether differences at the nascent RNA level could be observed^25^. Strikingly, we did not detect significant differences in the nascent RNA pool (FDR<0.05, FC>2) whilst at the steady-state level, we observed 217 transcripts differentially expressed between both clones (FDR<0.01, FC>2; Fig. 3a), suggesting that the observed differences in transcriptome state levels do not rely on transcriptional processes, at least in the cellular system evaluated here.

**Fig. 3.**
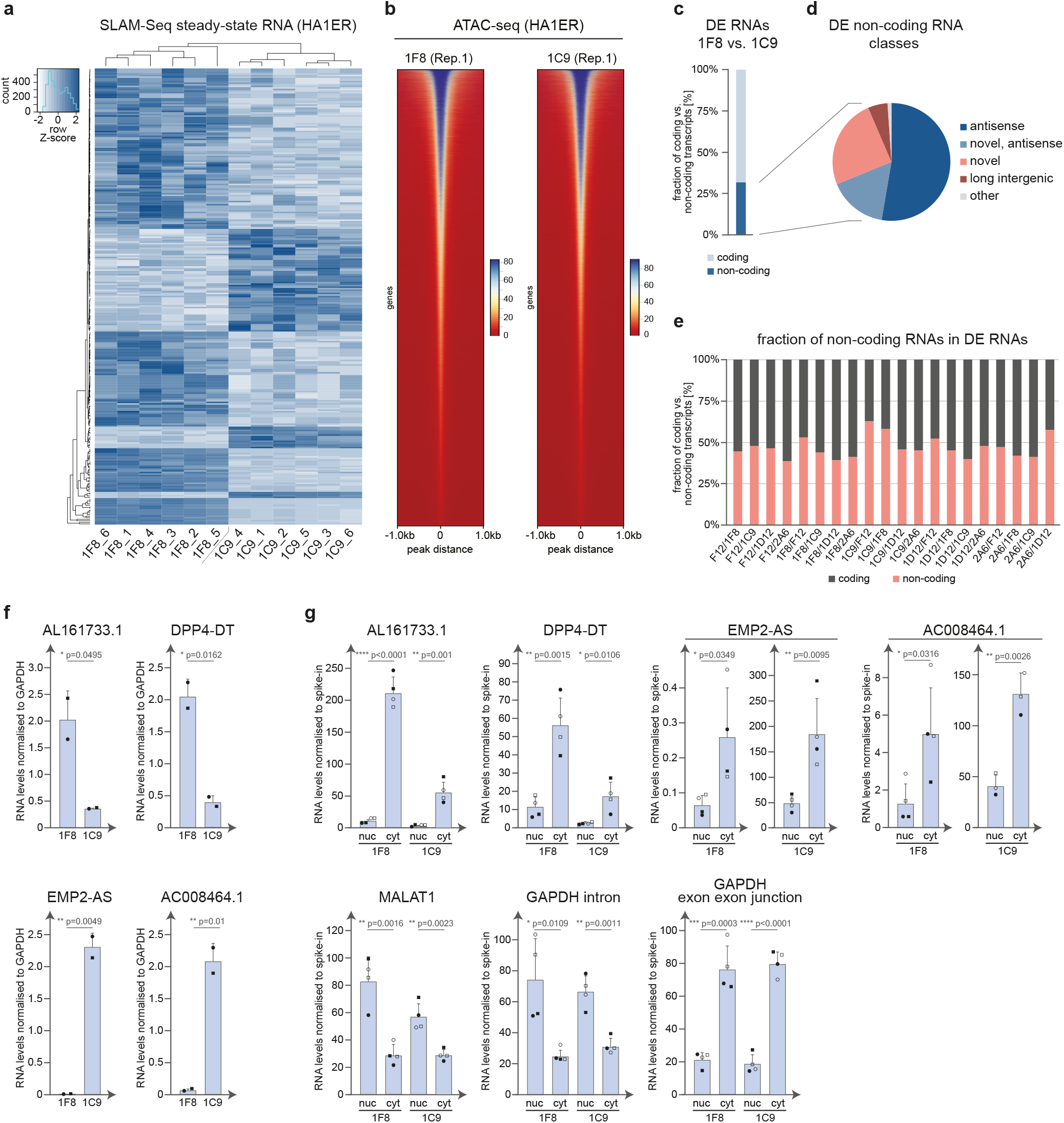
LncRNAs are major contributors to clonal divergence. **a**. Diverging clones 1F8 and 1C9 (HA1ER system) were subjected to SLAM-Seq analysis. Heatmap represents differentially expressed transcripts at the steady-state level (FDR<0.01; FC>2). No differences were observed in active transcription (FDR<0.05). Scale bar is depicted. Results represent 2 independent experiments performed in triplicate. **b**. ATAC-Seq was performed on HA1ER clones 1F8 and 1C9 to evaluate accessible chromatin regions. A representative heatmap displaying the identified peaks as determined by MACS using FDR<0.05 for each clone is depicted. Peaks distance ± 1Kb. Scale bar is depicted. **c**. Differentially expressed transcripts between 1F8 and 1C9 extracted from SLAM-Seq data in **a** were annotated based on their coding capacity. Histogram represents the fraction of coding vs non-coding transcripts. **d**. Differentially expressed non-coding RNAs were annotated (g:Profiler version e111_eg58_p18_f463989d^92^, Ensembl release 105^90^, LNCipedia v5.2^93^). Pie chart depicts the fraction of differentially expressed non-coding RNAs falling into known categories. **e**. Pairwise comparison of differentially expressed transcripts (FC>3 up-regulated) in scRNA-Seq datasets of HA1ER clones depicted in Fig. 2a and their annotation on coding vs non-coding potential. **f**. A subset of identified differentially expressed lncRNA transcripts between HA1ER clones 1F8 vs 1C9 were validated by reverse transcription followed by qPCR (RT-qPCR). Histograms depict AL161733.1, DPP4-DT, EMP2-AS and AC008464.1 transcript levels normalised to GAPDH levels. Results represent the mean ± standard deviation (SD) of 2 independent experiments performed in technical triplicates. Statistical significance was assessed by two-tailed unpaired Student’s t-test. Individual data points are depicted. **g**. A subset of differentially expressed lncRNA transcripts between 1F8 vs 1C9 were analysed for their intracellular localisation by subcellular fractionation (nucleus (nuc) vs cytoplasm (cyt)) and their levels evaluated by RT-qPCR. Histograms depict RNA levels of each fraction upon normalisation to the spike-in (*in-vitro transcribed* partial eGFP transcript). Data for AL161733.1, DPP4-DT, EMP2-AS and AC008464.1 is depicted. MALAT1 (enriched in nucleus), GAPDH intron (enriched in nucleus) and GAPDH exon-exon junction (enriched in cytoplasm) are shown as fractionation controls. Data represents the mean ± standard deviation (SD) of 3-4 independent experiments performed in technical triplicates. Statistical significance was assessed by two-tailed unpaired Student’s t-test. Individual data points are depicted.

Prompted by those results, we wondered whether our findings were a general phenomenon or if it was an isolated case only valid for the 1F8 vs 1C9 comparison. Thus, we performed SLAM-Seq analysis in the three clonal systems that we have previously shown (HA1ER, HA1E and HepG2) and compared all subclones with their corresponding founder clone for each system (T-to-C conversion rates: HA1ER 0.23-0.36, HA1E 0.18-0.34, HepG2 0.045-0.22, T-to-G background conversion rate: n.d., Supplementary Figs. 9b, 10). Strikingly, for most of the comparisons, no significant differences in nascent transcription were observed (HA1ER: F12/1C9, HA1E: 10E/A7A, 10E/B5F, HepG2: 1A/14F, 1A/48E; Supplementary Fig. 10) whereas when transcriptional differences were detected (HA1ER: F12/1F8, F12/2A6, F12/1D12, HA1E: 10E/A9H, 10E/B2F, HepG2: 1A/44D, 1A/32G), they only had a slight-to-none impact on steady-state RNA levels and quite often the observed changes in steady-state RNA (up or downregulation) did not follow the transcriptional trend (Supplementary Fig. 10b-d, f-h, j-l).

To further validate our striking observation, we evaluated general chromatin organisation by analysing chromatin accessibility by ATAC-Seq^28,29^ in all the clonal systems herein presented (HA1E: 10E, A7A, A9H, B2F and B5F; HA1ER: F12, 1F8, 1C9, 2A6 and 1D12; HepG2: 1A, 32G, 14F, 48E and 44D). Notably, we did not detect substantial changes in chromatin accessibility between any of the clones belonging to the same cellular system (HA1E: 178 598 peaks detected, HA1ER: 207 566 peaks detected and HepG2: 232 711 peaks detected. Fig. 3b, Supplementary Fig. 11) thus supporting our hypothesis that transcriptome state dynamics are not primarily controlled at the transcriptional level.

These data, obtained from three independent clonal cellular systems, suggest that steady-state RNA levels and hence metastable gene expression states are not established/maintained by diverging active transcription programmes but rather by a molecular machinery that operates at the post-transcriptional level. Furthermore, our data from multiple systems supports the hypothesis that this phenomenon, rather than an oddity, is a general biological feature.

### Altered transcriptomic states and phenotypic divergence between clones are not defined by major changes at the (phospho-) proteome level

The fact that transcriptomes diverge between cell lineages and that the observed divergence is accompanied by profound changes in phenotypic output raises the question whether major contributors to phenotype such as protein composition (e.g. translation) and/or signalling pathway status (e.g. MAPK phosphorylation) also diverge. To test these hypotheses, we first analysed the total protein content of clones 1F8 and 1C9 from the HA1ER system by label-free mass-spectrometry (LC-MS/MS) and performed a pairwise comparison between both phenotype-diverging subclones. Notably, from a pool of 3809 proteins analysed, only a small fraction was differentially expressed (FDR<0.01, FC>1.5, Fig. 4a) with 23 upregulated and 34 downregulated proteins, suggesting that, even though diverging clonal populations differ notably in their transcriptomes, they do not display a concordant difference in the proteome. Moreover, gene ontology analysis of the differentially represented proteins did not reveal enrichment in any biological pathways (Reactome v3.7^30^, Supplementary Fig. 12a) that could guide further hypotheses to explain the observed phenotypic output variability in, for example, such a complex phenotype as anchorage-independent growth. Therefore, these seemingly negative data prompted us to explore whether the diverging phenotypes could be explained by an imbalance in signalling pathway output and therefore we performed a phospho-proteomic analysis (TMT^31^) in the 1F8 and 1C9 populations (FDR<0.05, FC>0.4, Fig. 4c). Surprisingly, the obtained data including 305 detected phospho-sites across 141 proteins revealed only a minimal change in the phospho-proteome between both clonal populations with 33 phospho-sites displaying enhanced phosphorylation and 24 displaying reduced phosphorylation levels across only 28 proteins at bulk levels (Fig. 4c). Taken together, the remarkably little variance in the (phospho-) proteome between two subclonal populations which exhibit notable divergence in various phenotypic outputs encouraged us to look deeper into the data to elucidate the molecular mechanisms underlying this non-genetic phenotypic heterogeneity.

**Fig. 4.**
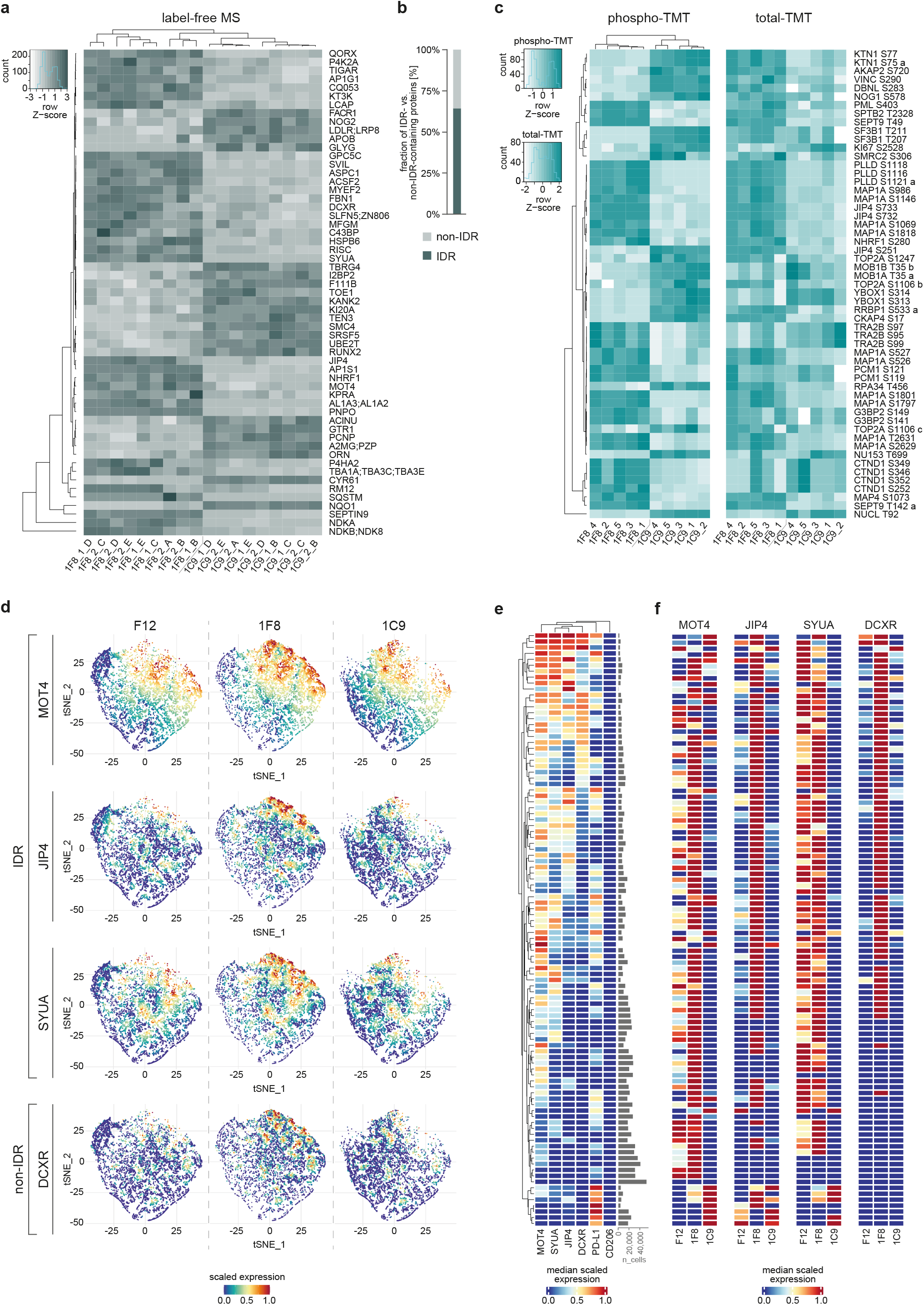
IDR-containing proteins show clonal divergence. **a**. Clones 1F8 and 1C9 from the HA1ER system were subjected to label-free LC-MS/MS. Heatmaps depict the identified differentially expressed proteins (FDR<0.01; FC>1.5). Data was pooled from 2 independent experiments containing 4 and 5 biological replicates each. Scale bar is depicted. **b**. Differentially expressed proteins identified in A were annotated considering their intrinsically disordered region (IDR) potential (AlphaFold2^16,17^). Histogram represents the fraction of IDR vs non-IDR proteins identified as differentially expressed. **c**. Clones from a were analysed by TMT-based phospho-proteomics. Heatmap (left) represents differentially represented phospho-sites between 1F8 and 1C9 (FDR<0.05; FC>0.4). Data was obtained from 5 biological replicates. Accompanying heatmap (right) displays total protein levels for differentially phosphorylated proteins identified. Scale bars are depicted. Letters at the end of a phosphorylation site represent the detection of a different peptide. **d**. A subset of differentially expressed IDR-containing proteins (1F8 vs 1C9) were analysed at a single-cell level by mass cytometry (CyTOF). Levels of MOT4, JIP4 and SYUA (all IDR-containing) were evaluated using specific antibodies in F12, 1F8 and 1C9. DCXR is also shown as a non-IDR diverging protein control. Scale bar is depicted. **e**. Heatmap represents the clustered expression of the antibody panel (MOT4, JIP4, SYUA and DCXR. PD-L1 and CD206 are included as a positive and negative control respectively) in collective data from all clones. The fraction of cells analysed in each cluster is depicted. Scale bar is depicted. **f**. Heatmaps represent the individual analysis for each antibody in the CyTOF panel (MOT4, JIP4, SYUA and DCXR) in each clone (F12, 1F8 and 1C9). Clustered expression levels are depicted. Scale bar is depicted.

### Clonally diverging lncRNAs localise and converge into perinuclear structures

To advance our understanding of the molecular mechanisms underlying phenotypic divergence in clonal cell populations, we endeavoured into annotating our various high-throughput datasets including SLAM-Seq, label-free mass spectrometry and phospho-proteomic data based on their molecular features. Intriguingly, a large fraction of the differentially expressed transcripts found by comparing 1F8 and 1C9 SLAM-Seq data were lncRNAs with approximately 30% of them corresponding to this category (FC>2. Fig. 3c, d). Given the ascribed regulatory role of lncRNAs in various cellular processes^32^, we evaluated the lncRNA content in our scRNA-Seq datasets where we validated our initial observations even across all three clonal systems in which the lncRNA content ranged from ∼30-64% between various conditions including clonal divergence and TGFβ-induced EMT (HA1ER, HA1E and HepG2. DE FC>3 for each pair-wise comparison included. Fig. 3e, Supplementary Fig. 9c).

Given the generality of our observations across the evaluated clonal lineages from the 3 different cell types, we combined data obtained from the SLAM-Seq assays with the scRNA-Seq datasets to build a composite lncRNA catalogue from each clonal system that could be experimentally queried. To do so, a subset of clone-specific lncRNAs were first evaluated by reverse transcription followed by qPCR (RT-qPCR), then by subcellular fractionation followed by RT-qPCR and finally by RNA-FISH followed by super-resolution microscopy. Notably, from the composite subset (SLAM-Seq + scRNA-Seq) of lncRNA candidates obtained from the HA1ER system, we were able to validate ∼45% (RT-qPCR) of the differentially expressed lncRNAs tested that diverged between 1C9 and 1F8 (Fig. 3f, Supplementary Fig. 9d). Moreover, from the HA1E and the HepG2 clonal systems, we validated ∼65% of the diverging lncRNAs by the same means (Supplementary Fig. 9e, f). Furthermore, we found that most of the lncRNA transcripts validated as divergent among states/clones display cytoplasmic localisation (subcellular fractionation followed by RT-qPCR) in both HA1ER and HepG2 clones (Fig. 3g, Supplementary Fig. 9g). Following up on these data, we moved forward to assess the intracellular localisation of these cytoplasmic clonally diverging transcripts by RNA-FISH (Zeiss Airyscan) and verified that all the evaluated lncRNA transcripts from the HA1ER and HepG2 systems indeed localised to the cytoplasm and more precisely, into perinuclear structures reminiscent of molecular condensates (Fig. 5, Supplementary Fig. 12a, b, RNA-FISH specificity control in Supplementary Fig. 12c).

**Fig. 5.**
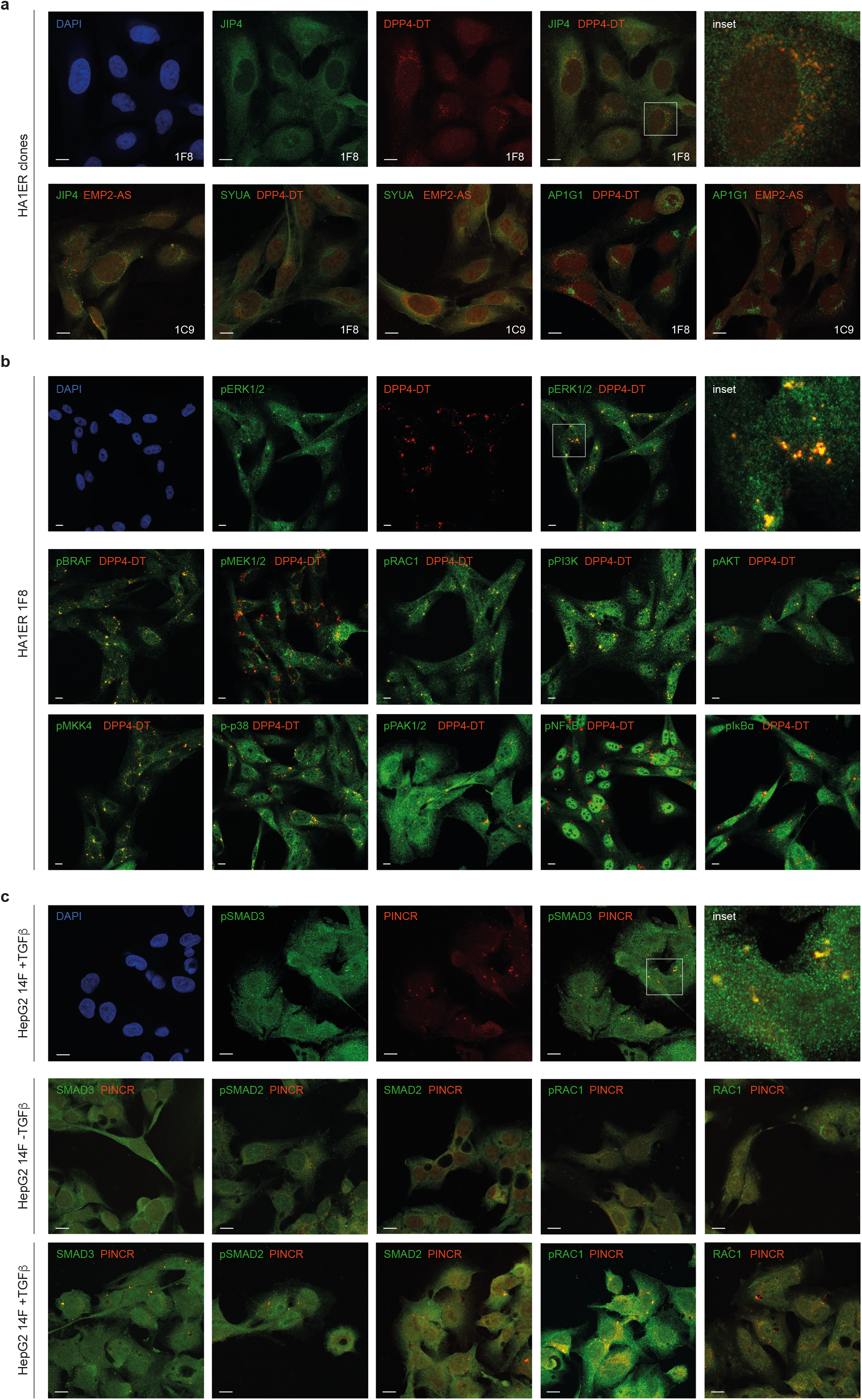
Cytoplasmic lncRNAs co-localize with IDR-containing proteins and activated signalling pathways. **a**. HA1ER clones 1F8 and 1C9 were subjected to immunocytochemistry for JIP4, SYUA and AP1G1 coupled to RNA-FISH targeting DPP4-DT or EMP2-AS lncRNAs and analysed by Airyscan microscopy. Upper panels depict individual channels (DAPI, JIP4 and DPP4-DT) and a merged image (JIP4/DPP4-DT) whilst lower panels depict overlay images for JIP4/EMP2-AS, SYUA/DPP4-DT, SYUA/EMP2-AS, AP1G1/DPP4-DT and AP1G1/EMP2-AS. Inset is displayed for the upper panel. Scale bar 10 μm. **b**. HA1ER clone 1F8 was subjected to immunocytochemistry for activated signalling pathways coupled to RNA-FISH targeting DPP4-DT lncRNA. Upper panels depict individual channels (DAPI, pERK1/2 and DPP4-DT) and a merged image (pERK1/2/DPP4-DT) whilst the lower panels depict merged images for DPP4-DT/pBRAF, DPP4-DT/pMEK1/2, DPP4-DT/pRAC1, DPP4-DT/pPI3K, DPP4-DT/pAKT, DPP4-DT/pMKK4, DPP4-DT/p-p38, DPP4-DT/pPAK1/2, DPP4-DT/pNFκB and DPP4-DT/pIκBα. Inset is displayed for the upper panel. Scale bar 10 μm. **c**. HepG2 clone 14F was either treated with TGFβ or with vehicle (+TGFβ or -TGFβ respectively) and subjected to immunocytochemistry for either active (pSMAD2, pSMAD3 and pRAC1) or their corresponding non-active (SMAD2, SMAD3 and RAC1) TGFβ-dependent signalling proteins coupled to RNA-FISH targeting PINCR lncRNA. Upper panel depicts individual channels (DAPI, pSMAD3 and PINCR) and an overlaid image (pSMAD3/PINCR) whilst the lower panels display SMAD3/PINCR, pSMAD2/PINCR, SMAD2/PINCR, pRAC1/PINCR and RAC1/PINCR. Inset is displayed for the upper panel. Scale bar 10 μm.

Given that our observations suggest that a large fraction of clonally diverging lncRNAs may modulate phenotypic output in a non-nuclear-linked manner and that most of the differentially expressed coding RNAs are not accompanied by increased/reduced protein levels, we hypothesised that most of the identified divergent RNAs (coding and non-coding) act as molecular scaffolds.

### Intrinsically disordered regions (IDR)-containing proteins clonally diverge and localize with peri-nuclear lncRNAs

Our mass spectrometry data (LC-MS/MS and Phospho-proteomics) demonstrated that only a small subset of proteins was differentially represented between 1F8 and 1C9 clones, suggesting that phenotypic divergence at the protein level may rely on discrete features such as intracellular localisation and scaffolding properties, among others. At the same time, we have shown that the cytoplasmic accumulation of divergent lncRNAs resembles the deposition of molecular condensates, a phenomenon for which a key role for lncRNAs has been previously described in the nucleus^33-37^. Therefore, as IDR-proteins have been shown to act as scaffolding molecules contributing to the formation of biomolecular condensates^38-40^, we hypothesised that these subsets of proteins could play a role in the nucleation/localisation of the identified perinuclear lncRNAs. To evaluate this possibility, we moved on to validate a subset of the identified differentially expressed proteins between clones 1F8 and 1C9 from the HA1ER system prioritising the annotation of IDR-containing proteins, to which the majority of the diverging proteins could be ascribed to (∼63%, Fig. 4b, AlphaFold2 annotation^41,42^).

To initially validate the observed differential protein expression between HA1ER clones 1F8 and 1C9, we set up a CyTOF analysis panel for a subset of differentially expressed proteins (IDR-proteins JNK-interacting protein 4 (JIP4), *α*-Synuclein (SYUA), and monocarboxylate transporter 4 (MOT4), myelin expression factor 2 (MYEF2), heat shock protein beta-6 (HSPB6), sequestosome-1 (SQSTM) and non-IDR-protein L-xylulose reductase (DCXR)) to determine the levels of each protein in the HA1ER founder clone F12 and in 1F8 and 1C9 subclones in a comparative manner. Notably, all the analysed proteins labelled subpopulations within the F12 clone (Supplementary Fig. 13d, e) suggesting that clonal divergence could also be traced at the proteome level even when the differentially expressed proteins are only a minimal fraction of the total proteome. Next, we assessed a subset of these proteins (JIP4, SYUA, MOT4, DCXR, PD-L1 (positive control) and CD206 (negative control)) for their divergent expression in subclones 1F8 and 1C9 together with their founder clone F12. Whilst none of the included proteins displayed cell cycle-dependent expression (Supplementary Fig. 13f), clone 1F8 contained a major cellular subpopulation where candidate antibody signal (JIP4, SYUA, MOT4 and DCXR) is predominant and that was nearly absent in clone 1C9 (Fig. 4d-f, Supplementary Fig. 13b, c).

Given that a large fraction of the clonally diverging proteins belongs to the IDR-containing family and could potentially act as molecular scaffolds, we next expanded our candidate list to accommodate an additional diverging IDR-protein, gamma1-adaptin (AP1G1), and moved forward to evaluate whether the diverging IDR-proteins and the observed lncRNAs enriched in the cytoplasm could indeed co-localise within the observed perinuclear condensates. Therefore, we performed RNA-FISH coupled to immunocytochemistry (ICC) for a subset of diverging lncRNAs and IDR-proteins and analysed their intracellular distribution by super-resolution microscopy (Zeiss Airyscan). Interestingly, we could show that in clonal lineages of HA1ER cells diverging lncRNAs and IDR-proteins co-localise in the perinuclear region (lncRNAs: DPP4-DT (clone 1F8) and EMP2-AS (clone 1C9). IDR-proteins: JIP4, SYUA), and only for IDR-protein AP1G1 could no such co-localisation be detected with either lncRNA (Fig. 5a, Supplementary Fig. 12a, RNA-FISH specificity control shown in Supplementary Fig. 12c). Moreover, we also validated the co-localisation of IDR-protein JIP4 and several diverging lncRNAs (PINCR and LINC00494 (clone 14F), SCAT8 and LINC2476-203 (clone 44D)) in the perinuclear region in the HepG2 clonal model (Supplementary Fig. 12b), for which the observations were in agreement with data obtained from the HA1ER system.

### The Signal Integration Portal (SIP)

So far, our findings have revealed that a subset of diverging lncRNAs and a subset of IDR-containing proteins co-localise into perinuclear cytoplasmic structures in a variety of clonally diverging cellular models. However, none of our findings indicated a functional/effector link that could explain the observed divergence in phenotypic output for a given complex cellular behaviour such as the capability to grow in anchorage-independent conditions, migration or fractional killing. Therefore, we hypothesised that cell-to-cell variability in phenotypic output relies on the re-arrangement of signalling pathways following different configurations.

To further explore this concept, we contemplated the possibility that the peri-nuclear structures that we have identified could nucleate signalling components that would alter phenotypic output. Given that we have merely detected 28 statistically significant differentially phosphorylated proteins between our flagship HA1ER clones 1F8 and 1C9, and for which phospho-site specific antibodies are not available, we decided to test whether multiple activated signalling pathways would converge within the observed peri-nuclear structures. To do so, we built a panel of phospho-site specific antibodies directed towards multiple major signalling pathways including RAF/MEK/ERK, PI3K/AKT, IκBα/NFκB and MKK4/p38, and performed RNA-FISH (targeting peri-nuclear localised diverging lncRNA DPP4-DT) coupled to ICC (phospho-site specific; Supplementary Table) in clone 1F8. Strikingly, after super-resolution imaging analysis, we observed that multiple active signalling pathways (effector proteins or phospho-proteins) were indeed localised in the peri-nuclear structures delineated by the lncRNA DPP4-DT (pARAF, pBRAF, pMEK1/2, pERK1/2, pRAC1, pMKK4, p-p38, pPI3K, pAKT, pPAK1/2, pIκBα and pNFκB. Fig. 5b, Supplementary Fig. 14a). Intriguingly, rather unrelated active signalling pathways coalesce into these cytoplasmic condensates suggesting that the observed structures could function as a signal organiser to determine phenotypic output and thus we termed this compartment the Signal Integration Portal (SIP).

To further verify the existence of the SIP and taking advantage of the capability of the HepG2 model to undergo TGFβ-induced EMT, we decided to evaluate if members of the TGFβ signalling pathway would converge into SIP structures delineated by HepG2-specific diverging lncRNA PINCR in clone 14F by RNA-FISH coupled to ICC. To do so, we included members of the canonical TGFβ pathway (SMAD2 and SMAD3) in addition to members of the non-canonical pathway also reported to become active by the presence of TGFβ (RAC1 and ROCK1) and evaluated their total levels and those of their corresponding phosphorylated active forms. As expected for TGFβ-naïve conditions, PINCR localised to the peri-nuclear region in a punctuated pattern across most of the cells (Fig. 5c, Supplementary Fig. 14b) whilst most of the components of the TGFβ signalling pathway and their phosphorylated counterparts displayed a diffuse cytoplasmic localisation (Fig. 5c, Supplementary Fig. 14b). However, upon TGFβ treatment, PINCR (proxy for SIP particles) co-localised with active members of the TGFβ signalling cascade including pSMAD2, pSMAD3 and pRAC1 (Fig. 5c, Supplementary Fig. 14b) suggesting that signalling pathway components are brought together to the SIP to integrate biological cues and fine-tune phenotypic output. Interestingly, whilst ROCK1 was observed within SIP condensates, we did not detect pROCK1 (Tyr913) suggesting that either this phospho-protein is not recruited to the SIP or the epitope is masked within SIP-condensates (Supplementary Fig. 14b).

Altogether, we showed that in phenotypically diverging clonal populations of fully committed and differentiated cellular models (HA1ER and HepG2) lncRNAs, IDR-containing proteins and active signalling effectors from multiple signalling cascades coalesce into a peri-nuclear structure which we termed Signal Integration Portal (SIP) that sits at the crossroads of phenotypic divergence nucleating inputs from the nucleus (lncRNA), the cytoplasm (IDR-proteins and signalling cascades) and the environment (signalling cascades).

### Remodelling cell states by altering the SIP lncRNA composition

We have already shown that diverging molecules from different families (lncRNAs, IDR-proteins, signalling proteins) converge into peri-nuclear SIP structures which may be involved in the establishment/propagation of transcriptome states and in dictating phenotypic output. Therefore, to further our claims, we hypothesised that altering the SIP composition may result in transcriptome re-arrangements. To do so, we designed antisense oligonucleotides (G, gapmers) targeting a subset of SIP-associated lncRNAs identified in the diverging HA1ER clones 1F8 and 1C9 (DPP4-DT, AL161733.1 and AC008464.1) and in HepG2 clones 14F and 44D (PINCR and SCAT8). Importantly, to strengthen our claims, we designed our assays to include a non-targeting control gapmer and two gapmers targeting different regions per target transcript (when possible) to evaluate knockdown effects in a confident manner. After verifying successful knock-down of the evaluated lncRNAs from either system by RT-qPCR (Fig. 6e, Supplementary Fig. 15c), cells from each experimental condition were processed for scRNA-Seq (10x Genomics Chromium 3’). After sequencing, obtained data was pre-processed to mitigate clustering artefacts due to cell cycle progression and abnormal mitochondrial counts and clustered using Seurat. Strikingly, upon knock-down of the assessed lncRNAs we observed transcriptome re-arrangements relative to the non-targeting control (ctrl) that ranged from the loss of a single transcriptome state whilst largely preserving transcriptome heterogeneity (PINCR-G: cluster 8; HepG2. Supplementary Fig. 15b, g), the reduced representation of certain states and the emergence of novel ones whilst preserving a certain degree of the original transcriptome states (AL161733.1: cluster 10 reduced, clusters 6 and 9 novel for G1, clusters 0, 11 and 12 novel for G2; HA1ER. SCAT8: cluster 8 reduced, cluster 10 novel for G1, cluster 5 novel/increased for G2; HepG2. Fig. 6a, Supplementary Fig. 15b, f, g) to a complete reprogramming of the metastable transcriptomic state landscape (AC008464.1-G and DPP4-DT-G1 and -G2; HA1ER. Fig. 6a, Supplementary Fig. 15a, f). Notably, while both gapmers targeting the same lncRNA (DPP4-DT and AL161733.1) led to distinct transcriptome state re-arrangements which might partly be due to differences in knockdown efficiency, we detected specific transcriptomic markers concordantly upregulated by both gapmers, indicating common intracellular effects (Fig. 6e, Supplementary Fig. 15d). These findings suggest that SIP-bound lncRNA levels harbour the capability to remodel the transcriptome landscape and hence modulate phenotypic output.

**Fig. 6.**
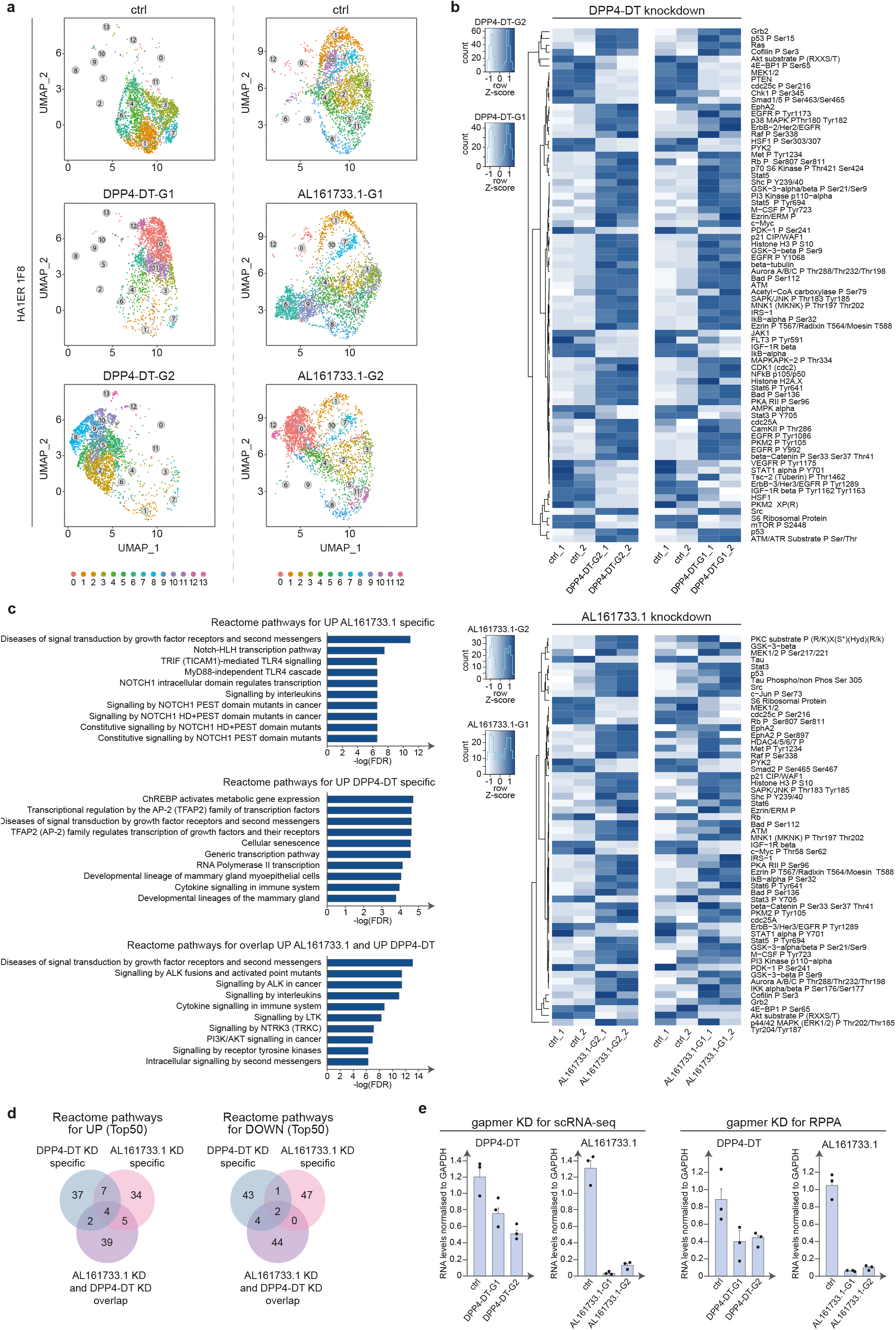
SIP destabilisation promotes transcriptome and signalling pathway re-organisation. **a**. HA1ER clone 1F8 was subjected to antisense mediated knock-down (Gapmer) for DPP4-DT or AL161733.1 lncRNAs and processed for scRNA-Seq after 48h (24h for DPP4-DT-G2). UMAP plots depict transcriptome states in control (non-targeting control gapmer) and upon knockdown (DPP4-DT-G1, DPP4-DT-G2, AL161733.1-G1 and AL161733.1-G2). Two alternative antisense molecules were used targeting different regions of the target transcript (G1 and G2). Individual states are depicted in colour and numbered. **b**. HA1ER clone 1F8 was subjected to antisense mediated knock-down (Gapmer) for DPP4-DT or AL161733.1 lncRNAs and further processed for Reverse Phase Protein Arrays (RPPA) after 48h (24h for DPP4-DT-G2) to evaluate signalling integrity. Two alternative antisense molecules were used targeting different regions of the target transcript (G1 and G2). Heatmaps display differentially represented signalling pathway members (≥20% increase/decrease to control) upon either DPP4-DT or AL161733.1 knockdown. Two biological replicates were processed for each condition. Scale bars are depicted **c**. Pathway analysis (Reactome) on HA1ER clone 1F8 upon DPP4-DT or AL161733.1 lncRNAs knockdown (FDR<0.01). Top 10 upregulated pathways are shown for antibody signals specifically upregulated upon DPP4-DT (DPP4-DT-specific) or AL161733.1 knock-down (AL161733.1-specific), as well as for upregulated signalling pathway members detected in both knock-downs (overlap). **d**. Venn diagrams depict the intersection between the Top 50 enriched terms (Reactome, Up- or downregulated) from DPP4-DT and AL161733.1 lncRNAs knockdown datasets for specific or overlapping signalling pathway members identified. **e**. lncRNA knockdown was evaluated for cells processed for scRNA-Seq shown in **a** (left) or for cells subjected to RPPA shown in **b** (right). Transcript levels of lncRNAs DPP4-DT and AL161733.1 were determined by reverse transcription followed by qPCR (RT-qPCR) and were normalised to GAPDH. Histograms represent the mean ± propagated error of 3 technical replicates. Individual data points are depicted.

### Cell state remodelling is accompanied by profound changes in signalling

To further explore the phenotypic consequences of SIP disruption and transcriptome remodelling in the HA1ER model, we evaluated whether major signalling pathways would be altered upon a given lncRNA knockdown. To do so, we took advantage of a pathway-specific antibody panel (180 antibodies; Supplementary Table) and evaluated signalling pathway remodelling by Reverse Phase Protein Array (RPPA^43^) in the HA1ER clone 1F8 (increased expression of lncRNAs AL161733.1 and DPP4-DT when compared to 1C9 clone). Upon lncRNA knockdown (∼50-94% KD when compared to the non-targeting control, Fig. 6e), we first validated transcriptome remodelling by evaluating a subset of identified expression markers that become upregulated under these conditions by RT-qPCR (MDM2 and CDKN1A. Supplementary Fig. 15d, e) and then proceeded to run the RPPA assay.

Strikingly, upon lncRNA AL161733.1 knockdown, from the 155 antibodies for which we observed a robust signal, we found 41 upregulated signals whilst 16 were downregulated with any of the antisense gapmers when contrasted to the non-targeting control (≥20% increase/decrease to control) and which showed the same trend in both gapmers (Fig. 6b bottom). Moreover, when knock-down of lncRNA DPP4-DT was evaluated, 50 signals were upregulated whilst 25 were downregulated with any of the gapmers with the same trend for both gapmers (Fig. 6b top, ≥20% increase/decrease to control). In ∼90 % of identified up- or down regulated targets both gapmers showed the same trend, indicating that similar intracellular effects at the signalling pathway level are triggered by both gapmers targeting the same lncRNA.

Notably, as we hypothesised that the lncRNAs evaluated are involved in the integration of signalling information by nucleating pathways, we annotated each of the assessed antibodies with their respective pathways (Reactome, Top50 Terms, FDR<0.01). For this, we extracted the upregulated and downregulated antibody signals with specific changes for lncRNA AL161733.1 and lncRNA DPP4-DT knockdowns, as well as the list with overlapping changes upon knock-down of lncRNA AL161733.1 and lncRNA DPP4-DT (Fig. 6c, d, Supplementary Fig. 15h). Outstandingly, separate knock-down of both lncRNAs led to a modulation of common pathways (overlap), such as upregulation of signalling by ALK in cancer, PI3K/AKT signalling and signalling by LTK, whereas the AP-2 (TFAP2) family of transcriptional regulators, several developmental lineages (mammary gland, integumentary system) and signalling by KIT in disease, amongst others, were downregulated (Fig. 6c, Supplementary Fig. 15h), suggesting that both lncRNAs contribute to a general mechanism of signalling pathway control. However, knocking down lncRNA AL161733.1 also resulted in the modulation of specific pathways such as the upregulation of NOTCH signalling and downregulation of signalling by TGFβ family members and signalling by WNT (Fig. 6c, d, Supplementary Fig. 15h), whilst knocking down lncRNA DPP4-DT resulted in the modulation of a completely different subset of pathways, including the deregulation of multiple transcription-regulating pathways (Fig. 6c, d, Supplementary Fig. 15h), thus suggesting the existence of an exquisite mechanism of signalling pathway control where individual lncRNAs modulate a defined set of pathways in a specific manner.

Finally, the observation that major – unrelated – signalling pathways were indeed altered upon lncRNA knockdown suggests that by solely adjusting SIP composition through the reduction of SIP-linked lncRNA levels, both transcriptome re-arrangement and, as hypothesised, major remodelling of signalling pathways occurs.

### Cell state remodelling is accompanied by signalling pathway reorganisation/redistribution and altered phenotypic output

Our data suggest that SIP-bound lncRNAs are major contributors to phenotypic output as altering SIP composition using lncRNA knockdown results in signalling pathway rewiring. Therefore, if SIP complexity is at the crossroad of transcriptome heterogeneity, signalling organisation/integration and phenotypic output, altering SIP composition should result in profound changes in phenotypic output. Therefore, to evaluate this hypothesis, we performed a series of phenotypic assays where we took advantage of the efficient gapmer knockdown and its observed consequences on transcriptome rearrangement and signalling pathway rewiring (Fig. 6, Supplementary Fig. 15).

First, given the profound changes in signalling pathway organisation upon DPP4-DT knockdown in the 1F8 clone from the HA1ER system, we set out to evaluate whether signalling pathway components would relocalise within the cell under these conditions and performed RNA-FISH followed by ICC for a subset of SIP-associated signalling proteins (pRAC1, pPAK1, pMKK4 and pMEK1/2). Strikingly, upon DPP4-DT knockdown in the HA1ER 1F8 clone, we could evidence redistribution of pRAC1 from SIP condensates to the nuclei, pPAK1 accumulation in the cytoplasm, a pronounced build-up of pMKK4 in the perinuclear area accompanied by the loss of distinct punctate structures and the coalescence of pMEK1/2 into a different kind of perinuclear accumulation reminiscent of biomolecular condensates (Fig. 7a, Supplementary Fig. 16) thus supporting the notion that SIP-condensates contribute to signal integration by acting as a signal organising hub. Encouraged by these results, we further set out to evaluate if SIP-linked lncRNA knockdown would affect, what we believe to be one of the most extreme phenotypes under SIP influence, transformation potential. Therefore, we knocked down lncRNA AC008464.1 in clone 1C9 of the HA1ER system (Fig. 7c), which displayed increased clonogenicity when compared to its parental clone F12 or sibling clone 1F8 as established in previous soft agar assays (Fig. 2d, Supplementary Fig. 7c). Obtained data revealed that altering SIP-linked lncRNA AC008464.1 levels reduced clonogenic potential of clone 1C9 to ∼53% when compared to non-targeting control gapmer treated cells (Fig. 7c) suggesting that the clonogenic potential of clone 1C9 is severely impaired when SIP composition is compromised.

**Fig. 7.**
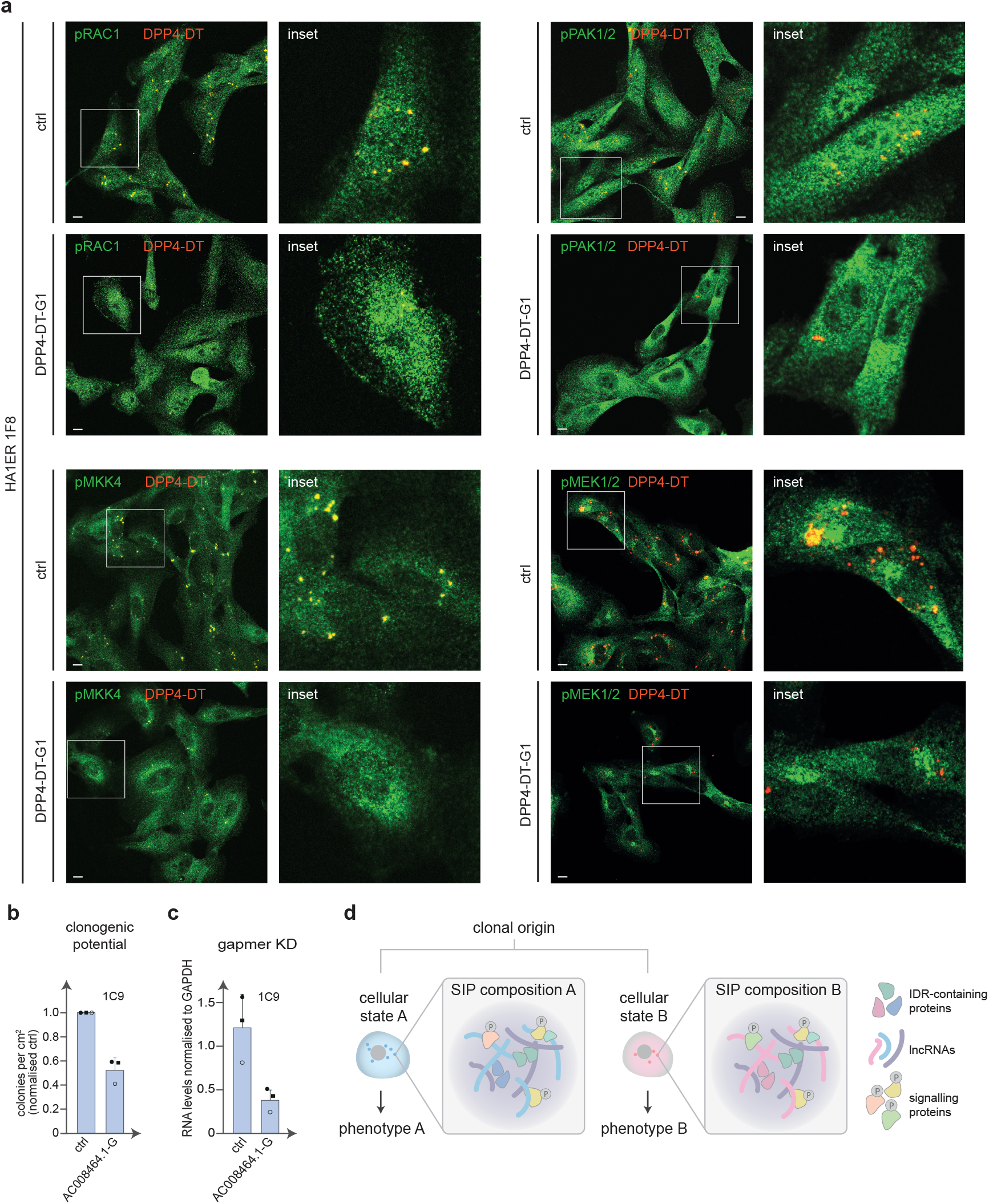
SIP disruption results in intracellular relocalisation of signalling pathways and alters phenotypic output. **a**. HA1ER clone 1F8 was subjected to antisense-mediated knock-down for DPP4-DT lncRNA (DPP4-DT-G1) or non-targeting control gapmer (ctrl) and further processed for immunocytochemistry for activated signalling pathways (pRAC1, pPAK1/2, pMKK4 and pMEK1/2) coupled to RNA-FISH targeting lncRNA DPP4-DT lncRNA. Panels depict overlay images of signalling protein (green) and DPP4-DT (red) and are accompanied by an inset. Bar 10 μm. **b**. Cells from HA1ER clone 1C9 were subjected to antisense-mediated knock-down (gapmer) targeting lncRNA AC008464.1 or non-targeting control (ctrl) and further processed for clonogenic soft agar assay. Clonogenic potential is presented as the number of colonies per cm^2^ upon knock-down normalised to the control. Histograms represent the mean value ± standard deviation (SD) of 3 independent experiments performed in 4-6 replicates. Individual data points are shown. **c**. Knock-down of lncRNA AC008464.1 prior to clonogenic assay shown in **b** was evaluated by reverse transcription followed by qPCR (RT-qPCR) normalised to GAPDH. Histograms represent the mean ± SD of 3 independent experiments. Individual data points are depicted. **d**. Toy diagram depicting the link between cellular state, SIP composition and phenotype. Identified components are depicted on the right.

Taken together, we propose the Signal Integration Portal (SIP) as a peri-nuclear molecular entity where biological information from various non-genetic compartments including cytoplasmic lncRNA content, intrinsically disordered region-containing proteins and active signalling pathways converge (Fig. 7d) to modulate phenotypic output, thus controlling cell plasticity and the degree of non-genetic heterogeneity for a given cellular system.

## Discussion

Cell plasticity, an inherent characteristic of all known organisms, is one of the most widespread phenomena observed in biology. However, though a myriad of hypotheses have been brought forward, very little is known about the molecular mechanisms underlying this phenomenon.

Paradoxically, years of research centred around the genetic compartment could not explain the plastic behaviour displayed by cells and organisms overall. However, a multitude of examples challenging the role of the genome as a master controller underlying this phenomenon have come to light including bet-hedging in bacteria^44,45^, protozoan development/evolution^46-49^ and the emergence of resistance to anti-cancer therapy^50-54^. Remarkably, the advent of multimodal single-cell technologies opened a new dimension of biological insight that fuelled the in-depth exploration of the non-genetic compartment and has enabled the study of the molecular determinants underlying cell plasticity^8,55-60^. In that regard, we and others ventured into exploring transcriptome content as a determinant of cell plasticity in mammalian cellular systems and found that indeed steady-state transcriptome heterogeneity harbours sufficient biological information to drive divergent phenotypic output even in fully controlled clonal cellular systems, thus fuelling molecular and functional heterogeneity within populations^8,47,61^. Strikingly, in all the models that we have tested so far, the contribution of active transcription to transcriptome heterogeneity is negligible and does not support a model where transcriptome divergence is instructed by changes in gene expression programmes (as defined by alterations in chromatin structure resulting in different subsets of genes being actively transcribed). Instead, transcriptome plasticity seems to rely on post-transcriptional regulatory modules wherein a subset of cytoplasmic lncRNAs and intrinsically disordered region-containing proteins (IDR-proteins) play a key role throughout phenotypic divergence and are enriched in subsets of cellular states. Both observations challenge the status quo as, to begin with, lncRNA function has been ascribed mainly to the nucleus where they, in principle, would contribute to genome control (active transcription, chromatin architecture, among others)^32,62^ whilst our data supports a key role for lncRNAs within the cytoplasm. Consequently, our data does not support a direct role for lncRNAs in genome control as the primary driver of transcriptome divergence, whilst highlighting marked differences between nascent and steady-state RNA pools. The latter observation comes as no surprise as it has been acknowledged for decades that the contribution of active transcription to steady-state RNA levels is rather modest and that major players in the regulation of transcriptome content operate at a post-transcriptional level^63-66^.

An important aspect of our study stems from the fact that to explore non-genetic divergence and evolution we required - first and foremost - to characterise the level of genetic divergence in each cellular population to ultimately derive genetically stable cellular models that would allow us to evaluate the non-genetic compartment avoiding genetic variation confounding factors. In that context, HA1E and HA1ER cell systems were an ideal choice as previous reports and our own observations confirmed that these cellular models accumulate over time negligible genetic alterations^67^. Together with our HepG2 clonal system, for which genetic stability has also been verified, our catalogue of isogenic cell systems comprises non-transformed (HA1E) and transformed cells originating from distant tissue-of-origin (HA1ER, kidney and HepG2, liver) and therefore displaying pronounced differences in their non-genetic state repertoire. Our coherent observations in unrelated cellular systems suggest that our findings can be juxtaposed to any cellular system and are a biological general feature rather than an odd occurrence.

Following this line of thought, we argue that cells thriving in vivo could be constantly rewiring their non-genetic landscape to accommodate their individual needs and/or the needs of the tissue and that cell plasticity is a pervasive feature across organismal biology. It follows that cell plasticity as defined in our report, may be wired to adapt to a myriad of both local and systemic signals acting on the tissue of origin (e.g. differentiation signals, immune challenge, therapeutic pressure) and respond in turn by expressing a variety of phenotypes to increase population fitness and thus securing their perpetuation. Importantly, our findings corroborate the concept that cell plasticity may underlie a fundamental constrain for therapeutic development as cell plasticity is ever-present and often overlooked in bulk analyses, resulting in a large panoply of phenotypic outputs often obscuring data interpretation. For instance, whilst some of the cellular states observed in a given cellular system or tissue could be therapy susceptible, the possibility to find intrinsic resistant states already present within the cellular population is almost a certainty. Moreover, as many state-of-the-art anti- cancer therapies are based on genotoxic attack, the risk associated with generating multiple levels of cell plasticity – a palette of metastable states associated with each drug-induced genotype – complicates the outlook further.

Strikingly, our data supports a model where cell plasticity is not infinite but defined by a limited number of predominant states that are characterised by state-specific SIP condensates, which prompts the question about how metastable states are inherited and propagated throughout the progeny. An obvious model would immediately consider the asymmetric inheritance of (SIP-) components as a centre piece underlying cell plasticity. In that regard, asymmetric cell division and asymmetry in the inheritance of RNA molecules and organelles has been widely documented as contributing to phenotypic output and stem-cell differentiation^68-71^, key mechanisms to keep in mind whilst delving further into SIP biology.

Interestingly, our model suggests that transcriptome organisation and divergence could result or be directed by a re-organisation of the molecular players involved in the process and is mainly controlled by their intracellular localisation. This exquisite molecular mechanism would ensure the generation of molecular and cellular heterogeneity by minimal energetic means and could provide the basis to maintain a basal level of phenotypic divergence within clonal populations. Moreover, it brings to light the observation that identity (i.e. cell type) and phenotypic output are not necessarily coupled and that cells, through the means of cell plasticity, can dramatically follow entirely different paths despite their identity. This concept opens a new dimension of biological insight as for instance in cancer research, most of the single cell analyses are based on cell type annotation (using hallmark markers) and not on what the cells are indeed doing in their context. Therefore, there is a clear need to further characterise and distinguish between cell identity and cell activity (expression of the phenotype) as it could completely alter the current view underlying how cellular systems operate.

Finally, further characterisation of SIP components and architecture, perhaps linked with a distinct set of phenotypes of interest, is expected to reveal suitable targets for SIP disruption or SIP reorganisation (i.e. lncRNAs, IDR-proteins, others) which in turn would allow the targeted fine-tuning of cellular states towards a desired phenotype (e.g. anti-cancer drug susceptible). However, it is our view based on observations made by us and others over the years^1,2,8,18,53,72-74^, that cell plasticity cannot be reduced/removed from a system, as plastic cell phenotypes and the dynamic intracellular molecular landscape impinges a fundamental biological property through which cell populations adapt when subjected to intra-/extracellular environmental changes. Therefore, the identification of molecular switches that would allow the controlled, targeted and non-reversible transition to a distinct – advantageous – phenotype might be a cumbersome task to achieve.

## Materials & Methods

### Cell lines

Immortalised HA1E (hTERT and SV40 ER) and tumorigenic HA1ER cells (hTERT, SV40ER and HRAS-G12V) from stepwise tumorigenesis models generated from normal human embryonic kidney cells were obtained from Prof. Weinberg (Broad Institute of MIT and Harvard, Cambridge, USA^19^). HepG2 cells were obtained from ATCC (ATCC, cat. HB-8065). HA1E and HA1ER cells were cultured in DMEM (1g/L glucose) (Gibco, cat. 21885025) supplemented with 10% heat-inactivated foetal calf serum (FCS, Gibco) and 100μg/ml hygromycin (hTERT selection, Thermo Fisher Scientific, cat. 10687010), 400μg/ml G418 (SV40-ER selection, Thermo Fisher Scientific, cat. 10131035), 0.5μg/ml puromycin (HRAS-G12V selection, Thermo Fisher Scientific, cat. A1113803) and 50 μg/ml gentamicin. HepG2 cells were cultured in DMEM high glucose (4.5g/L glucose) (Thermo Fisher, cat. 41966029) supplemented with 10% heat-inactivated FCS (Gibco) and 50U/ml Penicillin-Streptomycin (Thermo Fisher, cat. 15070063). Single cells were isolated by flow cytometric cell sorting (BD FACSAria III; BD Biosciences) and expanded in standard 2D culture conditions to establish clonal populations. LentiX 293T were obtained from Prof. Lacaud (Cancer Research UK Manchester Institute, Manchester, UK) and cultured in DMEM (1g/L glucose) (Gibco, cat. 21885025) supplemented with 10% heat-inactivated FCS (Gibco). HA1E and HA1ER cells were plated at a confluency of 1 × 10^6^ cells/6010mm^2^ surface with 10ml fresh medium, passed every 48⍰hours and kept in culture for a maximum of 15 passages. HepG2 cells were plated at a confluency of 1.5 × 10^6^ cells/6010mm^2^ surface with 10ml fresh medium, passed every 48⍰hours and kept in culture for a maximum of 15 passages. All cell lines were grown at 37°C with 5% carbon dioxide (CO_2_) saturation. Mycoplasma contamination was analysed by reverse transcription followed by quantitative Polymerase Chain Reaction (RT-qPCR) (VenorGeM qEP, cat. 11-9100).

### Lineage tracing libraries (LTv-BC-H2B-GFP)

The episomal vector GBX (EBNA1/OriP) was a gift from Linzhao Cheng (Addgene, cat. 64123) that was engineered to build a low-copy propagating vector in bacterial hosts (*bom and rop) an*d with the potential to express H2B-GFP fusion protein in mammalian systems (LTv-H2B-GFP). Using this template, Barcode decay Lineage Tracing (BdLT-Seq) plasmids were generated by introducing a molecular barcode into the 3’UTR of H2B-GFP sequence. Briefly, the 3’UTR of the LTv-H2B-GFP vector was PCR amplified using a primer that contains a 12 nucleotide long random sequence split into three segments by intervening nucleotides (NNNNGCGNNNNTGANNNN). Next, PCR amplicons were cloned into the LTv-H2B-GFP vectors using HindIII and XbaI restriction sites (LTv-BC-H2B-GFP vectors). The generated high-complexity library was propagated using electrocompetent MegaX DH10B T1 bacteria strain (Invitrogen, cat. C640003). The final LTv-BC-H2B-GFP lineage tracing episome library contained approximately 1.2 × 10^6^ unique barcodes and was already validated and used in our previous study^8^.

### Transfection with the LTv-BC-H2B-GFP episome libraries

Transfection of HA1ER cells with LTv-BC-H2B-GFP lineage tracing episome library was performed following standard direct transfection protocol using Lipofectamine LTX reagent (Invitrogen, cat. 153385000). The expression of H2B-GFP was verified by fluorescent microscopy (Thermo Fisher Scientific, cat. AMEX1000) and flow cytometry (BD LSRFortessa X-20 Cell Analyzer, BD Biosciences) 24- or 48-hours post-transfection depending on experimental set up (usually 25-35% of H2B-GFP positive cells). Flow cytometry data was analysed using FlowJo software (BD, version 10.8.0).

### BdLT-Seq

A detailed BdLT-Seq protocol can be found in our published protocol^75^. HA1ER cells (clones F12, 1F8 and 1C9) were transfected with LTv-BC-H2B-GFP episome library as described. Forty-eight hours post-transfection GFP positive cells were sorted using BD FACSAria III Cell Sorter (BD Biosciences) or BD InFlux Cell Sorter (BD Biosciences) aiming for >94% purity after sorting. Cells were expanded in standard culture conditions for five days – Phase I – a barcode decay phase in which most of the episomes are lost upon cell division. The second barcode decay phase – Phase II - begins approximately 5-7 days after transfection where the episomes stabilize however still displaying a characteristic episome loss. To perform BdLT-Seq (only during phase II), cells were trypsinised (Invitrogen, cat. 25300104), collected by centrifugation at 230 rcf, 20°C, and plated at 2,000 to 3,000 cells per well (0.32cm^2^) according to experimental conditions. Five to six days later (∼ 80,000 cells), cells from each clonal population were split into two samples: sample N1 (“Day 0”) was stored in liquid nitrogen (90% FCS, 10% DMSO) whereas sample N2 (“Day 4”) was expanded in standard culture conditions for four additional days. Four days later, cells were stored in liquid nitrogen - sample N2 (“Day 4”) (90% FCS, 10% DMSO). The day before single-cell RNA sequencing, cells were returned to standard 2D culture for 24 hours. The following day, samples were processed for scRNA-Seq sequencing (10x Genomics Chromium, Single-Cell RNA-Seq System, version 3.1, cat. PN-1000269) coupled to lineage tracing (BdLT-Seq). Briefly, cells were collected by trypsinisation, washed in 5% FCS PBS and resuspended in 5% FCS PBS. Hashtag antibodies were used to specifically label individual samples for multiplexing purposes. For that, cells were incubated with the hashtag antibodies coupled to a specific oligonucleotide (HTO) (BioLegend, cat. A0251-A0260) at a ratio of 0.5 μg per 50,000 cells (stock 10 μg/ml) for 30 min, on ice. Cells were then washed three times using 0.04% BSA PBS and counted using Neubauer counting chamber (Marienfeld, cat. 0640030). Samples were then combined according to the experimental setting and 20,000 cells per channel (for the expected outcome of 10,000 cells/channel) were loaded onto 3 channels of the 10x Chromium platform (10x Genomics, cat. 1000204). Samples were processed following standard 10x Chromium protocol (10x Genomics, version 3.1, cat. PN-1000269) up until cDNA amplification reaction. cDNA amplification reaction was supplemented with the HTO additive primer and followed by a cDNA clean-up separating the HTO fraction from the cell cDNA using standard methodologies (Beckman Coulter, AMPure XP, cat. A63880). All size selected fractions (cell cDNA and HTO cDNA) were assessed by Fragment Analyser 5200 (Agilent, cat. M5310AA). Next, HTO depleted cDNA fraction (cell cDNA) was used to capture H2B-GFP cDNA. For that, 5μl of Dynabeads MyOne Silane beads (Invitrogen, cat. 37002D) were washed twice in 500μl RLT lysis buffer (Qiagen, cat. 79216), resuspended in 140μl RLT buffer (3.5x volume of initial sample) and added to 40μl of the HTO depleted cDNA fraction. Then 180μl of absolute ethanol was added to the reaction (4.5x volume of initial sample), mixed well and incubated for 15 minutes at room temperature in a ThermoMixer C (300rpm, Eppendorf, cat. EP5382000015). The supernatant was discarded, and the beads were washed three times in 80% ethanol, dried for 3-4 minutes at room temperature, and eluted in 5μl nuclease free water (10-15 minutes at room temperature, ThermoMixer C at 300rpm). To capture H2B-GFP cDNA, a set of 4 sequence-specific biotinylated probes targeting H2B-GFP was used. The probes were hybridised H2B-GFP cDNA using xGen Hybridization and Wash Kit (IDT, cat. 1080577) following manufacturer’s instructions. Briefly, 8.5μl of Gen 2x Hybridization buffer was combined with 2.7μl of Gen Hybridization enhancer, 5μl cDNA fraction and 2μl of the probes (4 pmoles, pool of 4 probes), mixed, incubated for 10 minutes at room temperature, followed by the incubation in a thermocycler: 30 seconds at 95°C and 16 hours at 65°C (lid temperature 105°C). Following hybridisation, the H2B-GFP cDNA fractions were pulled down using streptavidin beads. Twenty microliters of streptavidin beads were washed twice using 100μl xGen 1x Bead wash buffer and then resuspended in 17μl bead resuspension buffer (8.5μl xGen 2x Hybridization buffer, 2.7μl Gen Hybridization buffer enhancer, 5.8μl nuclease free water). Seventeen microliters of beads from the previous step were added to the hybridisation reaction and incubated in a thermocycler at 65°C for 45 minutes (lid temperature 70°C) mixing the reaction every 10 minutes. After incubation, the samples were washed following manufacturer’s instructions. Briefly, 100μl of 65°C wash buffer 1 was added to the sample and carefully mixed. The supernatant was transferred into a new tube and kept on ice (cDNA for 10x gene expression library – GEx fraction). Beads were then washed by adding 150μl of 65°C stringent buffer, mixed well and incubated in a thermocycler for 5 minutes at 56°C (lid temperature 70°C). Next, the supernatant was discarded, and the previous step was repeated for a total of two washes. 150μl of wash buffer 1 was added to the beads, incubated for 2 minutes at room temperature, supernatant was then discarded, 150μl of wash buffer 2 was added to the beads and incubated for 2 minutes at room temperature. After discarding the supernatant, 150μl of wash buffer 3 was added to the beads and incubated for 2 minutes at room temperature. To elute H2B-GFP cDNA from the beads, 100μl of 100mM NaOH 0.1mM EDTA was added to the complex beads/cDNA and incubated 15 minutes at room temperature in a ThermoMixer C (300 rpm). The supernatant was transferred into a new tube and 10μl 1M HCl, 10μl 1M Tris-HCl (pH 7.5) was added. Both H2B-GFP cDNA fractions (lineage tracing fraction) as well as the fraction that contained the rest of the cDNA for the 10x gene expression library (GEx fraction) were purified using Dynabeads MyOne Silane beads (Invitrogen, cat. 37002D) following the protocol described above. 10x GEx libraries were generated using 120ng of the cDNA fraction following 10x Chromium protocol (Fragmentation, End repair and Library construction steps). H2B-GFP and HTO libraries were generated by amplifying either H2B-GFP or HTO cDNA using primers that specifically recognise H2B-GFP or HTO on the 5’-end and Read 1 on the 3’-end, generating standard Illumina paired-end sequencing constructs containing P5 and P7 (Supplementary Table 1). Final sequencing library quality was assessed by Fragment Analyser 5200 (Agilent, cat. M5310AA). Libraries were sequenced on a NovaSeq 6000 version 1.5 (Illumina, cat. 20013850). Sequencing depth: GEx library – 35000/50000 read pairs per cell, HTO library – 2000/3000 read pairs per cell, GFP and mCherry library – 10000/12000 read pairs per cell. Sequencing type: paired-end, dual indexing. Sequencing read: Read 1 – 28 cycles, i7 Index – 10 cycles, i5 Index – 10 cycles, Read 2 – 90 cycles.

### 10x Gene expression libraries and sequencing

10x libraries for HA1E (10E, A7A, A9H, B2F and B5F clones), HA1ER (F12, 1D12, 1C9, 2A6, 1F8, 3C5, 1E8, 2B12 and 1F5 clones) and HepG2 (1A, 32G, 14F, 48E and 44D clones) and for cells following gapmer-mediated knock-down were prepared following manufacturer’s instructions. In order to perform a direct comparison of scRNA-Seq output avoiding mathematical integration cell preparations from each cell system analysed were multiplexed using HTOs (BioLegend, cat. A0251-A0260) prior to library construction. Libraries were sequenced on a NovaSeq 6000 version 1.5 (Illumina, cat. 20013850). Sequencing depth: GEx library – 35000/50000 read pairs per cell, HTO library – 2000/3000 read pairs per cell. Sequencing type: paired-end, dual indexing. Sequencing read: Read 1 – 28 cycles, i7 Index – 10 cycles, i5 Index – 10 cycles, Read 2 – 90 cycles.

### Cell cycle analysis

Cells were plated at a confluency of 1 × 10^6^ cells/6010mm^2^ surface. Twenty-four to 48 hours later, cells were collected, fixed and permeabilised with methanol. Cell cycle progression was assessed using DAPI nuclear counterstain (Sigma-Aldrich, cat. MBD0015) and analysed by flow cytometry (BD LSRFortessa X-20 Cell Analyzer, BD Biosciences or Attune Nxt, Thermo Fisher Scientific).

### Bright field imaging

Cells were plated at a confluency of 1-1.5 10^6^ cells/10cm Petri dishes (6010mm^2^ surface). Twenty-four hours later, cells were imaged using a fluorescent microscope (Thermo Fisher Scientific, cat. AMEX1000).

### Cell proliferation

Cells were plated in standard 2D culture conditions at a defined number per well (HA1ER: 50,000 cells per 9.6cm^2^, HA1E: 120,000 cells per 9.6cm^2^, HepG2: 150,000 cells per 9.6cm^2^) and over a period of 4 days, three wells per clone were harvested each day and counted manually using a Neubauer counting chamber.

### Anchorage-independent growth in 3D soft agar

0.6% and 1% Difco agar noble solutions were prepared (BD Biosciences, cat. 214230) and autoclaved. Agar solutions were melted and equilibrated to 42°C in a water bath together with 2X growth medium (HepG2: DMEM high glucose (Sigma-Aldrich cat. D7777), 0.7% sodium bicarbonate (Sigma-Aldrich, cat. S8761), 20% FCS and 100U/ml Penicillin-Streptomycin (Thermo Fisher, cat. 15070063); HA1ER/HA1E: DMEM (Sigma-Aldrich cat. D5523), 0.7% sodium bicarbonate, 20% FCS and 100 μg/ml gentamicin (Thermo Fisher, cat. 15750037)). The lower 0.5% agar layer was generated by mixing the 1% agar solution 1:1 with the 2X medium and 1.5ml was added to each well of 6-well plates. After solidification of the bottom layer, the top 0.3% agar layer was prepared by mixing the 0.6% agar solution 1:1 with the 2X medium containing a fixed number of cells in a single-cell suspension (HA1ER: 10,000 cells/well, HA1E: 75,000 cells/well, HepG2: 22,500 cells/well) and 1.5ml was added per well. Once the top layer was set, 1ml of normal growth medium was added per well and 0.5ml of growth medium was added to each well every 3-4 days. For TGFβ-treated HepG2 samples, the growth medium was supplemented with 5ng/ml TGFβ. Colony growth was assessed by crystal violet staining after 2-3 weeks (HA1ER/HA1E: 3 weeks). For this, wells were fixed and stained with 0.1% crystal violet (Sigma-Aldrich, cat. C6158) and 10% ethanol for 30min, followed by multiple washes over several days. Finally, plates were imaged on Optronics GelCount at 600dpi and processed for colony counting using ImageJ (version 2.0.0).

### Transwell assay

Serum-free growth medium was used to preincubate transwells (Corning, cat. 3422) at 37°C during preparation of cells. Cells were trypsinised and resuspended in serum-free growth medium at a concentration of 1×10^6^ cells/ml. For TGFβ-treated HepG2 samples, medium was supplemented with 5ng/ml TGFβ. 0.5ml of complete growth medium (+5ng/ml TGFβ for TGFβ-treated HepG2 samples) was added to each well of a 24well plate and transwells were transferred to these wells, ensuring no bubbles were present between the transwell membrane and the lower medium. Finally, 100 μl of the cell suspension (100,000 cells) was added to the upper chamber of the transwell and incubated for 72 hours at 37°C. A cotton swab was used to remove non-migrated cells in the upper chamber, before migrated cells were fixed and permeabilised in ice-cold methanol for 10 minutes. Cells were stained in PBS supplemented with 10ug/ml DAPI (Sigma-Aldrich, cat. MBD0015) for 15min at room temperature and imaged using a 4x objective (EVOS FL Cell Imaging System). Adobe Photoshop was used to stitch together individual images covering the entire well and migrated cells were quantified using ImageJ.

### TRAIL treatment and PARP labelling

Cells were plated at a confluency of 1 × 10^6^ cells/6010mm^2^ surface. Twenty-four hours later, HA1ER-derived clone F12 and subclones 1F8 and 1C9 were challenged with recombinant human TRAIL (rhTRAIL, 1μg/ml, R&D Systems, cat. 375-LT-010) for 4 hours. TRAIL-induced apoptosis was determined by assessing the percentage of cleaved PARP positive cells using flow cytometry (BD LSRFortessa X-20 Cell Analyzer, BD Biosciences) and FlowJo software (BD, version 10.8.0) was used for data analysis. Briefly, TRAIL-treated cells were collected and fixed with ice-cold methanol. Methanol fixated cells were washed once with 1x PBS and incubated in blocking solution (0.5% BSA PBS) for 1⍰hour at 20°C. Immunolabelling of cleaved PARP was performed using fluorescently labelled Cleaved PARP Alexa 647 antibody (1:200 dilution, Cell Signalling, cat. 6987) for 1⍰hour at 20°C, followed by two washes in blocking solution (0.5% BSA PBS).

### TGFβ treatment

Prior to TGFβ treatment, HepG2 cells were plated in a 6-well plate at 150T cells per well. The next day, TGFβ was diluted in 10 μl medium per well and added to reach a final concentration of 5 ng/ml. As vehicle control, the same volume of water was diluted in 10 μl medium per well and added. Cells were incubated at 37 °C for 48h, before exchanging the growth medium supplemented again with 5 ng/ml TGFβ. After additional incubation for 48h at 37 °C, cells were collected for analysis or plated without TGFβ for withdrawal experiments.

### Western blot

Cells were lysed in RIPA buffer and whole cell extracts were quantified by Bradford assay to homogenize protein concentration across samples prior to protein separation by 10% SDS PAGE (including PageRuler Plus Prestained Protein Ladder, Thermo Fisher Scientific, cat. 26619) and transfer to nitrocellulose membranes. Membranes were incubated in 5% milk in PBS-T (0.1% Tween in PBS) for 1 h at 20°C and then washed 3 times in PBS-T for 5min. The membranes were incubated with antibodies against N-cadherin (1:2000) or α-Tubulin (1:8000) in 5% milk at 4°C overnight. Membranes were washed 3 times in PBS-T for 5 min and then incubated with the corresponding secondary antibody (1:10,000) for 1 h. Finally, membranes were washed 3 times in PBS-T for 5min and imaged on a LI-COR Odyssey imaging system.

### Generation of a lentiviral library containing 26 RAS variants with allocated molecular barcodes in their H2B-GFP reporter

As described previously^8^, gBlock DNA fragment encoding the H2B-GFP fusion containing a NdeI site in the 3’UTR and matching restriction sites at the ends for further downstream cloning into pLEX_307 (gift from David Root; Addgene, cat. 41392) (KpnI and SbfI) was amplified by PCR and cloned into the pGEM-T Easy vector system. Following verification of the construct by Sanger sequencing, this H2B-GFP fragment was further sub-cloned into the pLEX vector via One Shot ccdB Survival 2 T1 bacteria strain (Invitrogen, cat. A10460). To introduce a molecular barcode into the 3’UTR of the H2B-GFP gene, a gBlock DNA fragment with compatible restriction sites (NdeI and SbfI) encoding a modified version of the plasmid sequence downstream of the H2B-GFP stop codon was amplified by PCR using a primer with 12 random nucleotide barcodes split into three segments by intervening nucleotides (NNNNGCGNNNNTGANNNN), and further cloned into the pGEM-T Easy vector. Pooled plasmid DNA purification was performed (Qiagen Plasmid Midi Kit cat. 12143), the plasmid mix was digested using NdeI and SbfI for sub-cloning the barcoded fragment into the previously generated pLEX-H2B-GFP plasmid, and in-house generated electrocompetent One Shot ccdB Survival 2 T1 bacteria were transformed. Approximately 50,000 bacteria colonies were pooled for plasmid DNA purification of the barcoded pLEX-H2B-GFP plasmid mix. RAS variants were obtained from the Target Accelerator Pan-Cancer Mutant Collection (Addgene, cat. 1000000103). Individual liquid cultures of bacteria expressing pDONR plasmids encoding each of the 26 RAS variants included in our assay were pooled according to their OD_600_ to achieve a balanced representation, and then pDONR-RAS plasmid mix was purified using Qiagen Plasmid Midi Kit (cat. 12143). Following this, the pDONR-RAS plasmid mix was used to shuttle the oncogene encoding sequences into the previously generated pLEX-H2B-GFP plasmid mix by recombination using Gateway LR Clonase II Plus enzyme (Invitrogen, cat. 12538120) following the manufacturer’s instructions. A unique molecular barcode was assigned to each RAS variant by (Sanger) sequencing the barcode region in the 3’UTR of H2B-GFP as well as the RAS variant region. Finally, liquid cultures of the bacteria clones containing the 26 final lentiviral transfer vectors (pLEX-H2B-GFP-BC-RAS: HRAS-A11D, HRAS-G12A, HRAS-G12C, HRAS-G12D, HRAS-Q61K, HRAS-Q61L, HRAS-Q61R, HRAS-wt, KRAS-G12A, KRAS-G12C, KRAS-G12D, KRAS-G12V, KRAS-G12S, KRAS-D33E, KRAS-A59G, KRAS-E62K, KRAS-wt, NRAS-G12A, NRAS-G12C, NRAS-Q61H, NRAS-Q61K, NRAS-Q61L, NRAS-Q61R, NRAS-Y64D, NRAS-wt) were pooled based on their optical density and plasmid DNA was purified, rendering the transfer vector mix for lentivirus production.

Determination of the functional lentiviral titre and transduction of HA1E cells with 26 RAS variants Increasing volumes of lentivirus stocks (0-500 μl) were added to 150,000 HA1E cells and supplemented with polybrene (Sigma, cat. H9268-5G) to a final concentration of 2 μg/ml. The percentage of GFP positive cells was determined 48 hours post-transduction by flow cytometry (BD LSRFortessa X-20 Cell Analyzer, BD Biosciences) and FlowJo software (BD, version 10.8.0) was used for data analysis. The functional viral titre, calculated as the number of viral particles (i.e. transducing units (TU)) capable to transduce a particular cell line per volume was calculated using the following formula:

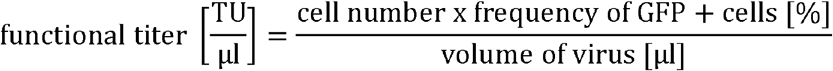

Based on this, HA1E cells were transduced with the pooled lentiviral library containing 26 RAS variants at a multiplicity of infection (MOI) of 0.05, supplemented with polybrene at a final concentration of 2μg/ml. Forty-eight hours post-transduction, FACS was used to collect GFP-positive cells (BD InFlux Cell sorter or FACSAria Fusion Cell Sorter, both BD Biosciences) with a purity of >90% and sorted cells were re-plated in standard growing conditions.

### Single-cell data and BdLT-Seq analysis

10x Chromium single-cell data was analysed using Seurat v4.0^76^. HTO-labelled cells were demultiplexed using the HTODEMUX module of Seurat and further processed for quality control and cell filtering taking into account nFeature_RNA, nCount_RNA and mitochondrial content. Further pre-processing included the mathematical regression of cell cycle related gene expression levels to remove cell cycle related bias in cluster processing. Similarly, variance in cluster detection due to high levels of mitochondrial transcripts present per cell was also regressed prior to final clustering. HTO and BdLT-Seq data was processed using CITE-seq-count^77^ prior to demultiplexing or barcode counting. BdLT-Seq barcode reads were pre-processed to remove sequencing artefacts, flanking and intervening sequences using cutadapt v3.7^78^. Clean barcodes were then processed using CITE-seq-count to generate lineage tracing files. A custom Node script^8^ was used to determine lineage relationships based on a similarity score considering the number of similar barcodes between two cells and data obtained by these means was used to build a directional lineage tree. Lineage visualisation was generated using CIRCOS^79^. All data analysed across the manuscript was pre-processed/processed using R scripts. Final image processing and output was obtained using Adobe Illustrator.

### Whole Exome Sequencing

Founder clones and thereof derived subclones from each cellular system were evaluated to assess possible genetic divergence at the exome level (WES, Whole Exome Sequencing. 100x coverage). Briefly, cells were grown as described above and processed for DNA purification using the QIAmp DNA mini kit (Qiagen. Cat 51304). High quality DNA Samples were then either processed for exome capture using the SureSelect Human All Exon V6 (Agilent, cat 5190-8872) and then processed for DNBSeq compatible library construction (BGI) or directly streamlined into DNBSeq library construction (BGI). WES sequencing was performed following a 100PE sequencing protocol in a DNBSeq instrument (BGI). WES was mapped to hg38 from each clonal lineage analysed, piled-up using mpileup^80^ (-q1 -Q30) and further processed through VarScan Somatic pipeline^81^ (--min-coverage 50). Chromosome level data visualisation was generated using chromo_plot from the vcfR package^82^.

### Whole Genome Sequencing

Founder clones (HA1ER, clone F12 and HepG2 clone 1A) and thereof derived subclones from each cellular system (HA1ER, clones 1F8 and 1C9; HepG2, clones 14F and 44D) were evaluated to assess possible genetic divergence at the whole genome level (WGS, Whole Genome Sequencing. 30x coverage). Briefly, cells were grown as described and processed for DNA purification using the QIAmp DNA mini kit (Qiagen. Cat 51304). High quality DNA Samples were then directly streamlined into DNBSeq library construction (BGI). WGS sequencing was performed following a 100PE sequencing protocol in a DNBSeq instrument (BGI). WGS was mapped to hg38 from each clonal lineage analysed, piled-up using mpileup^80^ (-q1 -Q30) and further processed through VarScan Somatic pipeline^81^ (--min-coverage 50). Chromosome level data visualisation was generated using chromo_plot from the vcfR package^82^.

### SLAM-Seq and SLAM-DUNK

SLAM-Seq was used to determine the fraction of nascent RNAs in contrast to steady-state RNA levels for each clonal lineage (at least 3 biological triplicates). Briefly, the founder clone and its corresponding subclones from each system were pulsed with 10μM 4-thiouridine (4-SU. Sigma, cat. 4509) for 15 min (1F8 vs 1C9) or 45 minutes (5 clones of HA1ER, HA1E and HepG2) and collected in TRIzol (Invitrogen, cat. 15596-026). After RNA quality control assessment (Fragment Analyzer 5200, Agilent, cat. M5310AA) and mass determination (Qubit, Invitrogen), 5 μg of RNA from each sample was incubated in 4-SU conversion buffer (10 mM IAA in H_2_O, 50 mM sodium phosphate pH 8, 0.15mM DTT, 50% DMSO (v/v) and H_2_O to a final volume of 50 μl) for 15 min at 50°C on a heat block. Immediately after, the conversion reaction was stopped by addition of 2 μl of 1M DTT and EtOH precipitated over-night at -80°C (adding 200μl H_2_O, 2 μl glycogen (20 mg/ml), 30 μl 3M pH 5.2sodium acetate and 900 μ l ice-cold absolute EtOH). Retrieved RNA was further processed to build an Illumina compatible strand-specific RNA sequencing library using either NEBNext Ultra II Directional RNA Library Prep Kit for Illumina (NEB, cat. E7760S, for 1F8 vs 1C9) or QuantSeq 3⍰ mRNA-Seq V2 Library Prep Kit FWD with UDI (Lexogen, all clonal systems) kits following manufacturer’s instructions. Pooled libraries were sequenced on a NovaSeq 6000 (3’ 150bp) aiming to sequence 30M reads per sample. SLAM-Seq sequencing data was processed using a T>C aware mapper (NextGenMap^26^) and processed following SLAM-DUNK pipeline^27^ (NextGenMap align to hg38 reference genome, mapping quality filter, snp annotation filtering and modified reads count). Final count tables were further processed using the beta binomial test to determine transcriptional differences from steady-state RNA levels (FDR<0.05, FC>2). Data visualisation was generated using heatmap.2 (R v4.2.0).

### Chromatin accessibility (ATAC-Seq)

ATAC-seq was performed for all clones evaluated herein (HA1E: 10E, A7A, A9H, B2F and B5F; HA1ER: F12, 1F8, 1C9, 2A6 and 1D12; HepG2: 1A, 32G, 14F, 48E and 44D) following the Omni-ATAC protocol^83^ but using an in-house prepared Tn5 transposase^84^. Briefly, 75,000 cells were processed for ATAC-Seq as follows: HA1E and HA1ER clones were seeded at 3.5 × 10^5^ cells per 6-cm dish whilst HepG2 clones at 5.25 × 10^5^ cells and grown for 2 days. Next, cells from every system were rinsed twice with 4ml PBS and then trypsinised and collected by centrifugation (250 g, 4 min), resuspended in PBS, placed on ice and counted in a Neubauer chamber (Marienfeld, cat. 0640030). Aliquots containing 75,000 cells were transferred to pre-chilled DNA low-binding tubes, centrifuged (350 g, 5 min, 4 °C; swing-bucket rotor with FACS-tube adaptor, Eppendorf 5810 R) and resuspended in 75 µl ice-cold ATAC Lysis Buffer (9.7 mM Tris-HCl pH 7.5, 9.7 mM NaCl, 2.91 mM MgCl_2_, 0.1% (wt/vol) IGEPAL CA-630, 0.1% (wt/vol) Tween-20, 0.01 % (wt/vol) digitonin). After 3 min on ice, lysates were diluted with 1 ml ice-cold ATAC Wash Buffer (9.9 mM Tris-HCl, 9.9 mM NaCl, 2.97 mM MgCl_2_, 0.1% Tween-20), mixed by inversion and nuclei were collected by centrifugation (500 g, 10 min, 4 °C). Next, cell pellets were gently resuspended in 75 µl ice-cold transposition buffer (10 mM Tris-HCl pH 7.6, 5 mM MgCl_2_, 10 % (vol/vol) dimethylformamide, 33 % PBS, 0.1% Tween-20, 0.01% digitonin, 1.25 µM loaded Tn5) and incubated at 37°C in a Thermomixer comfort at 1000 rpm for 30 min. Transposition reactions were stopped by the addition of 375 µl of DNA Binding Buffer (DNA Clean & Concentrator-5 kit, Zymo Research D4013) and further processed for transposed DNA enrichment using DNA Clean & Concentrator-5 kit (Zymo Research D4013). Finally, transposed DNA was eluted in 21 µl Elution Buffer into DNA low-binding tubes. Next, for multiplexing purposes, barcoding PCR reactions were conducted for each sample. Each PCR reaction contained 25 µl NEBNext Ultra II Q5 2× Master Mix (M0544S, New England Biolabs), 2.5 µl 5 µM i5 index primer, 2.5 µl 5 µM i7 index primer, and 20 µl tagmented DNA and were amplified as follows: 72°C 5 min; 98 °C 30 s; 13 cycles (HA1ER, HepG2) or 14 cycles (HA1E) of 98°C 10 s, 65°C 1 min 15 s; final extension 65°C 3 min, hold 4°C.

Barcoded PCR products (sequencing-ready library) were subjected to size selection using Mag-Bind TotalPure NGS beads (range 150 – 1000 nt. M1378-00, Omega Bio-tek, Inc.) and then assessed on an Agilent 4200 TapeStation using D1000 ScreenTape and D1000 Reagents (35–1,000 bp detection range). Validated libraries were then sequenced in a NextSeq2000 apparatus generating a 50PE reads for each sample. Obtained reads were trimmed (Tn5 adaptor sequence: CTGTCTCTTATACACATCT) using cutadapt^78^ prior to mapping to hg38 using Bowtie2^85^. After mapping peak calling was performed (FDR<0.05) using MACS2 v2.2.7.1^86^ pipeline. Finally, a catalogue containing all peaks identified on every sample from each cellular system analysed was built to extract genomic coordinates used for heatmap visualisation (Deeptools v3.5.1^87^).

### Label-free proteomic analysis (LC-MS-MS)

Cell lysis was performed in 5% SDS, 50 mM TEAB followed by protein content measurement using the 2D quant kit (Cytiva, cat. 80-6483-56). Then, 200 μg of protein from each sample was reduced with 5mm TCEP at 55°C for 15 min followed by alkylation with 15mM MMTS for 10 min at 20°C. Samples were then loaded onto STRAP minis (Protifi, cat. C02-mini) and digested with 20 μg sequencing grade trypsin (Sigma, cat. TRYPSEQ-RO) for 18 h at 37°C. Following STRAP elution, peptide digests were dried to completeness in a vacuum centrifuge at 45°C before resuspension at 800ng/μl on 0.2% formic acid.

LC-MS was performed in technical quintuplicate in a RSLC HPLC system (mobile phase A: water 0.1% formic acid; mobile phase B: acetonitrile) coupled to an Orbitrap Lumos mass-spectrometer via an EasySpray source (Thermo Fisher Scientific). Peptides were injected directly onto an EasySpray column (50 cm long, 7 μm ID pepmap C18) at 180 nl/min in 1% mobile phase B and were separated at 180 nl/min with a gradient of 1-24% over 90 min. LC eluent was sprayed directly into the MS source at an ion spray voltage of 1.9Kv. The MS was operated in data dependent mode where an MS1 survey scan was acquired in the Orbitrap detector at 120K resolution (at m/z 200) in the mass range of 350-1200m/z at a target value of 2e5 ions and a maximum fill time of 40ms. All available charged ions were selected for MS2 by HCD at a normalised collision energy of 35% with product ions detected in the Orbitrap detector at a resolution of 30K, 54ms max fill time and a target value of 1e5 ions. A maximum cycle time of 1.2s was allowed.

### TMT-proteomics

Cell lysis was performed in 5% SDS, 50mM TEAB followed by protein content measurement using the 2D quant kit (Cytiva, cat. 80-6483-56). Then, 200 μg of protein from each sample was reduced with 5 mm TCEP at 55 °C for 15 min followed by alkylation with 15 mM MMTS for 10 min at 20°C. Samples were then loaded onto STRAP minis (Protifi, cat. C02-mini) and digested with 20 μg sequencing grade trypsin (Sigma, cat. TRYPSEQ-RO) for 18 h at 37°C. Following STRAP elution, peptide digests were dried to completeness in a vacuum centrifuge at 45°C. Peptide digests were resuspended in 500 mM TEAB at a concentration of 2 μg /μl and TMT 10-plex labelled according to the manufacturer’s instructions (Thermo Fisher Scientific). Reactions were quenched by addition of 8 μl 5% hydroxylamine prior to pooling. Samples were acidified to pH 2.7 by addition of trifluoracetic acid before revers phase solid phase extraction.

Pooled, acidified peptides were loaded onto C18 RP cartridges (Sep-Pak 500mg tC18, Waters, cat. 186004619), washed with 10 ml 0.1 % formic acid before elution in 500 μl 60% acetonitrile, 0.1% formic acid. 10% of the pooled sample was aspirated off, the remaining 90% of peptide was split equally into 2 tubes and all aliquots were dried to completeness in a vacuum centrifuge at 45°C. Approximately a 10 % aliquot (200 μg) was reserved for proteomics profiling and both 45% aliquot (900 μg) were resuspended for subsequent enrichment either by TiO2 (Thermo Fisher cat. A32993) or FeNTA enrichment (Thermo Fisher cat. A32992). Following elution from either TiO2 or FeNTA columns, samples were dried to completeness in a vacuum centrifuge at 20°C followed by resuspension in 5 μl of 0.05% TFA and were ready for LC-MS-MS analysis.

LC-MS was performed in technical quintuplicate in a RSLC HPLC system (mobile phase A: water 0.1% formic acid; mobile phase B: acetonitrile) coupled to an Orbitrap Lumos mass-spectrometer via an EasySpray source (Thermo Fisher Scientific). Following, peptides/phospho-peptides were injected directly onto an EasySpray column (50 cm long, 75 μm ID pepmap C18) at 180 nl/min in 1% mobile phase B. Peptides were separated at 180 nl/min with a gradient of 1-24% over 90 min. LC eluent was sprayed directly into the MS source at an ion spray voltage of 1.9 Kv. The MS was operated in data dependent mode where an MS1 survey scan was acquired in the Orbitrap detector at 120 K resolution (at m/z 200) in the mass ranges of 350-1200 m/z at a target value of 2e5 ions and a maximum fill time of 40 ms. All available charged ions were selected for selected for MS2 by HCD at a normalised collision energy of 42%. Product ions were detected in the Orbitrap detector at a resolution of 50K, 86 ms max fill time and a target value of 1e5 ions. A maximum cycle time of 4s was allowed.

### Label-free proteomics data processing

Raw data was imported into Progenesis QI proteomics pipeline (Waters Nonlinear Dynamics) where peak detection was performed at maximum sensitivity. Protein relative quantitation was performed using the top 3 method. Proteins were identified by extraction of a mascot generic file (MGF) that was exported to Mascot (Matrix science) with the following search criterion. Species human, specificity was trypsin allowing up to 2 missed cleavages, precursor mass accuracy 10ppm, product ion sensitivity 20 ppm, fixed modification Methylthio (C) variable modifications oxidation (M) deamidation (NQ) searched against Uniprot_Swissprot_2022 using a decoy FDR method limited at 1%. Protein identities were imported back into Progenesis QI and data was further processed in R for visualisation purposes.

### TMT proteomic analysis (PEAKS)

Raw data was imported into Peaks Studio 10.6 (Bioinformatics Solutions Inc.) where both protein identification and quantitation were performed using the default TMT 10 plex template. Database search evaluated human acquired data against the Uniprot_Sprot_2022.fasta database using fixed modification of methylthio (C) variable modifications oxidation (M) deamidation (NQ) and the following advance settings: merge scans [DDA], correct precursor [DDA], mass only, associate features with chimera scan [DDA], filter features charge between 2 and 4 10 ppm precursor mass tolerance, fragment ion 0.1 Da, digest mode specific trypsin, max missed cleavages per peptide 2, estimated FDR by decoy-fusion, quant TMT 10 plex, MS2 quant tolerance 20 ppm, FDR threshold 1%.

### TMT phospho-proteomic analysis (PEAKS)

Raw data was imported into Peaks Studio 10.6 (Bioinformatics Solutions Inc.) where both protein identification and quantitation were performed using the default TMT 10 plex template. Database search evaluated human acquired data against the Uniprot_Sprot_2022.fasta database using fixed modification of methylthio (C), variable modifications oxidation (M), deamidation (NQ), phosphorylation (STY) and the following advance settings: merge scans [DDA], correct precursor [DDA], mass only, associate features with chimera scan [DDA], filter features charge between 2 and 4 10ppm precursor mass tolerance, fragment ion 0.1Da, digest mode specific trypsin, max missed cleavages per peptide 2, estimated FDR by decoy-fusion, quant TMT 10 plex, MS2 quant tolerance 20ppm, FDR threshold 1%. Differentially represented phosphorylation sites between subclones were identified by applying the beta binomial distribution (FDR<0.05, FC>0.4) using countData package (R).

### CyTOF

HA1ER cells (clones F12, 1C9 and 1F8; 3×10^6^ each) were pulsed for 30 min with IdU following manufacturer’s instructions (Maxpar cell cycle panel kit, cat. 201313) and then collected by trypsinisation. Cells were next spun-down and resuspended in 1ml 1X Maxpar Fix I Buffer (Fluidigm, cat. 201065) for 10 min at 20°C. Then cells were spun down and resuspended in 7 ml Maxpar Cell Staining Buffer (Fluidigm, cat. 201068), mixed gently and further incubated for 10 min at 20°C. Next, cells were spun down once again and after repeating this washing step, cells were resuspended in 50 μl of Maxpar Cell Staining buffer and then permeabilised with ice-cold methanol for 15 min on ice. Cells were then washed twice with 2 ml Maxpar Cell Staining Buffer +EDTA (Maxpar Cell Staining Buffer supplemented with 5mM EDTA) and spun down for 5 min at 800g. Immediately after, 1 μl of each metal isotope-labelled specific antibody (MOT4, JIP4, SYUA, DCXR, SQSTM, MYEF2, Hsp20, pRb, CyclinB1, Ki67, pH3 (Supplementary Table), using Maxpar X8 Antibody Labeling Kit (Fluidigm, cat. 201144A, 201167A, 201155A, 201171A, 201176A, 201147A, 201159A)) was added to Maxpar Cell Staining Buffer +EDTA and a final volume of 50 μl was added per sample and incubated for 30 min at 20°C. Following incubation, cells were washed twice with 2 ml Maxpar Cell Staining Buffer +EDTA and further spun down at 800 g for 4 min. Immediately after, supernatant was removed and 1ml of 4% PFA and 1 μl Iridium were added to each sample and stored at 4°C for 16 h. Next day, samples were washed in 2 ml of Maxpar Cell Staining Buffer +EDTA and spun down. Finally, samples were washed twice in Maxpar Cell Acquisition Plus Solution (Fluidigm, cat. 201244). Samples were analysed in a Helios apparatus and data was processed using flowCore v2.16.0^88^, CATALYST v1.28.0^89^ and visualised in R v4.2.0.

### Gapmer-mediated knock-down

Antisense LNA gapmers were designed against lncRNA transcript sequences (Ensembl release 105^90^) using the Qiagen Antisense LNA GapmeR custom builder (Supplementary Table). One day prior to transfection, 100T cells (HA1ER) or 150T cells (HepG2) were seeded into a 6well plate in 2ml growth medium. Next day, around 450 μl of medium were removed from each well for a final volume of 1.5ml per well. 40 μM gapmer was mixed with 75 μl OptiMEM (Invitrogen, cat. 31985062) per well to be transfected. The X-tremeGENE 360 transfection reagent (Roche, cat. 8724105001) was equilibrated to 20°C and briefly vortexed before immediately adding 2.5 μl per well to be transfected to the gapmer mix. After thorough mixing and incubation at for 30min at 20°C, 75 μl of the transfection mix was added dropwise to each well of cells (final gapmer concentration in 1.5ml: 26 nM). Following gentle mixing, cells were incubated at 37°C. The growth medium was exchanged 24h post-transfection. Cells were used in RT-qPCR, scRNA-seq or RPPA assay 48h post-transfection, only for DPP-DT-G2 cells were used 24h post-transfection.

### Nuclear-Cytoplasmic fractionation for HA1ER clones

HA1ER clones F12, 1F8 and 1C9 were seeded at 1×10^6^ onto a 10 cm culture dish and were grown to ∼70% confluency. Cells were rinsed twice with 10 ml ice-cold PBS and trypsinised with 1 ml of 0.25% trypsin. Cells were pelleted by centrifugation at 220g for 3 minutes at 4°C and then washed twice in 1 ml ice-cold PBS, counted, and 2×10^6^ cells were then taken for further fractionation. Pellets were resuspended in 100 µl of hypotonic buffer (20 mM Tris pH 7.6, 0.1 mM EDTA, 2 mM MgCl_2_, 0.5 mM NaF, and 0.5 mM Na_3_VO_4_) supplemented with 1x cOmplete EDTA-free Protease Inhibitor Cocktail (Roche, cat. 11873580001), 1 mM phenylmethylsulfonyl fluoride (PMSF), and 260 U/ml RiboLock RNase Inhibitor (Thermo Fisher Scientific, cat. EO0381), and transferred to a fresh 2ml DNase/RNase-free DNA-low binding tube. Cells were then incubated for 2 minutes at 20°C, then for 10 minutes on ice to induce hypotonic swelling, followed by addition of 10 µl of 10% IGEPAL CA-630 (Sigma-Aldrich, cat. I8896) and pipetted up-and-down approximately 20 times. Nuclei, unbroken, or partially lysed cells were pelleted at 550g for 3 minutes at 4°C and then 80% of the supernatant was taken from the top (containing the cytoplasmic fraction), transferred to a fresh tube. Next, 150 µl of hypotonic buffer was added to the pellet and remaining cytoplasmic extract; the pellet was resuspended and once again spun down. 80% of the supernatant was taken from the top and combined with the cytoplasmic fraction. Nuclei pellets were then resuspended in hypotonic buffer in a volume equal to the final cytoplasmic fraction (373 µl). 20 ng of spike-in eGFP transcript for HA1ER cell system (*in-vitro transcribed* from an eGFP encoding plasmid. HighYield T7 RNA Synthesis Kit, Jena Bioscience, cat. NT-101) was added to each fraction, and total RNA was extracted using 1 ml of TRIzol (Invitrogen, cat. 15596-026). 773 µl of the aqueous phase from each fraction was mixed with 25.5 µl of 3M NaOAc, 1.5 µl of 5 mg/ml linear acrylamide (Invitrogen, cat. AM9520), 1 volume of isopropanol, and RNA was precipitated overnight at -20°C. Pellets were washed with 75% ice-cold ethanol, dried, and re-dissolved in 12 µl of nuclease-free water. RNA concentrations were measured using Nanodrop.

### Nuclear-Cytoplasmic fractionation for HepG2 clones

HepG2 clones 1A, 14F and 44D were seeded at 1.5×10^6^ onto a 10 cm culture dish and were grown to ∼70% confluency. Cells were harvested and resuspended in ice-cold PBS and from thereon kept on ice. For each sample, 2×10^6^ cells were transferred to a fresh tube, pelleted and resuspended in 380 µl ice-cold hypotonic lysis buffer (HLB, 10mM Tris-HCl pH7.5, 10mM NaCl, 3mM MgCl_2_, 10% glycerol, 0.3% IGEPAL CA-630) with 100U Protector RNase inhibitor (Roche, cat. 3335399001) and incubated on ice for 10min. Following brief vortexing and pelleting by centrifugation, the supernatant was transferred to a fresh tube (cytoplasmic fraction). The pelleted nuclei were washed twice in 380 µl HLB and finally resuspended in 380 µl (nuclear fraction). 1 ng of a custom spike-in plasmid was added to each sample to allow for normalisation during RT-qPCR analysis. After adding 1ml TRIzol to each fraction, RNA was extracted according to the manufacturer’s instructions. To ensure quantitative comparisons between samples in RT-qPCR, care was taken to treat samples equally during RNA extraction, e.g. collecting equal volume of aqueous phase for each sample.

### Reverse transcription followed by qPCR

Total RNA was isolated using TRIzol (Invitrogen, cat. 15596-026) following manufacturer’s instructions. Reverse transcription was performed using QuantiTect Reverse Transcription Kit (Qiagen, cat. 205311) following the manufacturer’s instructions with 1 or 2μg RNA input. qPCR was performed using PowerUp SYBR Green Master Mix (Invitrogen, cat. A25742) in a QuantStudio3 Real Time PCR System (Thermo Fisher Scientific, cat. A28136). Data analysis was conducted using Real-Time qPCR Connect Data Analysis Tool (Thermo Fisher Scientific Connect Platform).

### FISH probes design and production

FISH probe amplicon regions between 260-300bp were determined based on transcript sequences (Ensembl release 105^90^) and primers to amplify these regions were then designed using the Connect OligoPerfect tool (Thermo Fisher). RNA purified from cells known to express target RNAs at high level was reverse transcribed with QuantiTect reverse transcription kit (Qiagen, cat. 205311) using oligo-dT primers following the manufacturer’s recommendations. cDNA was then amplified by the Expand high fidelity PCR system (Roche, cat. 11732641001) using specific primers and PCR-products were then resolved by agarose gel electrophoresis. Bands of the expected molecular weights were purified and quantified using the Qubit dsDNA BR assay kit (Invitrogen, ca. Q32850). 1μg amplicon DNA was then digested using Lambda exonuclease (NEB, cat. M0262) at 37°C for 1 hour, followed by heat inactivation at 75°C for 10 minutes. Digestion products were then isolated on streptavidin beads and then washed twice with 1X B+W buffer (5mM Tris-HCl pH7.5, 1mM EDTA, 2M NaCl). Next, beads were resuspended in 25μl amplification mix 1 (1μl dNTP mix (10mM dATP, 10mM dCTP, 10mM dGTP, 7.5mM dTTP and 2.5mM aminoallyl dUTP (Thermo Scientific, cat. R1101)), 2μl 20μM random hexamers, 22μl H_2_O) before the addition of 25μl amplification mix 2 (0.75μl Expand high fidelity enzyme mix (Roche, cat. 11732641001), 5μl 10X Expand high fidelity buffer with 15mM MgCl_2_ (Roche, cat. 11732641001) and 19.25μl H_2_O). Beads were then washed twice with H_2_O and resuspended in 25μl H_2_O. 15μl labelling buffer (25 mg/ml sodium bicarbonate (Sigma-Aldrich, cat. S5761) was added to the resuspended beads before addition of 10μl dye (Alexa Fluor 555/647 Reactive Dye (Invitrogen, cat. A32757, A32756) dissolved in 40 μl DMSO). Samples were vortexed briefly and incubated at room temperature for 1 hour blocked from light. Beads were then washed three times in H_2_O and FISH probes were released through chemical denaturation by resuspending beads in 100μl filtered denaturation buffer (100mM NaOH, 0.1mM EDTA pH8.0) and incubating at room temperature for 15 min blocked from light. The supernatant was separated from the beads using a magnetic separator and collected into a fresh tube. Probes were then precipitated from the supernatant using 1/10 vol sodium acetate, 2μl Ethachinmate (Nippon Gene, cat. 318-01793) and 2.5X vol 100% ethanol and then eluted in 20μl H_2_O. Probes stocks were stored at -80°C.

### RNA-FISH

Cells were grown to 70% confluency on glass coverslips in 24-well plates and treated according to experimental design. Cells were then washed twice with 1X ice-cold PBS and fixed with 3% formaldehyde (Thermo Scientific, cat. 28906) and 4% sucrose in 1X PBS for 15 minutes at 20°C. Following fixation, cells were washed three times in 1X PBS and permeabilised with 0.25% Triton-X in 1X PBS for 10 minutes at 20°C, followed by three washes in 1X PBS. Fluorescently labelled RNA-FISH probes were diluted in formamide (10μl total volume per coverslip) and denatured at 75°C for 7 minutes. 10μl 2X hybridisation buffer (4X SSC, 40% dextran sulphate (Sigma-Aldrich, cat. D8906), 2mg/ml BSA) per coverslip was then added to each probe dilution. 20μl probe mix was hybridised to each sample of cells in a Thermobrite 110/120 VAC hybridisation chamber (Abbott, cat. 07J91-010), using the incubation program as follows: 65°C for 5 minutes, 37°C for 16 hours. Following hybridisation, samples were washed three times in 50% formamide/2X SSC at 42°C for 5 minutes each, three times in 2X SSC at 42°C for 5 minutes each and three times in 1X SSC at 42°C for 5 minutes each. Samples were then counterstained with DAPI and washed twice in 1X SSC, before mounting in ProLong Gold Antifade Mountant (Molecular Probes, cat. P36930). Cells were analysed on a Zeiss LSM880 microscope equipped with Airyscan module.

### RNA-FISH coupled to ICC

Cells were grown to 70% confluency on glass coverslips in 24-well plates and treated according to experimental design. Cells were then washed once with 1X PBS and fixed with 3% formaldehyde (Thermo Scientific, cat. 28906) and 4% sucrose in 1X PBS for 15 minutes at room temperature. Following fixation, cells were washed three times in 1X PBS and permeabilised with 0.25% Triton-X in 1X PBS for 10 minutes at room temperature, followed by three washes in 1X PBS. Cells were blocked in blocking buffer (1% BSA in 1X PBS for at least 2 hours at 20°C, before incubation with 1:200 primary antibody in blocking buffer overnight at 4°C. Cells were then washed twice with 0.1% Tween-20 in 1X PBS and incubated with 1:1000 secondary antibody in blocking buffer for 1 hour at 20°C, blocked from light. Cells were then washed twice with blocking buffer and then crosslinked again with 3% formaldehyde and 4% sucrose in 1X PBS for 10 minutes at 20°C. Next, cells were washed twice with 2X SSC (Sigma-Aldrich, cat. S6639) / 50% formamide (Sigma-Aldrich, cat. F9037). Next, fluorescently labelled RNA-FISH probes were diluted in formamide (10μl total volume per coverslip) and denatured at 75°C for 7 minutes. 10μl 2X hybridisation buffer (4X SSC, 40% dextran sulphate (Sigma-Aldrich, cat. D8906), 2mg/ml BSA) per coverslip was then added to each probe dilution. 20μl probe mix was hybridised to each sample of cells in a Thermobrite 110/120 VAC hybridisation chamber (Abbott, cat. 07J91-010), using the incubation program as follows: 65°C for 5 minutes, 37°C for 16 hours. Following hybridisation, samples were washed three times in 50% formamide/2X SSC at 42°C for 5 minutes each, three times in 2X SSC at 42°C for 5 minutes each and three times in 1X SSC at 42°C for 5 minutes each. Samples were then counterstained with DAPI and washed twice in 1X SSC, before mounting in ProLong Gold Antifade (Invitrogen, cat. P36930). Cells were visualised with a Zeiss LSM880 microscope equipped with Airyscan module. Images were obtained using ZEN software (Zeiss). ImageJ version 2.0.0 was used to split channel information. Adobe Photoshop 2025 was used to prepare pseudocoloured panels.

### Reverse Phase Protein Array (RPPA)

Cells were transfected and cultured as described in the section ‘Gapmer-mediated knock-down’. To prepare samples for RPPA, cells were kept on ice and washed twice with ice-cold PBS. Next, lysis buffer (1 % Triton X-100 (v/v), 0.05 M HEPES pH7.4, 1 mM EGTA ph7.5-8, 0.15 M NaCl, 1.5 mM MgCl_2_, 1 mM Na_3_VO_4_, 10 mM TSPP, 112.5 mM NaF, 10 % glycerol, 1 tablet/10ml cOmplete Mini EDTA-free protease inhibitor (Roche, cat. 11836170001), 1 tablet/10ml PhosStop phosphatase inhibitor (Roche, cat. 04906837001) was added and cells were scraped and collected into a tube. After a 20min incubation on ice, samples were centrifuged at 16,900 rcf for 10min at 4°C and the supernatant protein lysate was collected and stored at -20°C. Protein concentration was determined by Bradford assay. Samples were prepared for printing by addition of 4X Sample buffer (40% Glycerol, 8% SDS, 0.25 M Tris-HCl, 10% β-mercaptoethanol) and heat denaturation at 95°C for 5 min. All samples were diluted to 1.5 mg/ml (D1), 0.75 mg/ml (D2), 0.375 mg/ml (D3) and 0.1875 mg/ml (D4) using PBS containing 10% glycerol. Array spotting was carried out with the Quanterix 2470 Arrayer platform using 185 μM pins. Each sample was spotted at four dilutions (D1-D4) onto single pad ONCYTE SuperNOVA nitrocellulose slides (Grace Bio-Labs, cat. 705278) at a 500 μM spot-to-spot distance and in 3 technical replicates. 180 signalling pathway markers (Supplementary Table) were profiled in a standard RPPA assay as described briefly below^43^. After spotting, slides were incubated with antigen retrieval solution (1x Reblot strong. Merck Millipore, cat. 2504) for 10 min before being placed in a microfluidic structure to individually address the arrays with primary or secondary antibody solutions. Following blocking buffer (Superblock T20. Thermo Scientific, cat. 37536) for 10 min the detection of marker antibodies was performed in a two-step sequential assay. First the array is incubated with primary analyte-specific antibody in blocking buffer for 60 min at room temperature followed by the removal of excess antibody by washing arrays with 0.1% PBS-Tween followed by a further incubation with blocking buffer and PBS-T washes, and incubation with secondary antibody (Dylight-800-labeled anti-species antibodies diluted 1:2500 in Superblock T20) for 30 min. After further washing and slide drying, the arrays were imaged in the Innopsys Innoscan 710 scanner (Innopsys). Blank signals were determined by omitting the primary antibody and instead incubating the array with Superblock T20 alone, followed by secondary antibody incubation. Sample loading on arrays (for normalisation) was determined by staining one slide with fast green protein dye and scanning at 800 nm. Microarray images were analysed using Mapix software (Innopsys) with feature (spot) diameter of the grid set to 270 μm. Average signal intensities was determined for each individual feature and the median background from the adjacent area was subtracted from each feature signal leading to a net signal per feature. Fluorescence intensity for each feature (spot) on the array was measured and a test was performed for linear fit through the 4-point dilution series for all samples on all arrays using a flag system where R2 > 0.9 (green flag) is deemed good, >0.75 (amber flag) is deemed acceptable and <0.75 (red flag) is poor and was excluded from data analysis. The median values from the 4-point dilution series were calculated and used as a measure of fluorescence intensity. This median fluorescence intensity for each sample and target was normalised to the median fluorescence intensity of the fast green protein stain (total protein) and thereafter averaged across the technical triplicates. The final (phospho-) protein levels were averaged across both biological replicates and a threshold of ≥20% increase/decrease compared to the mean of the non-targeting control gapmer was applied to determine differential levels.

## Statistics and reproducibility

All histograms displayed in the manuscript show mean value⍰± ⍰standard deviation (SD) or propagated error with the number of replicates and, if performed, the method to determine statistical significance indicated in the figure legends.

As part of scRNA-Seq data pre-processing, cells found to be displaying multiple HTO (hashtag oligo) labelling after demultiplexing were excluded from further analysis. Moreover, to attenuate artefactual clustering, cells displaying high levels of mitochondrial transcripts and abnormal read counts were masked. Further processing included the mathematical regression of the datasets based on cell cycle-related genes expression to remove cell cycle-related bias in cluster mapping. Random sampling was applied to BdLT-Seq data libraries to optimise the number of barcodes needed to build lineage trees and to reduce computational burden.

All attempts of replication were successful. No statistical method was used to predetermine the sample size. For the analysis of the label-free MS in Fig. 3a, one sample from 1F8 (1_A) and one sample from 1C9 (1_A) were excluded from the analysis due to poor performance in the MS, presumably linked to a technical issue related to stabilisation/optimisation of the instrument. Given the fragile nature of the soft agar in the clonogenicity assays, some damaged wells were excluded from the analysis. The experiments were not randomised and the investigators were not blinded during experiments and outcome assessment.

## Data availability

All data used to generate figures in this study is provided as Source Data. Sequencing data have been deposited in the Gene Expression Omnibus (GEO) database under the accession code GSE223496. WES and WGS have been deposited the Short Read Archive (SRA) and can be retrieved under the accession code PRJNA1219504 and PRJNA1279408. The mass spectrometry proteomics and phospho-proteomics data have been deposited to the ProteomeXchange Consortium via the PRIDE^91^ partner repository with the dataset identifiers PDX063769, PXD064047, 10.6019/PDX063769 and 10.6019/PXD064047.

## Code availability

All data were analysed using publicly available, benchmarked, and validated pipelines (R v4.2.0, Seurat v4.0, CITE-seq-count v1.4.5, cutadapt v3.7, CIRCOS, Bowtie2 v.2.5.4, MACS2 v2.2.7.1, Deeptools v3.5.1, samtools v1.15.1, bcftools, VarScan v2.4.4, vcfR v1.15.0, SLAM-DUNK v0.4.3, NextGenMap v0.5.5, FlowCore v2.16.0, CATALYST v1.28.0). Data analysis rationale is described in the Methods section. For any data analysis-related inquiries, please contact M.M.P.

## Acknowledgements

We thank Valeria Pavet for helpful discussions and critical reading of the manuscript. We are also grateful to Richard Marais for his support. We also thank Naveed Khan for IT assistance. Y.S. was a Cancer Research UK—Manchester Institute postdoctoral fellow, B.B. and K.L.M. are recipients of a Cancer Research UK—Manchester Institute PhD studentship, A.M. is a Cancer Research UK-Scotland Institute postdoctoral fellow and M.M.P. was a Cancer Research UK Fellow and is currently a CRUK Scotland Institute Fellow and Senior Lecturer of the University of Glasgow. We acknowledge the HTPU microarray service at the Institute of Genetics and Cancer at the University of Edinburgh for their technical support with RPPA. We thank the Core Facilities at the CRUK Manchester Institute (Molecular Biology Core, Flow Cytometry, Biological Mass Spectrometry and Visualisation, Irradiation & Analysis) and Finance and Logistics Services at Cancer Research UK Manchester Institute and Scotland Institute. This research was funded by CRUK-MI core grant number C5759/A27412 and CRUK-SI core grant number A31287.

## Contributions

M.M.P. conceived the research project. Y.S., B.B., K.L.M. and M.M.P. designed and performed experiments. A.M. performed nuclear/cytoplasmic fractionation assay on HA1ER cells and ATAC-Seq on all clonal systems. R.B.J. prepared recombinant Tn5 for ATAC-Seq experiments. M.M.P. performed computational analysis. N.O.C. set up the RPPA platform and A.F.M. performed RPPA assays. Y.S., B.B., K.L.M. and M.M.P. wrote the Material & Methods section and prepared figures. M.M.P. wrote the manuscript. Y.S., B.B., K.L.M., A.M., R.B.J., A.F.M., N.O.C. and M.M.P. edited the final manuscript.

