## Supplementary Figures for "Non-genetic control of cell plasticity and phenotypic cell states is mediated by cytoplasmic IncRNA nucleated signalling pathways"

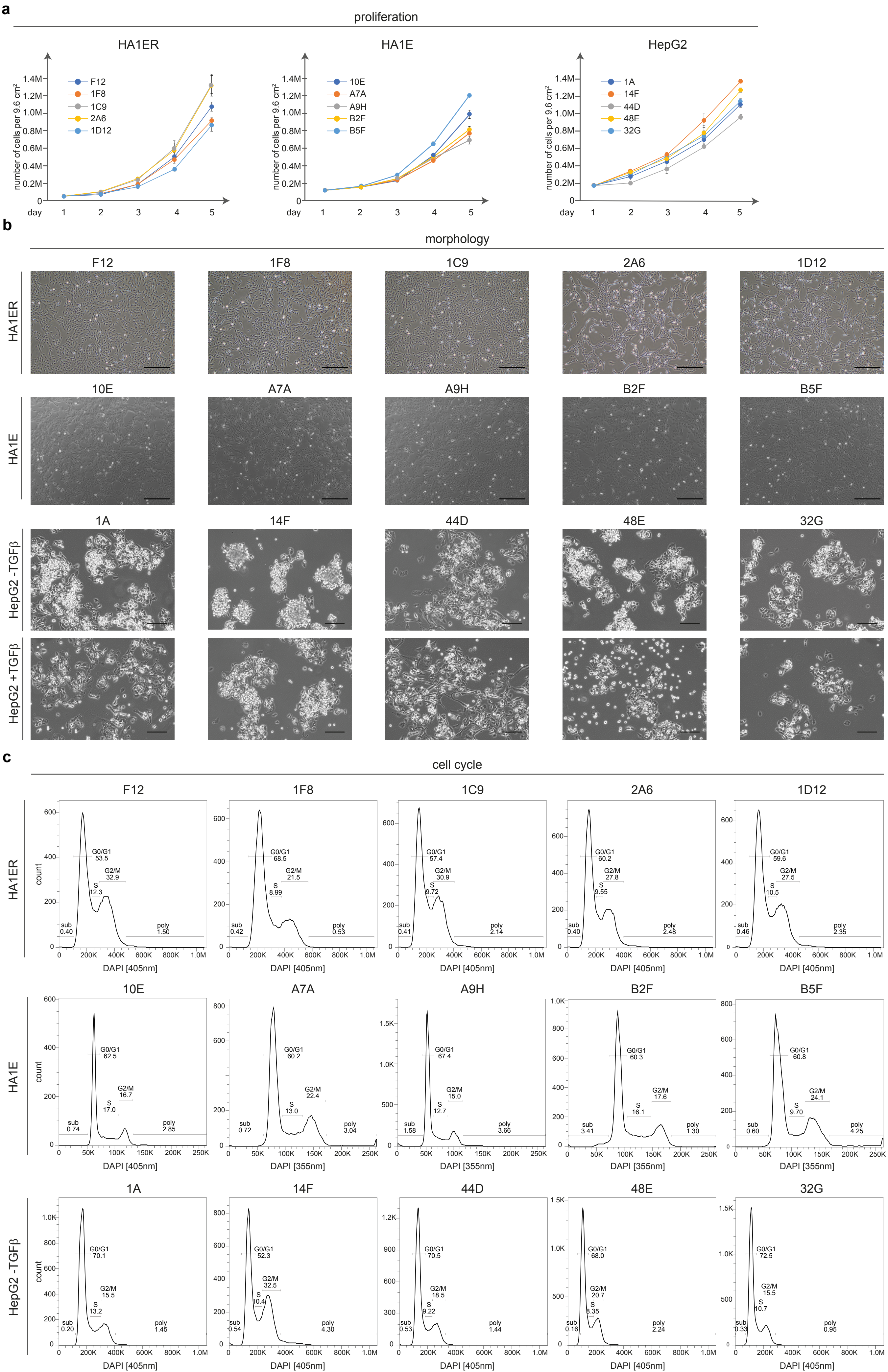
a
proliferation
HA1ER
F12
1F8
1C9
2A6
1D12
HA1E
10E
A7A
A9H
B2F
B5F
HepG2
1A
14F
44D
48E
32G
number of cells per 9.6 cm²
0
0.2M
0.4M
0.6M
0.8M
1.0M
1.2M
1.4M
day
1
2
3
4
5
b
morphology
HA1ER
F12
1F8
1C9
2A6
1D12
HA1E
10E
A7A
A9H
B2F
B5F
HepG2 -TGFβ
HepG2 +TGFβ
1A
14F
44D
48E
32G
c
cell cycle
HA1ER
F12
sub 0.40
G0/G1 53.5
S 12.3
G2/M 32.9
poly 1.50
1F8
sub 0.42
G0/G1 68.5
S 8.99
G2/M 21.5
poly 0.53
1C9
sub 0.41
G0/G1 57.4
S 9.72
G2/M 30.9
poly 2.14
2A6
sub 0.40
G0/G1 60.2
S 9.55
G2/M 27.8
poly 2.48
1D12
sub 0.46
G0/G1 59.6
S 10.5
G2/M 27.5
poly 2.35
count
DAPI [405nm]
HA1E
10E
sub 0.74
G0/G1 62.5
S 17.0
G2/M 16.7
poly 2.85
DAPI [405nm]
A7A
sub 0.72
G0/G1 60.2
S 13.0
G2/M 22.4
poly 3.04
DAPI [355nm]
A9H
sub 1.58
G0/G1 67.4
S 12.7
G2/M 15.0
poly 3.66
DAPI [405nm]
B2F
sub 3.41
G0/G1 60.3
S 16.1
G2/M 17.6
poly 1.30
DAPI [355nm]
B5F
sub 0.60
G0/G1 60.8
S 9.70
G2/M 24.1
poly 4.25
DAPI [355nm]
HepG2 -TGFβ
1A
sub 0.20
G0/G1 70.1
S 13.2
G2/M 15.5
poly 1.45
14F
sub 0.54
G0/G1 52.3
S 10.4
G2/M 32.5
poly 4.30
44D
sub 0.53
G0/G1 70.5
S 9.22
G2/M 18.5
poly 1.44
48E
sub 0.16
G0/G1 68.0
S 8.35
G2/M 20.7
poly 2.24
32G
sub 0.33
G0/G1 72.5
S 10.7
G2/M 15.5
poly 0.95
DAPI [405nm]

**Supplementary Fig. 1: HA1ER, HA1E and HepG2 diverging clonal systems. a.** Line graphs represent the number of cells per 9.6 $cm^2$ wells for HA1ER (F12, 1F8, 1C9, 2A6 and 1D12), HA1E (10E, A7A, A9H, B2F and B5F) and HepG2 (1A, 14F, 44D, 48E and 32G) clonal cell systems throughout 4 days in culture. Parental clones for each system are depicted in dark blue whilst diverging subclones are labelled in orange, grey, yellow and light blue. Clone identity is depicted. Data points represent the mean value ± standard deviation (SD) of 3 biological replicates. **b.** Cells from the HA1ER, HA1E and HepG2 (-TGFβ and +TGFβ) diverging clonal system were imaged by light microscopy. Panels from each parental clone (first images on the left) and diverging subclones are depicted. Scale bar represents 500 µm (HA1ER and HA1E) or 150 µm (HepG2). **c.** Cells from the HA1ER, HA1E and HepG2 clonal cell systems were subjected to cell cycle analysis using flow cytometry. Histograms show the representation of cells in each phase of the cell cycle as analysed by DAPI staining. Data from one representative experiment for each clone is shown. Percentage of cells in different phases of the cell cycle (G0/G1, S and G2/M), sub-G1 (sub) and polyploidy levels (poly) are depicted.

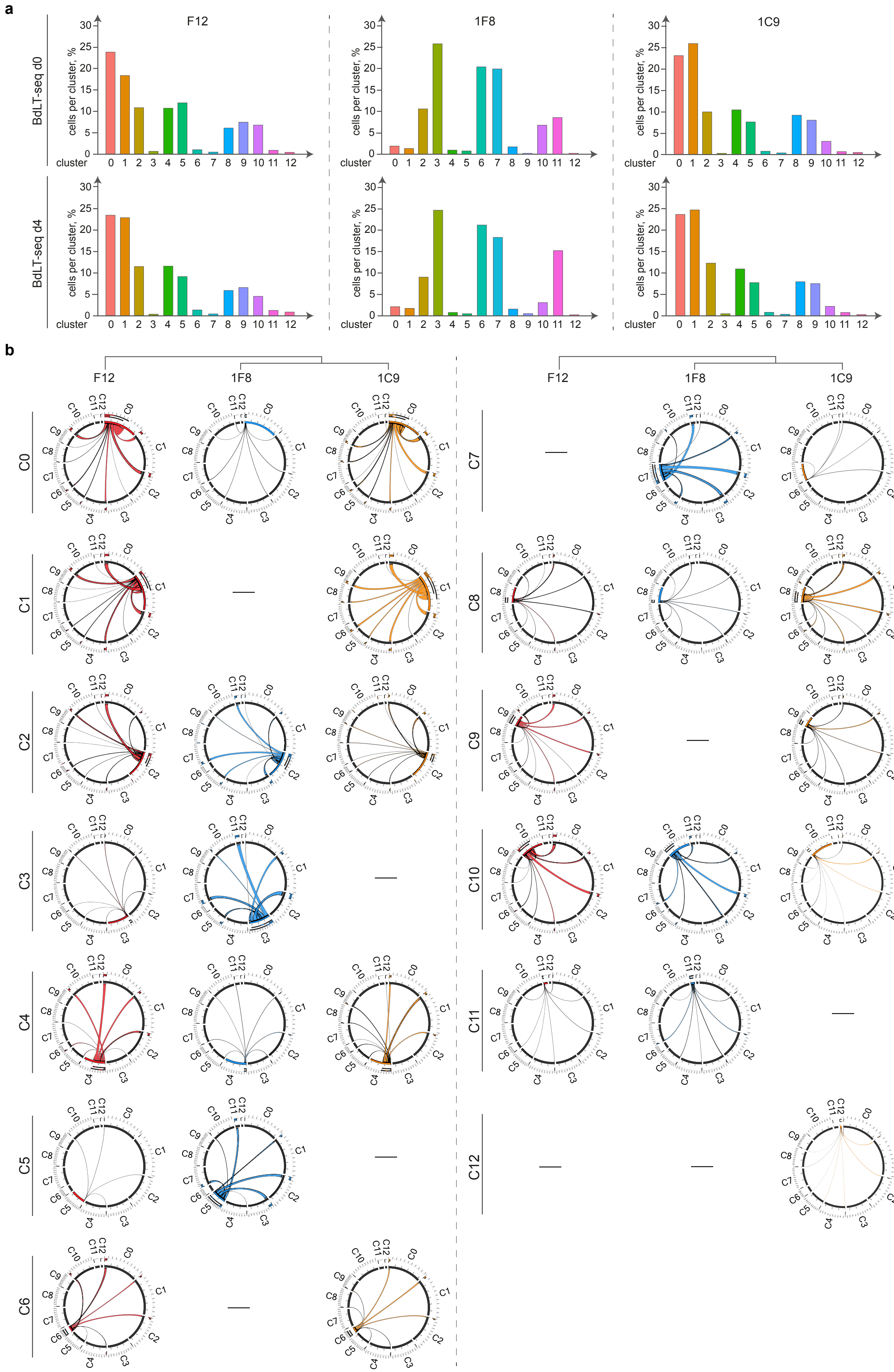
a
F12
1F8
1C9
BdLT-seq d0
BdLT-seq d4
cells per cluster, %
cluster
b
F12
1F8
1C9
C0
C1
C2
C3
C4
C5
C6
C7
C8
C9
C10
C11
C12

**Supplementary Fig. 2: Barcode Decay Lineage Tracing (BdLT-Seq) coupled to scRNA-Seq on HA1ER clones F12, 1F8 and 1C9. a.** Histograms represent the percentage of cells for each clone (F12, 1F8 and 1C9) that belong to individual identified gene expression clusters at day 0 (BdLT-Seq d0) or day 4 (BdLT-Seq d4) of tracing shown in Fig. 1a. **b.** Chord diagrams represent transcriptome state dynamics for cells belonging to a particular state/cluster at the beginning of tracing and their divergence after 4 days for the parental clone F12 and subclones 1F8 and 1C9. Detected clusters are depicted (C0 to C12) and integrate the collapsed behaviour of all cells that belong to each gene expression state. Origin clusters are depicted on the left (Day 0) and chords represent the endpoint cluster association (Day 4).

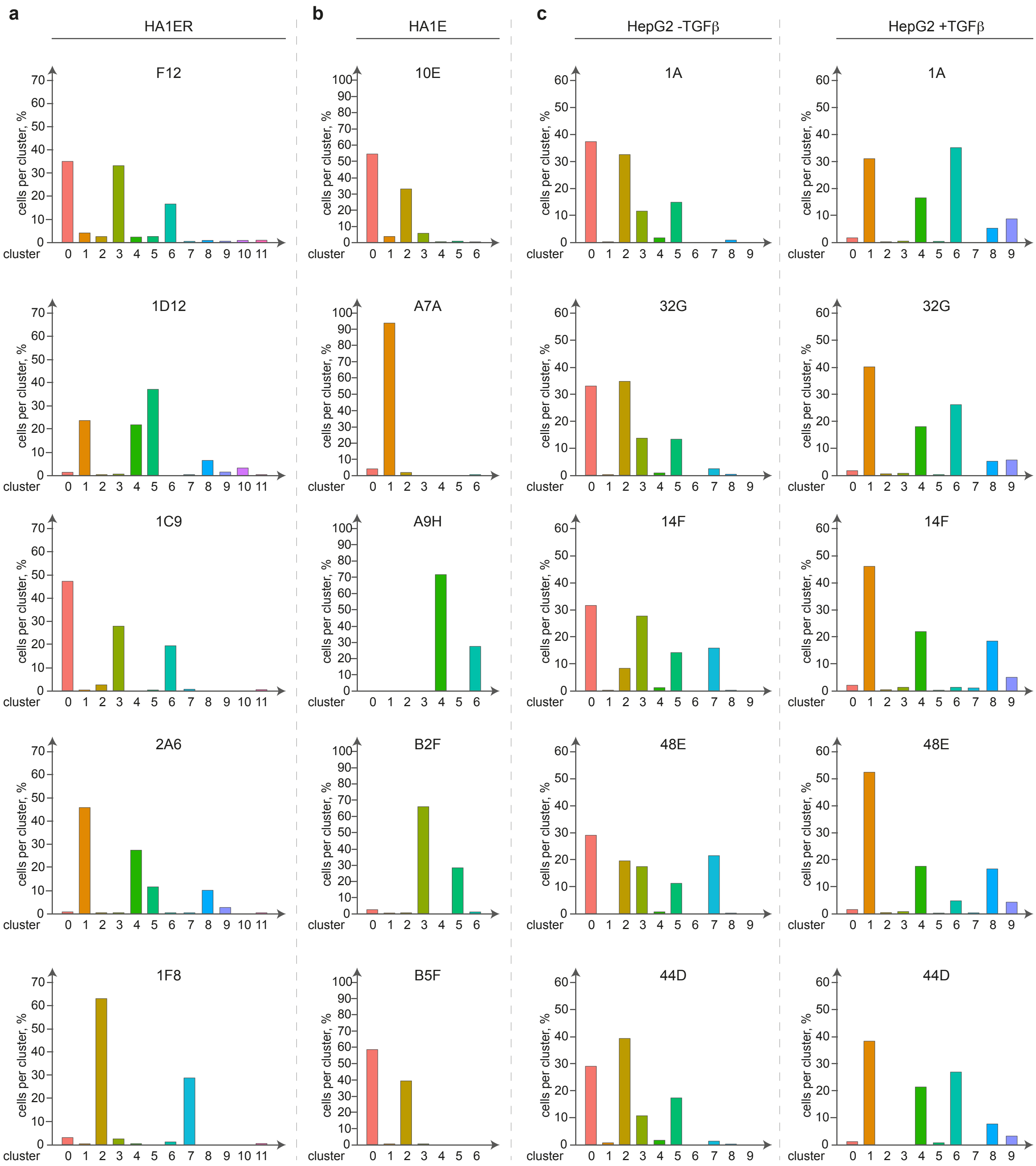
a
HA1ER
F12
1D12
1C9
2A6
1F8
b
HA1E
10E
A7A
A9H
B2F
B5F
c
HepG2 -TGFβ
HepG2 +TGFβ
1A
32G
14F
48E
44D
cells per cluster, %
cluster

**Supplementary Fig. 3: HA1ER, HA1E and HepG2 clonal systems cluster representation. a.** Cells from the diverging clonal system HA1ER (F12, 1F8, 1C9, 2A6 and 1D12) were subjected to scRNA-Seq. Histograms represents percentage of cells in each cluster for each clone. **b.** Cells from the diverging clonal system HA1E (10E, A7A, A9H, B2F and B5F) were subjected to scRNA-Seq. Histograms represents percentage of cells in each cluster for each clone. **c.** Cells from the diverging clonal system HepG2 (1A, 14F, 44D, 48E and 32G) were subjected to scRNA-Seq. Histograms represents percentage of cells in each cluster for each clone. HepG2 clonal system data from either vehicle (HepG2 -TGFβ) or TGFβ treated cells (HepG2 +TGFβ) are depicted.

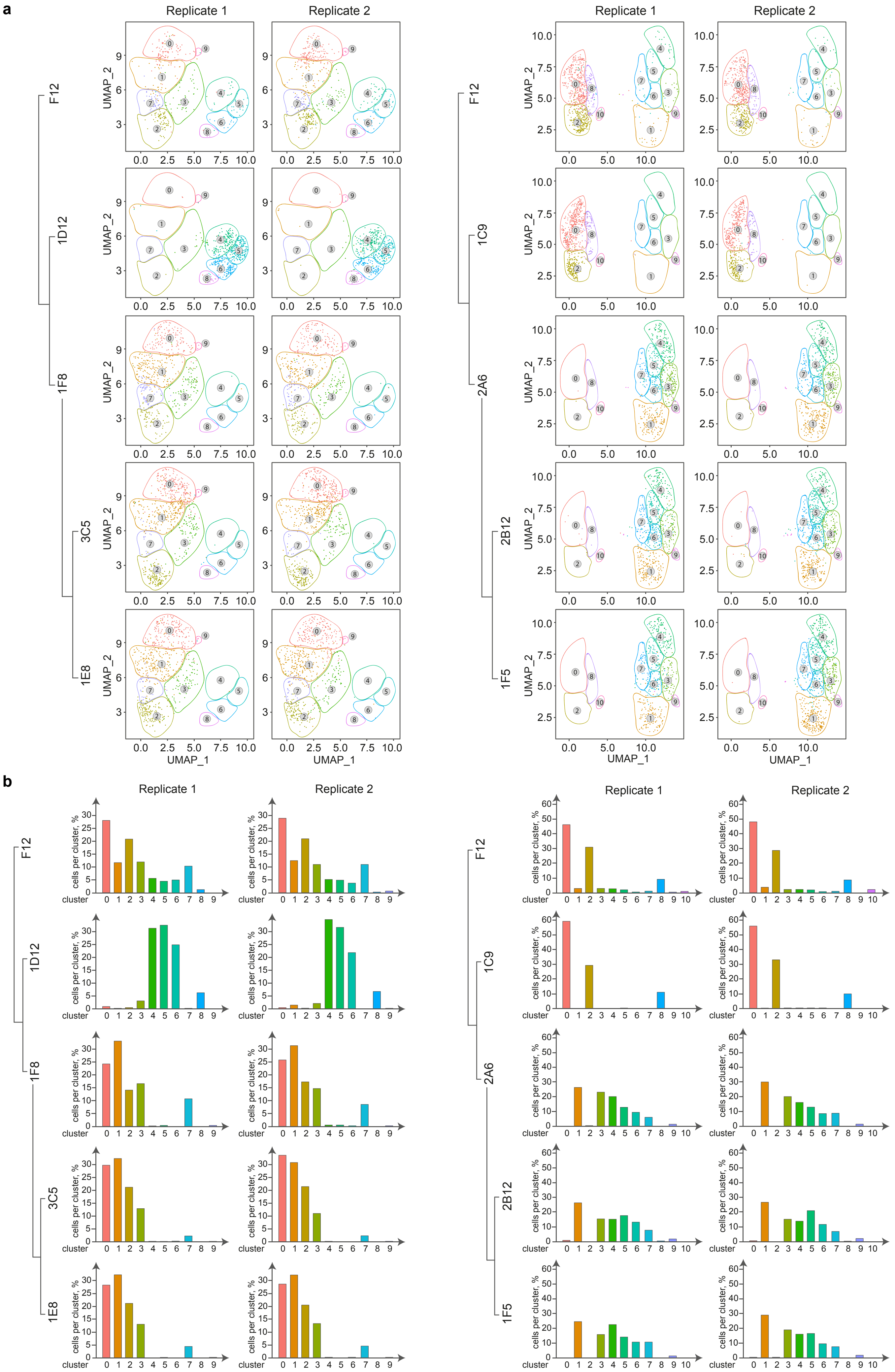
a
Replicate 1
Replicate 2
F12
1D12
1F8
3C5
1E8
1C9
2A6
2B12
1F5
UMAP_1
UMAP_2
b
cells per cluster, %
cluster

**Supplementary Fig. 4: HA1ER diverging clones propagate transcriptomic states further down the lineage. a.** Cells from the HA1ER diverging clonal system (1F8 and 2A6) were further subjected to subcloning (1F8 subclones 3C5 and 1E8; 2A6 subclones 2B12 and 1F5) and analysed by scRNA-Seq in parallel to parental clone F12 and subclones 1D12 and 1C9. UMAP plots depict the transcriptomic states displayed by each clone analysed. Biological replicates are depicted. Lineage relationship is shown. **b.** Cells from the diverging clonal system HA1ER (F12, 1D12, 1F8, 3C5, 1E8, 2A6, 1C9, 2B12 and 1F5) were subjected to scRNA-Seq. Histograms represents percentage of cells per cluster for each clone. Biological replicates are depicted. Lineage relationship is shown.

HA1ER WES

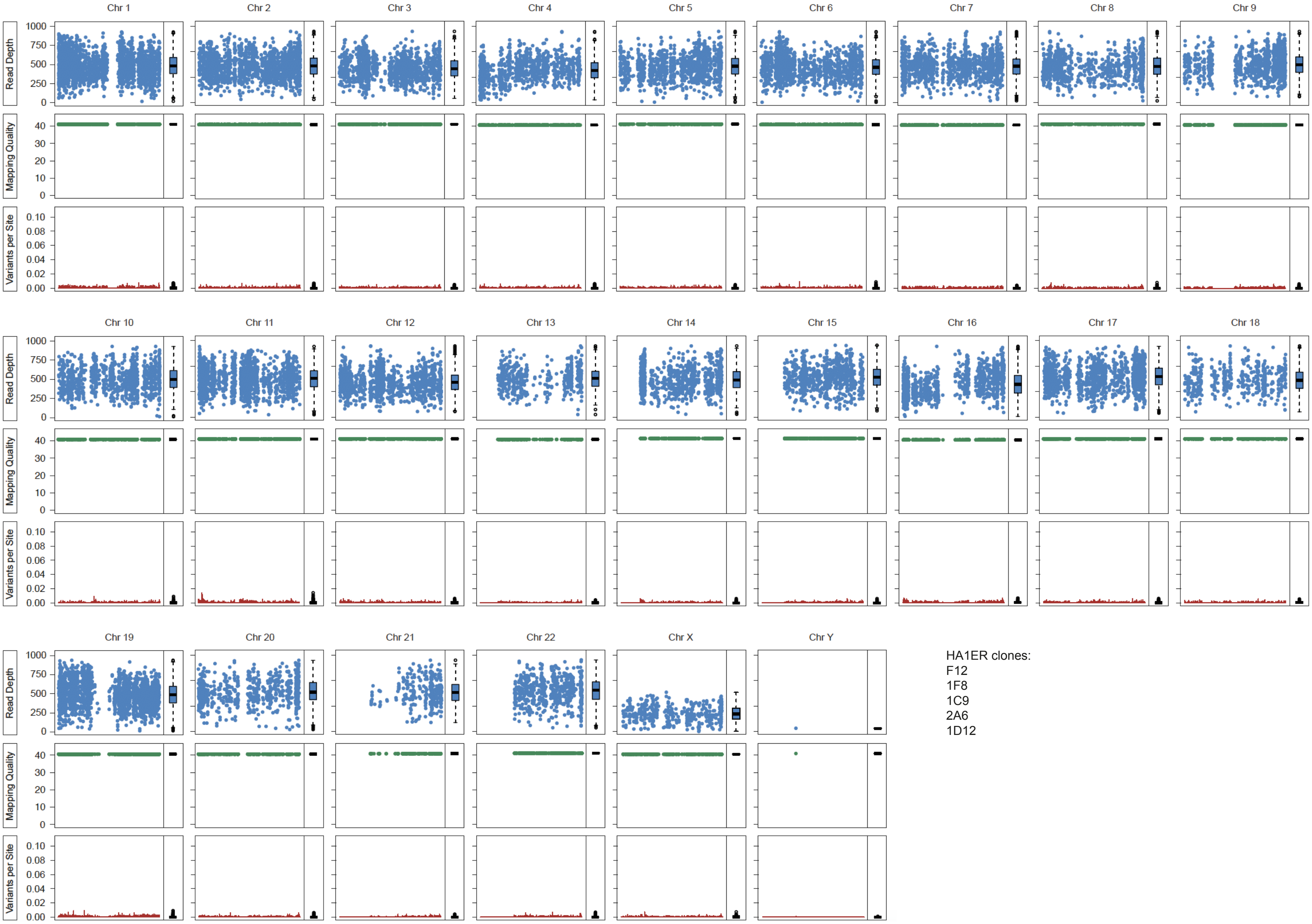
Chr 1
Chr 2
Chr 3
Chr 4
Chr 5
Chr 6
Chr 7
Chr 8
Chr 9
Chr 10
Chr 11
Chr 12
Chr 13
Chr 14
Chr 15
Chr 16
Chr 17
Chr 18
Chr 19
Chr 20
Chr 21
Chr 22
Chr X
Chr Y
Read Depth
Mapping Quality
Variants per Site
1000
750
500
250
0
40
30
20
10
0
0.10
0.08
0.06
0.04
0.02
0.00
HA1ER clones:
F12
1F8
1C9
2A6
1D12

HA1E WES

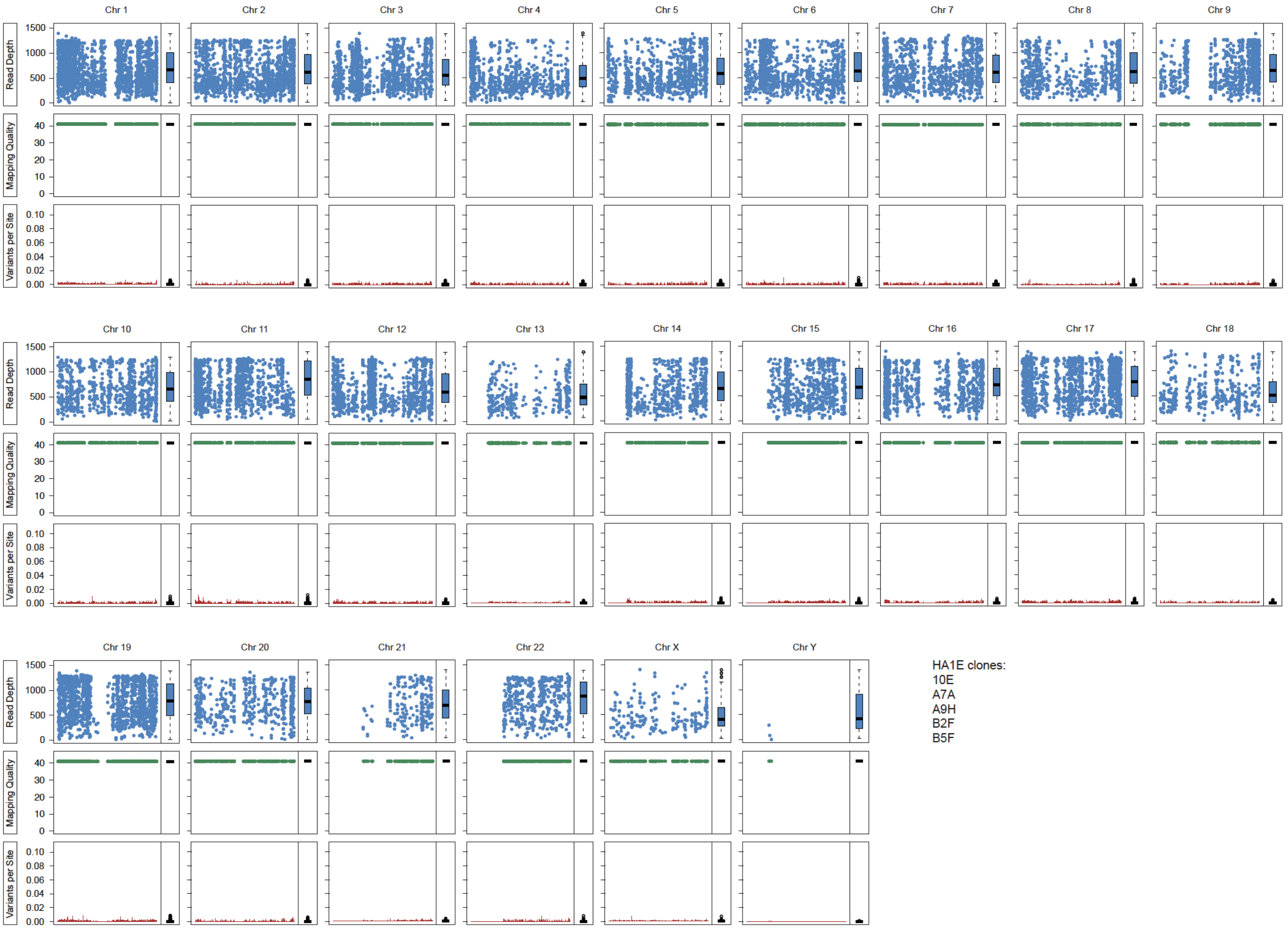
Chr 1
Chr 2
Chr 3
Chr 4
Chr 5
Chr 6
Chr 7
Chr 8
Chr 9
Chr 10
Chr 11
Chr 12
Chr 13
Chr 14
Chr 15
Chr 16
Chr 17
Chr 18
Chr 19
Chr 20
Chr 21
Chr 22
Chr X
Chr Y
Read Depth
Mapping Quality
Variants per Site
1500
1000
500
0
40
30
20
10
0
0.10
0.08
0.06
0.04
0.02
0.00
HA1E clones:
10E
A7A
A9H
B2F
B5F

HepG2 WES

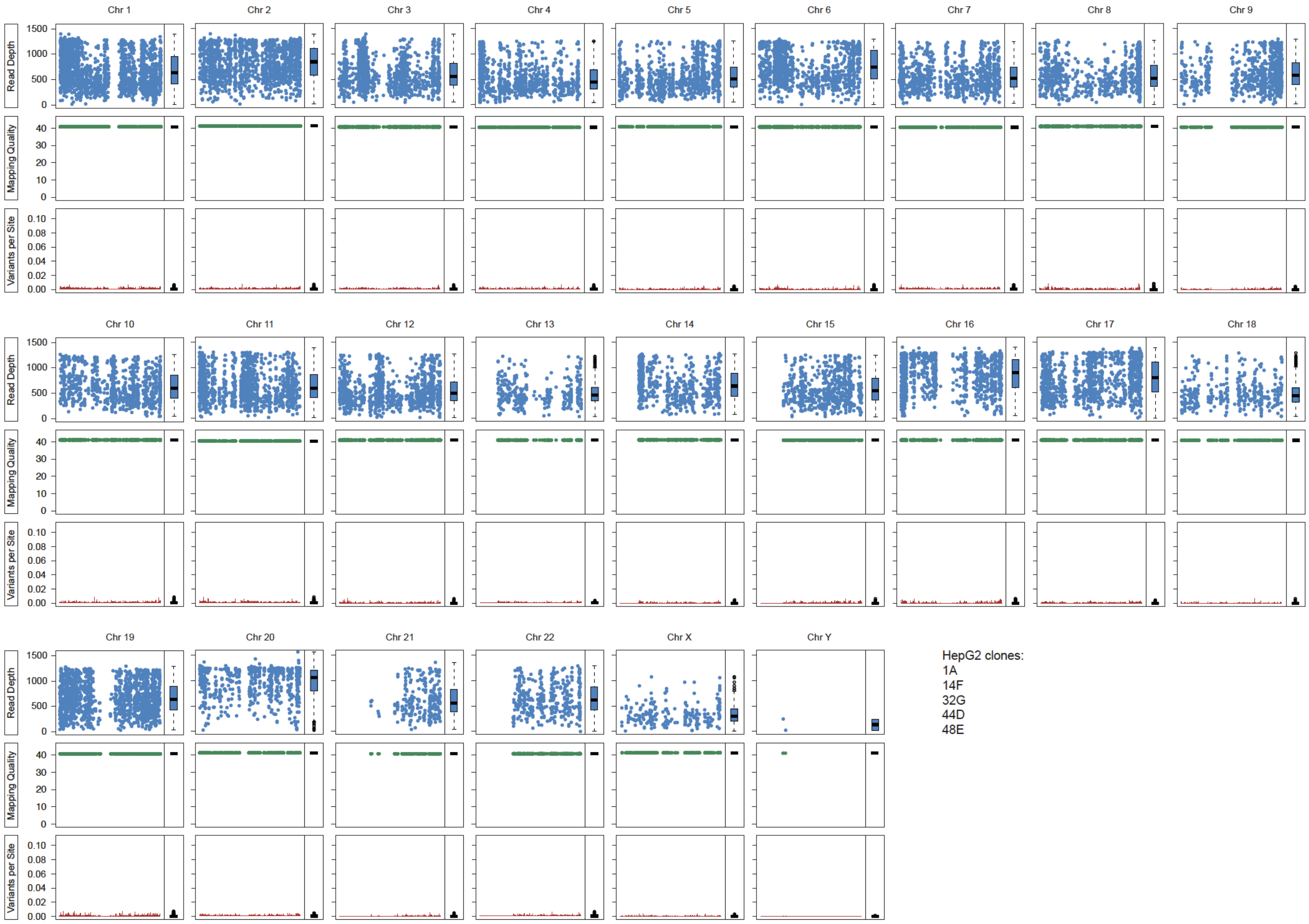
Chr 1
Chr 2
Chr 3
Chr 4
Chr 5
Chr 6
Chr 7
Chr 8
Chr 9
Chr 10
Chr 11
Chr 12
Chr 13
Chr 14
Chr 15
Chr 16
Chr 17
Chr 18
Chr 19
Chr 20
Chr 21
Chr 22
Chr X
Chr Y
Read Depth
0
500
1000
1500
Mapping Quality
0
10
20
30
40
Variants per Site
0.00
0.02
0.04
0.06
0.08
0.10
HepG2 clones:
1A
14F
32G
44D
48E

**Supplementary Fig. 5: HA1ER, HA1E and HepG2 diverging clonal cells systems do not show genetic variation within the exome.** Whole exome sequencing (WES; 100X coverage) scatter plots depicting read depth, mapping quality and identified variants per-site are displayed. Data is shown by chromosome and for all clones from the same cellular system (HA1ER, HA1E, HepG2). Box-plots summarising each data level are depicted.

HA1ER WGS

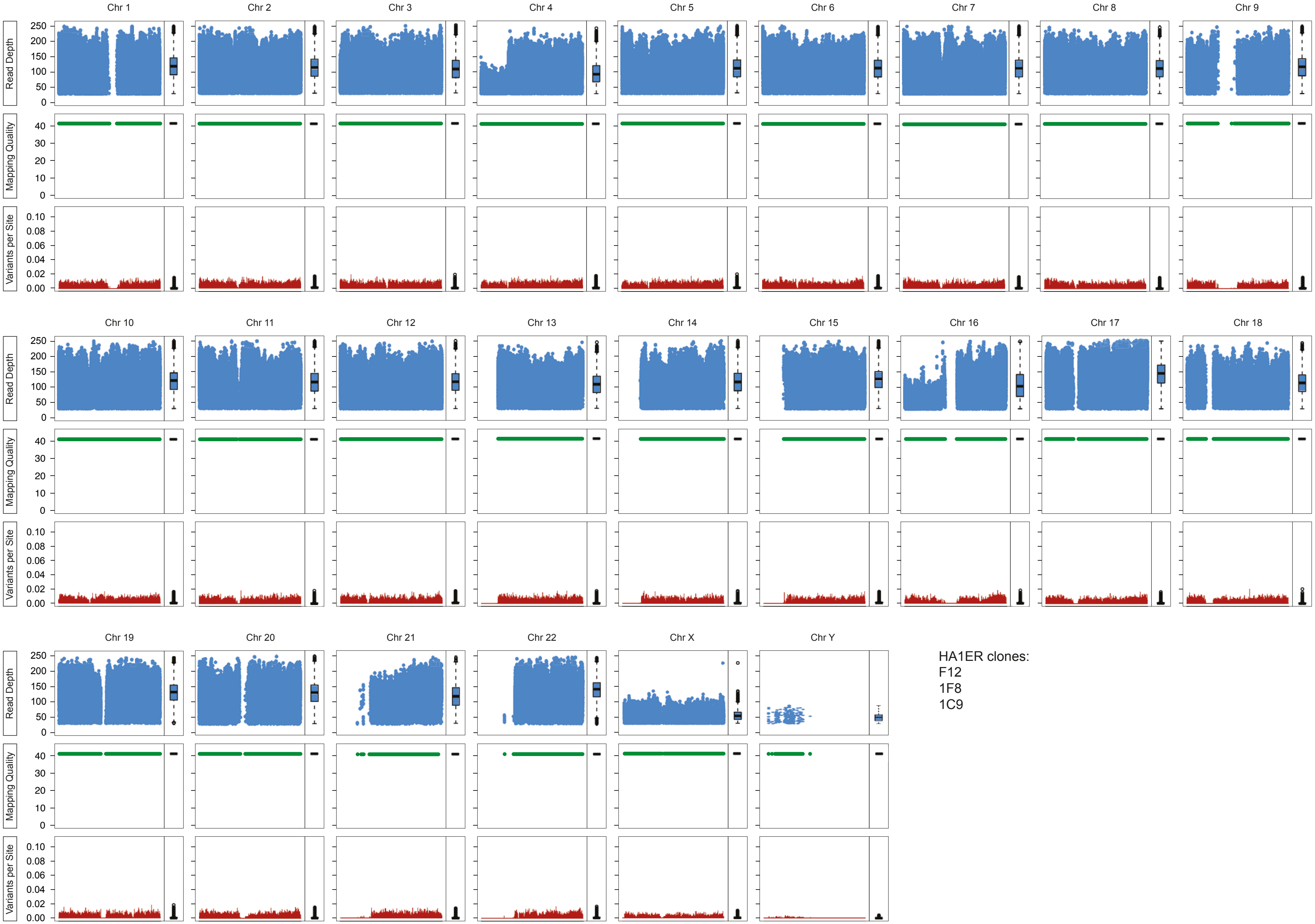
Chr 1
Chr 2
Chr 3
Chr 4
Chr 5
Chr 6
Chr 7
Chr 8
Chr 9
Chr 10
Chr 11
Chr 12
Chr 13
Chr 14
Chr 15
Chr 16
Chr 17
Chr 18
Chr 19
Chr 20
Chr 21
Chr 22
Chr X
Chr Y
Read Depth
Mapping Quality
Variants per Site
0
50
100
150
200
250
0
10
20
30
40
0.00
0.02
0.04
0.06
0.08
0.10
HA1ER clones:
F12
1F8
1C9

HepG2 WGS

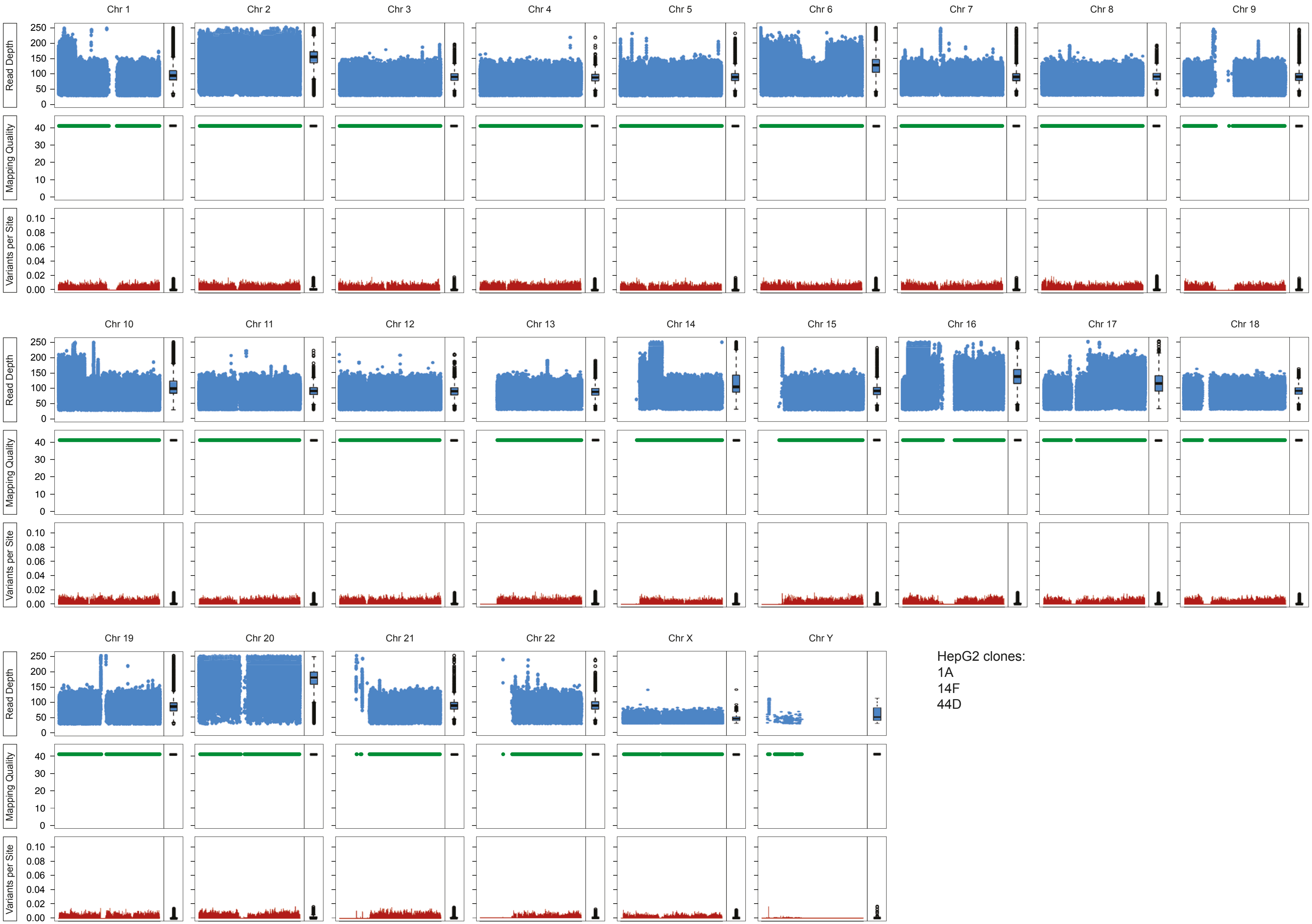
Chr 1
Chr 2
Chr 3
Chr 4
Chr 5
Chr 6
Chr 7
Chr 8
Chr 9
Chr 10
Chr 11
Chr 12
Chr 13
Chr 14
Chr 15
Chr 16
Chr 17
Chr 18
Chr 19
Chr 20
Chr 21
Chr 22
Chr X
Chr Y
Read Depth
Mapping Quality
Variants per Site
0
50
100
150
200
250
10
20
30
40
0.00
0.02
0.04
0.06
0.08
0.10
HepG2 clones:
1A
14F
44D

**Supplementary Fig. 6: No genetic variation was detected for HA1ER and HepG2 diverging clones.** Scatter plots of whole genome sequencing data (WGS; 30X coverage) representing read depth, mapping quality and identified variants per-site. Data is shown by chromosome for HA1ER clones F12, 1F8 and 1C9 and separately for HepG2 clones 1A, 14F and 44D. Box-plots summarising each data level are shown.

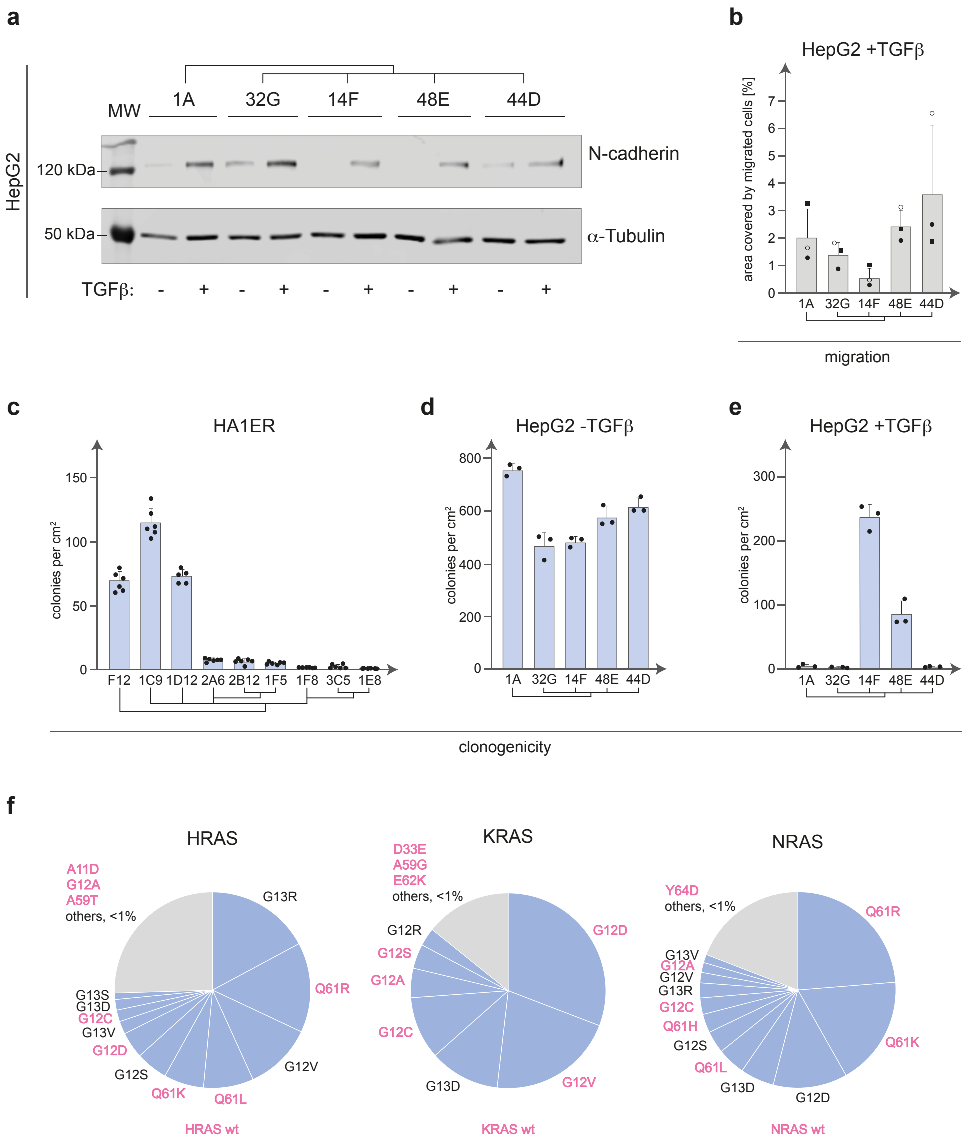


**Supplementary Fig. 7: Phenotypic output varies among diverging clones. a.** Clonal cell populations from the HepG2 clonal system (1A, 14F, 44D, 48E and 32G) were subjected to either vehicle (-TGFβ) or TGFβ treatment (+TGFβ) and processed for Western blot. Staining with antibodies against N-cadherin to monitor epithelial to mesenchymal transition (EMT) and against α-Tubulin as a loading control. Molecular weight markers are indicated. **b.** Clonal cell populations from the HepG2 clonal diverging system (1A, 14F, 44D, 48E and 32G) treated with TGFβ were assessed for their capability to migrate in transwell assays. Histograms depict the mean area covered by migrated cells ± standard deviation (SD) of 3 independent experiments performed in triplicate. Individual data points are shown. Clonal relationship is depicted. **c.** Clonogenicity was analysed for all HA1ER clones depicted in Supplementary Fig. 4 (F12, 1D12, 1F8, 3C5, 1E8, 2A6, 1C9, 2B12 and 1F5). The number of colonies per cm$^2$ is depicted. Individual data points are depicted and represent the mean value ± standard deviation (SD) of six replicates. Lineage relationships are depicted. **d.** Clonogenicity was analysed for HepG2 diverging clones (1A, 14F, 44D, 48E and 32G) in -TGFβ conditions. The number of colonies per cm$^2$ is depicted. Individual data points are shown and represent the mean value ± standard deviation (SD) of 3 replicates. Lineage relationships are depicted. **e.** Clonogenicity was analysed for HepG2 diverging clones (1A, 14F, 44D, 48E and 32G) in +TGFβ conditions. The number of colonies per cm$^2$ is depicted. Individual data points are shown and represent the mean value ± standard deviation (SD) of 3 replicates. Lineage relationships are depicted. **f.** Pie charts representing the pan-cancer frequency of RAS variants from the three main families (H-, K and N-) found in COSMIC database (v92[92]). RAS mutations accounting for >1% for each RAS family are annotated. RAS variants used to build the multiplexed library used in Fig. 2e to evaluate clonogenicity of HA1E clones are depicted in pink.

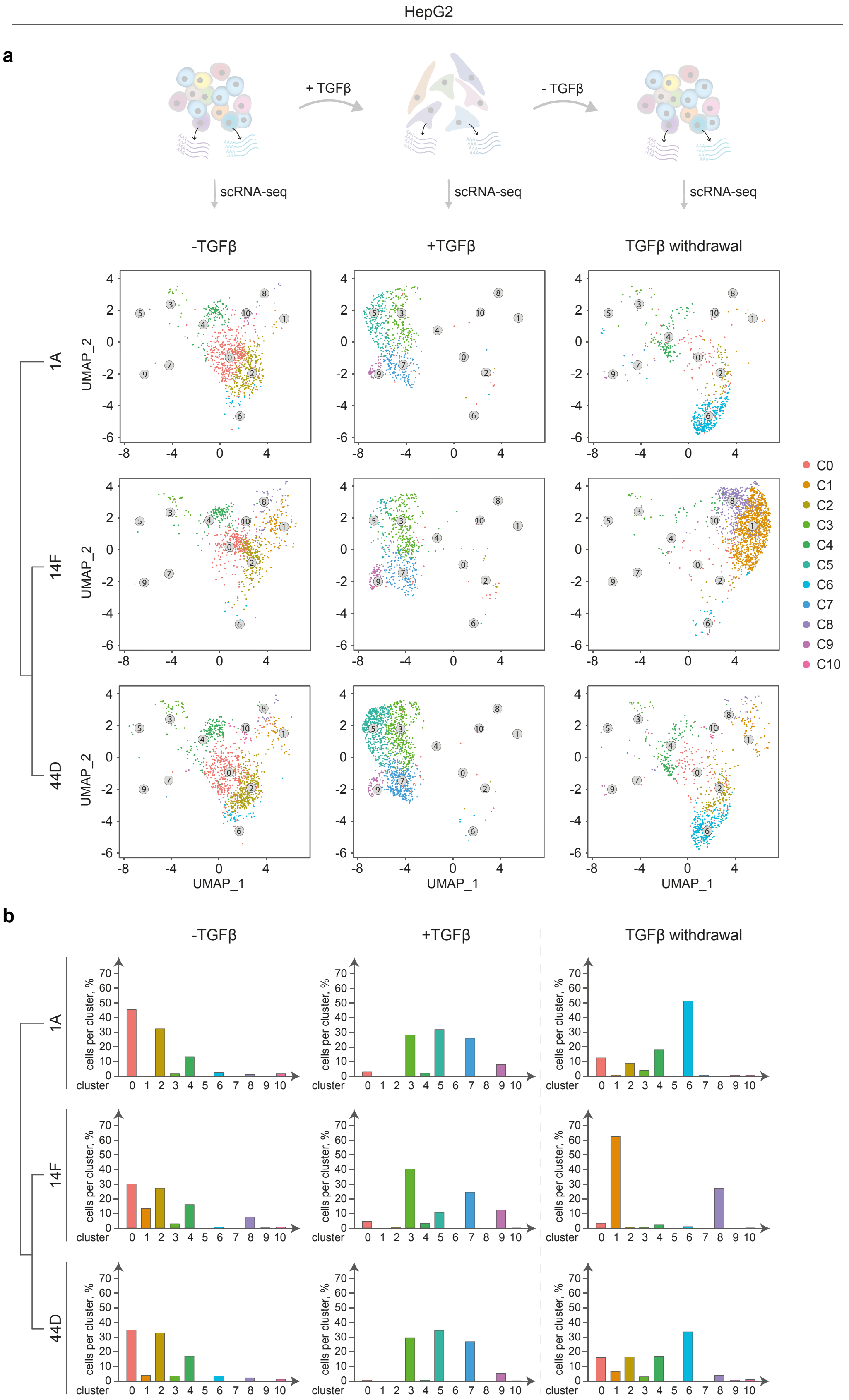


**Supplementary Fig. 8: HepG2 diverging clones retain a molecular memory of past states even when a transcriptional programme is enforced. a.** Cells from the HepG2 clonal system (1A, 4F and 44D) were subjected to either vehicle (-TGFβ) or TGFβ treatment (+TGFβ) for 4 days and then TGFβ was withdrawn (TGFβ withdrawal) for additional 4 days. Samples were then processed for scRNA-Seq. UMAP plots represent the identified transcriptomic states for each clone analysed and subjected to the EMT (+TGFβ) and MET (TGFβ withdrawal) conditions. Lineage relationships are depicted. **b.** Histograms represents the percentage of cells in each cluster for each clone analysed in **a**. Lineage relationship is shown.

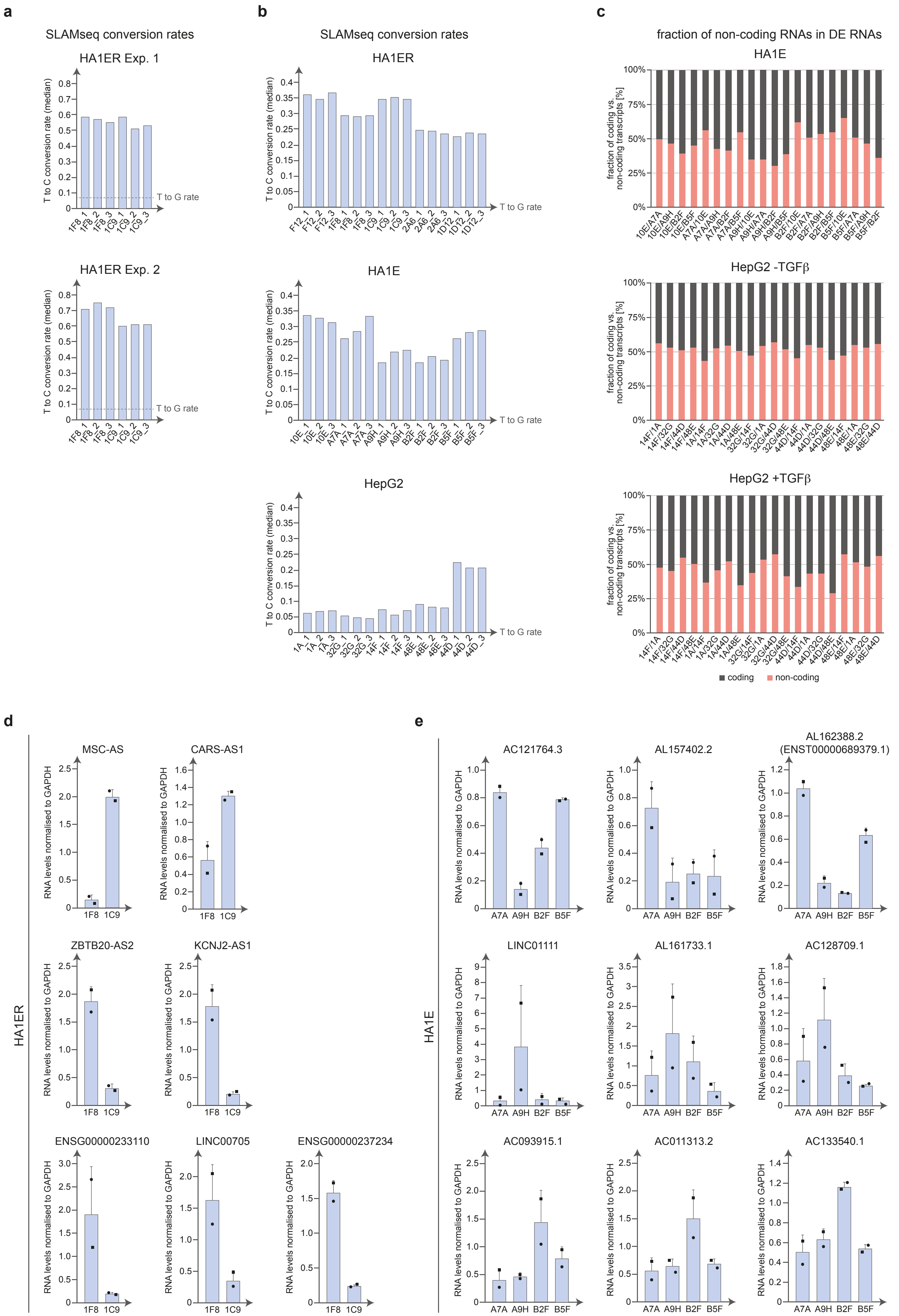

a
SLAMseq conversion rates
HA1ER Exp. 1
HA1ER Exp. 2
T to C conversion rate (median)
T to G rate
b
SLAMseq conversion rates
HA1ER
HA1E
HepG2
c
fraction of non-coding RNAs in DE RNAs
HA1E
HepG2 -TGFβ
HepG2 +TGFβ
fraction of coding vs. non-coding transcripts [%]
coding
non-coding
d
HA1ER
MSC-AS
CARS-AS1
ZBTB20-AS2
KCNJ2-AS1
ENSG00000233110
LINC00705
ENSG00000237234
RNA levels normalised to GAPDH
1F8
1C9
e
HA1E
AC121764.3
AL157402.2
AL162388.2 (ENST00000689379.1)
LINC01111
AL161733.1
AC128709.1
AC093915.1
AC011313.2
AC133540.1
A7A
A9H
B2F
B5F

f

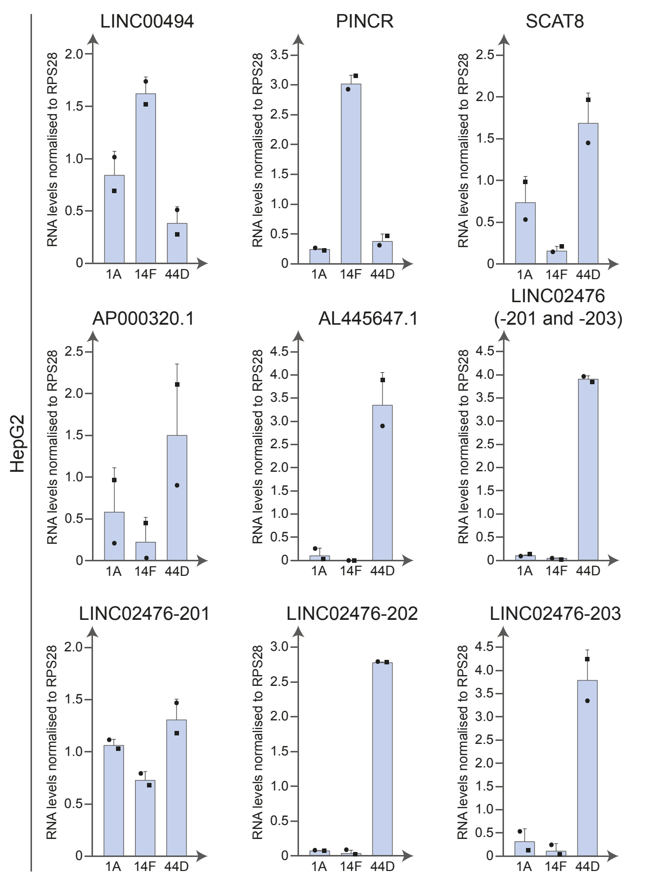


g

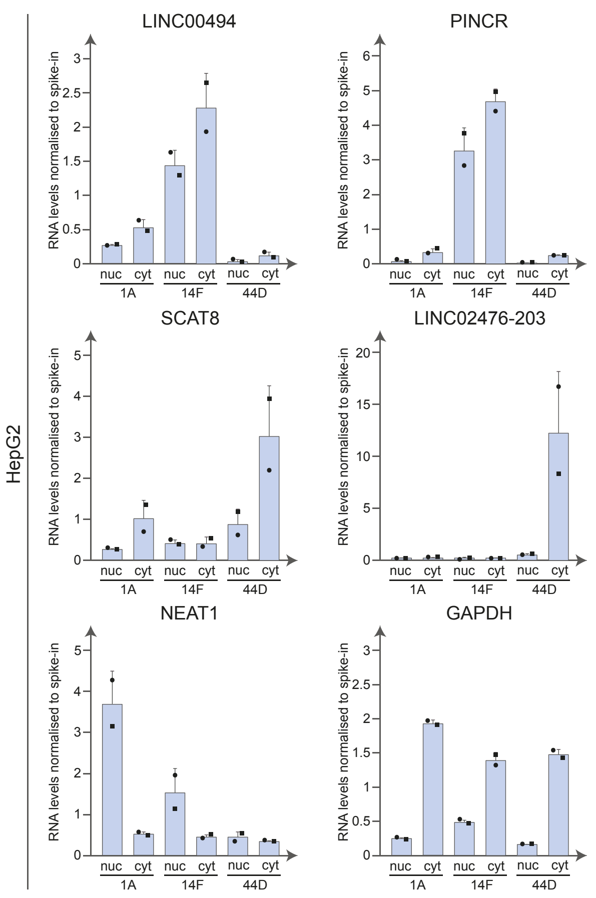


**Supplementary Fig. 9: SLAM-Seq quality control, coding vs non-coding RNA annotation and validation of diverging lncRNAs. a.** SLAM-Seq analysis was performed in HA1ER clones 1F8 and 1C9. Histograms depict the median conversion rate of T to C transition. T to G conversion rate is shown as a baseline. SLAM-Seq analysis was performed in 2 independent experiments (Exp.1 and Exp. 2) performed in biological triplicates. **b.** Cells from the HA1ER (F12, 1F8, 1C9, 2A6 and 1D12), HA1E (10E, A7A, A9H, B2F and B5F) and HepG2 (1A, 14F, 44D, 48E and 32G) clonal cell systems were subjected to SLAM-Seq analysis. Histograms depict the median conversion rate of T to C transition. T to G conversion rate is shown as a baseline. SLAM-Seq analysis was performed in 3 biological replicates for each clone analysed. **c.** Cells from the HA1E (10E, A7A, A9H, B2F and B5F) and HepG2 (1A, 14F, 44D, 48E and 32G; with (+TGFβ) and without (-TGFβ) TGFβ treatment) clonal cell systems were subjected to scRNA-Seq analysis (see Fig. 2a-c) Differentially expressed transcripts between clones (FC>3 up-regulated) were determined in a pairwise manner and annotated based on their coding vs. non-coding potential. **d.** Additional identified differentially expressed lncRNA transcripts between 1F8 vs 1C9 (see Fig. 3c, f) were validated by reverse transcription followed by qPCR (RT-qPCR). Histograms depict MSC-AS, CARS-AS1, ZBTB20-AS2, KCNJ2-AS1, ENSG00000233110, LINC00705 and ENSG00000237234 transcript levels normalised to GAPDH levels. Results represent the mean ± standard deviation (SD) of 2 independent experiments performed in technical triplicates. Individual data points are depicted. **e.** A subset of identified differentially expressed lncRNA transcripts between HA1E clones (A7A, A9H, B2F and B5F) were validated by reverse transcription followed by qPCR (RT-qPCR). Histograms depict AC121764.3, AL157402.2, AL162388.2 (ENST00000689379.1), AL161733.1, AL161733.1, AC128709.1, AC093915.1, AC011313.2 and AC133540.1 transcript levels normalised to GAPDH levels. Results represent the mean ± standard deviation (SD) of 2 independent experiments performed in technical triplicates. Individual data points are depicted. **f.** A subset of identified differentially expressed lncRNA transcripts between HepG2 clones (1A, 14F and 44D) were validated by reverse transcription followed by qPCR (RT-qPCR). Histograms depict LINC00494, PINCR, SCAT8, AP000320.1, AL445647.1 and LINC02476 (either transcript variants 201 + 203 or each transcript variant individually) transcript levels normalised to RPS28 levels. Results represent the mean ± standard deviation (SD) of 2 independent experiments performed in technical triplicates. Individual data points are depicted. **g.** A subset of differentially expressed lncRNA transcripts between HepG2 clones 1A, 14F and 44D were analysed for their intracellular localisation by subcellular fractionation (nucleus (nuc) vs cytoplasm(cyt)) and their levels evaluated by RT-qPCR. Histograms depict RNA levels of each fraction upon normalisation to the spike-in (custom plasmid). Data for LINC00494, PINCR, SCAT8 and LIONC02476-203 are depicted. NEAT1 (enriched in nucleus), GAPDH (enriched in cytoplasm) are shown as fractionation controls. Data represents the mean ± standard deviation (SD) of 2 independent experiments performed in technical triplicates. Individual data points are depicted.

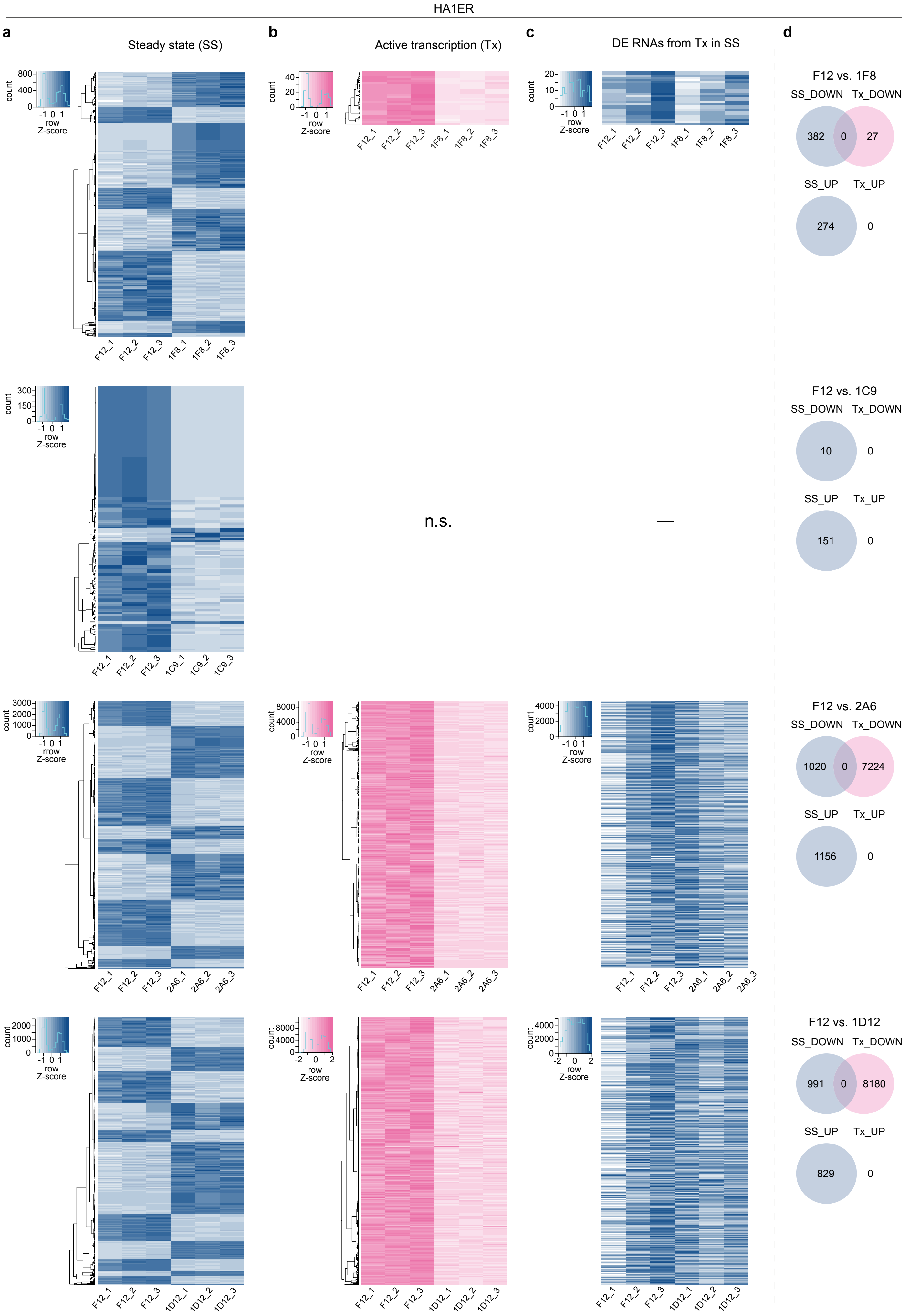
HA1ER
a
Steady state (SS)
b
Active transcription (Tx)
c
DE RNAs from Tx in SS
d
F12 vs. 1F8
SS_DOWN
Tx_DOWN
382
0
27
SS_UP
Tx_UP
274
0
F12 vs. 1C9
SS_DOWN
Tx_DOWN
10
0
SS_UP
Tx_UP
151
0
n.s.
—
F12 vs. 2A6
SS_DOWN
Tx_DOWN
1020
0
7224
SS_UP
Tx_UP
1156
0
F12 vs. 1D12
SS_DOWN
Tx_DOWN
991
0
8180
SS_UP
Tx_UP
829
0
count
row
Z-score
rrow
F12_1
F12_2
F12_3
1F8_1
1F8_2
1F8_3
1C9_1
1C9_2
1C9_3
2A6_1
2A6_2
2A6_3
1D12_1
1D12_2
1D12_3

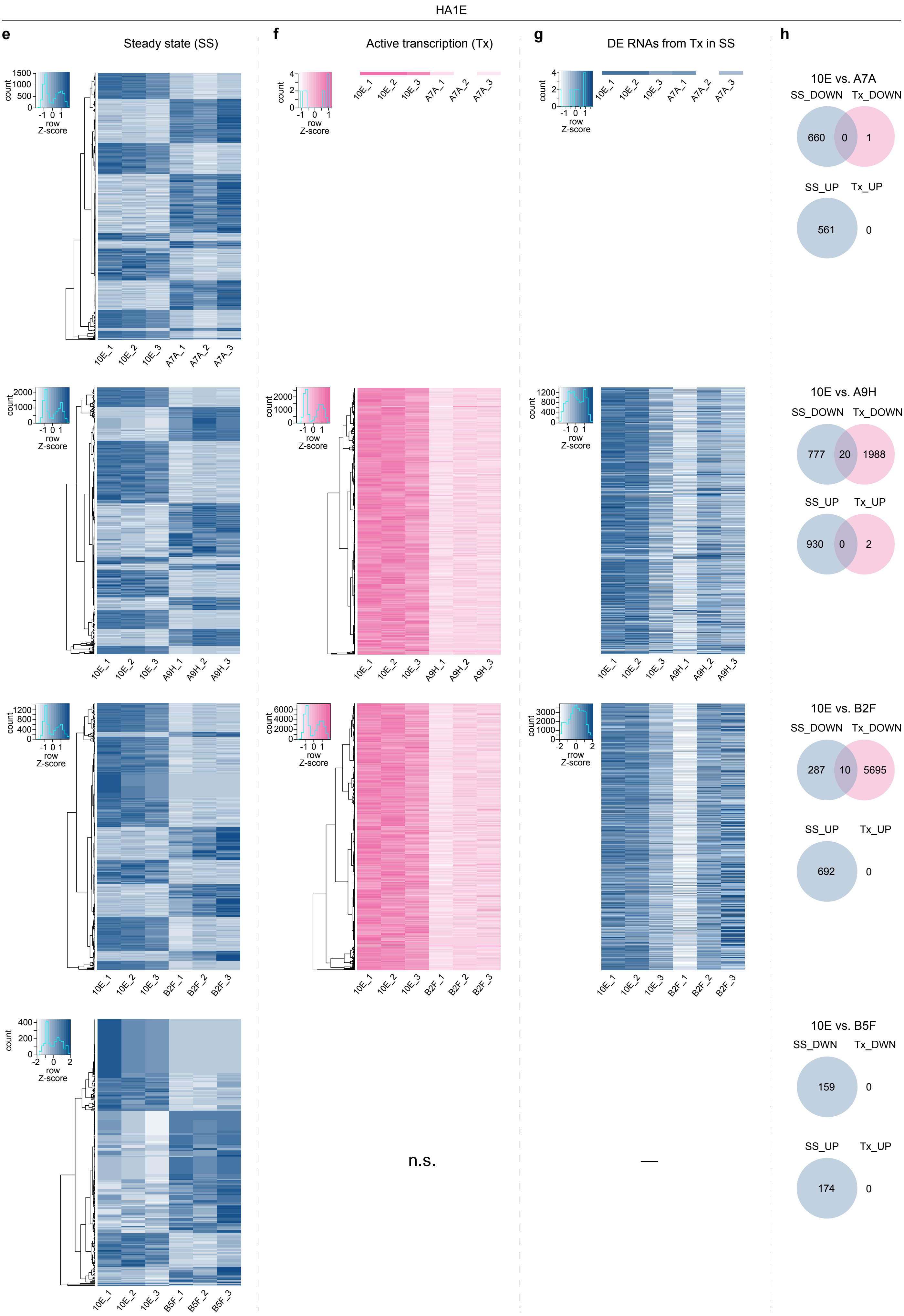
HA1E
e
Steady state (SS)
f
Active transcription (Tx)
g
DE RNAs from Tx in SS
h
count
row Z-score
10E_1
10E_2
10E_3
A7A_1
A7A_2
A7A_3
A9H_1
A9H_2
A9H_3
B2F_1
B2F_2
B2F_3
B5F_1
B5F_2
B5F_3
10E vs. A7A
SS_DOWN
Tx_DOWN
660
0
1
SS_UP
Tx_UP
561
0
10E vs. A9H
SS_DOWN
Tx_DOWN
777
20
1988
SS_UP
Tx_UP
930
0
2
10E vs. B2F
SS_DOWN
Tx_DOWN
287
10
5695
SS_UP
Tx_UP
692
0
10E vs. B5F
SS_DWN
Tx_DWN
159
0
SS_UP
Tx_UP
174
0
n.s.
—

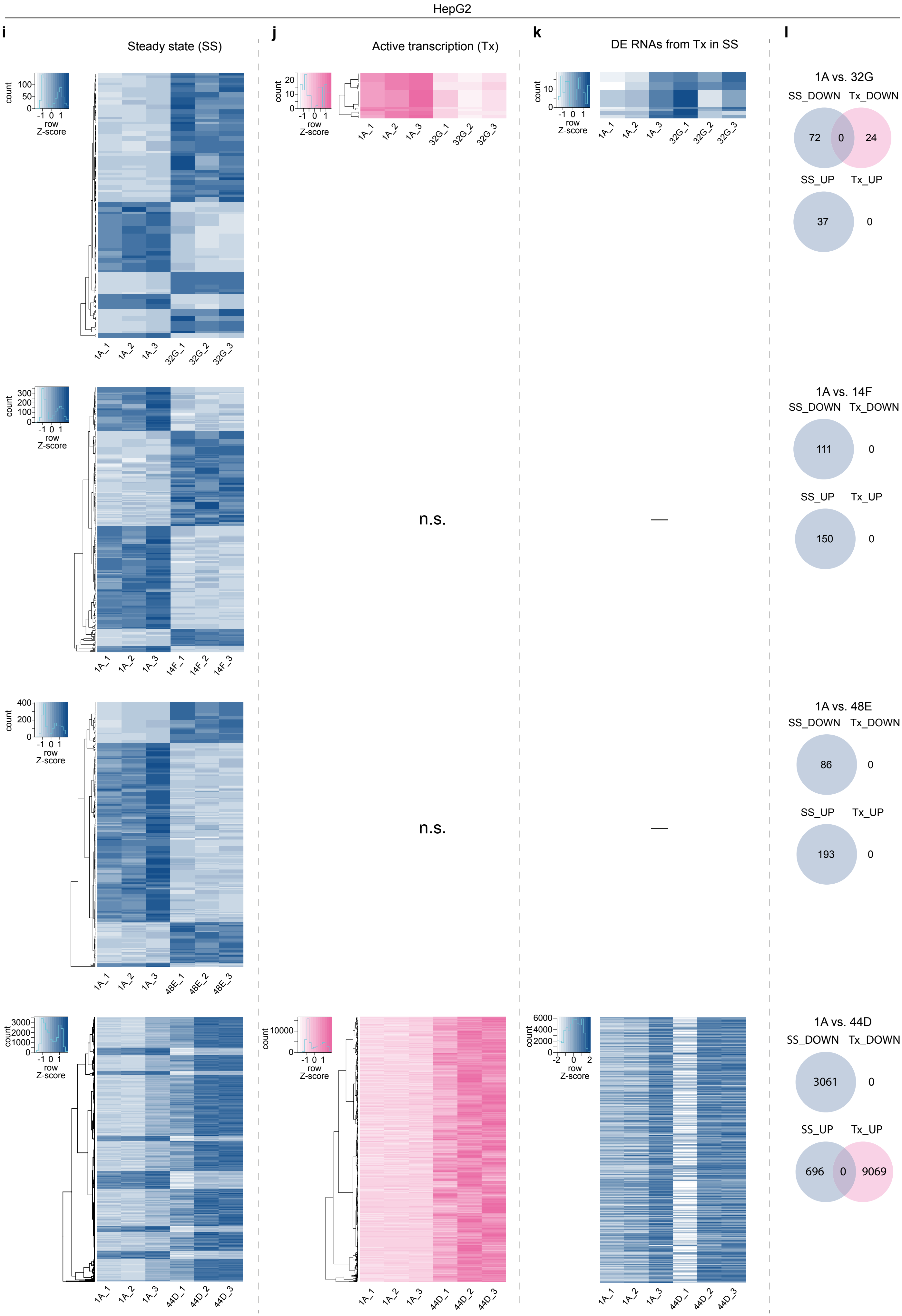
HepG2
i
Steady state (SS)
j
Active transcription (Tx)
k
DE RNAs from Tx in SS
l
count
row Z-score
1A_1
1A_2
1A_3
32G_1
32G_2
32G_3
14F_1
14F_2
14F_3
48E_1
48E_2
48E_3
44D_1
44D_2
44D_3
n.s.
n.s.
1A vs. 32G
SS_DOWN
Tx_DOWN
72
0
24
SS_UP
Tx_UP
37
0
1A vs. 14F
SS_DOWN
Tx_DOWN
111
0
SS_UP
Tx_UP
150
0
1A vs. 48E
SS_DOWN
Tx_DOWN
86
0
SS_UP
Tx_UP
193
0
1A vs. 44D
SS_DOWN
Tx_DOWN
3061
0
SS_UP
Tx_UP
696
0
9069

**Supplementary Fig. 10: SLAM-Seq analysis of HA1ER, HA1E and HepG2 diverging clonal systems.** Cells from the HA1ER (F12, 1F8, 1C9, 2A6 and 1D12) (**a-d**), HA1E (10E, A7A, A9H, B2F and B5F) (**e-h**) and HepG2 (1A, 14F, 44D, 48E and 32G) (**i-l**) clonal cell systems were subjected to SLAM-Seq analysis. **a, e, i.** Heatmaps represent the differential expression (FDR<0.01; FC>2) for each founder clone (F12 or 10E or 1A) vs individual subclones of steady-state RNA levels. Scale bar is shown. **b, f, j.** Heatmaps represent the differential expression (FDR<0.05; FC>2) for each for each founder clone (F12 or 10E or 1A) vs individual subclones of active transcription levels. Scale bar is shown. N.s. is shown when no differential expression was observed. **c, g, k.** Whenever differential expression is observed at the level of active transcription, differentially expressed transcript levels are retrieved at the steady-state level and plotted in a heatmap. **d, h, l.** Venn diagrams depicting the overlap between differentially expressed transcripts from active transcription vs steady-state RNA levels.

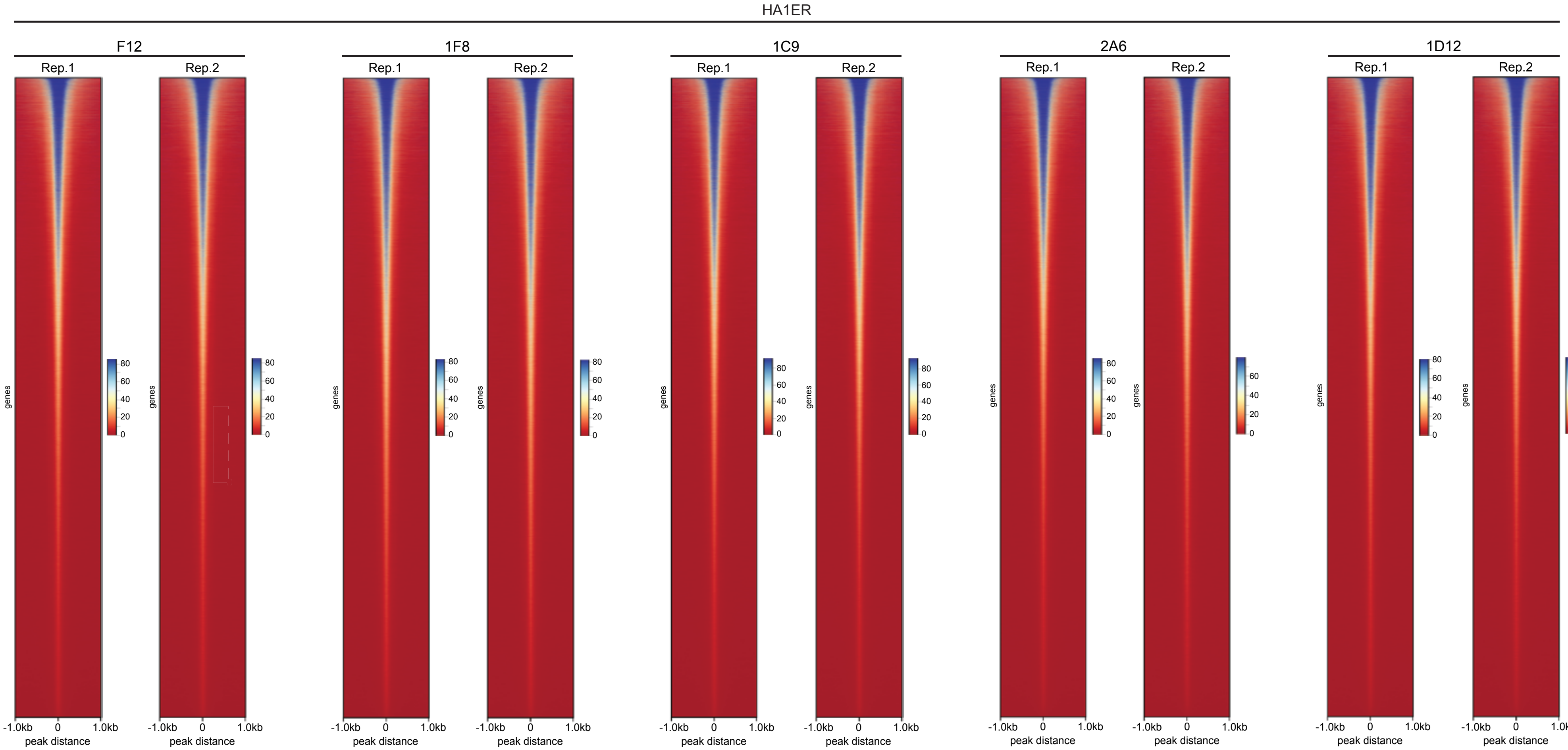

HA1ER
F12
1F8
1C9
2A6
1D12
Rep.1
Rep.2
genes
80
60
40
20
0
-1.0kb
0
1.0kb
peak distance

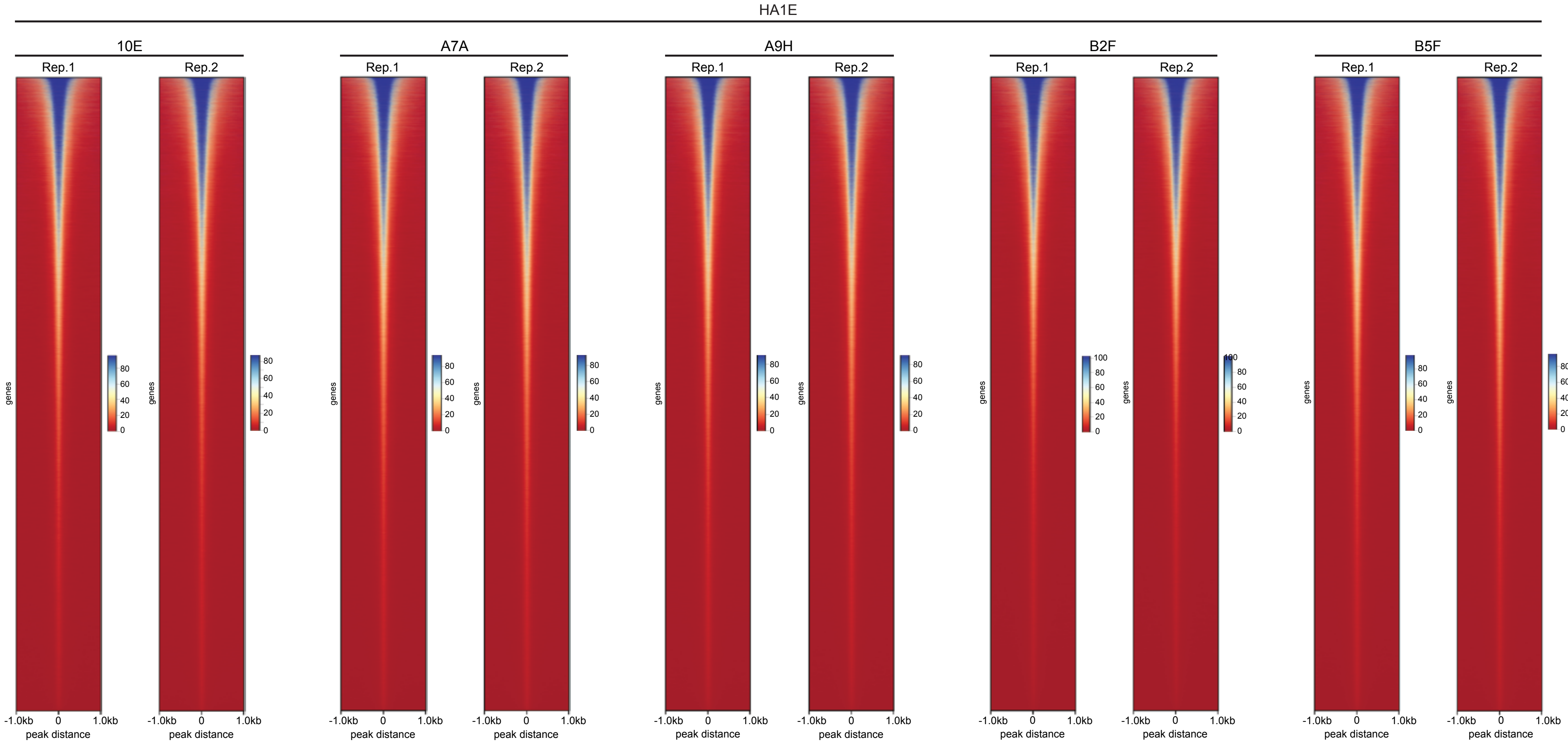
HA1E
10E
A7A
A9H
B2F
B5F
Rep.1
Rep.2
genes
0
20
40
60
80
100
-1.0kb
0
1.0kb
peak distance

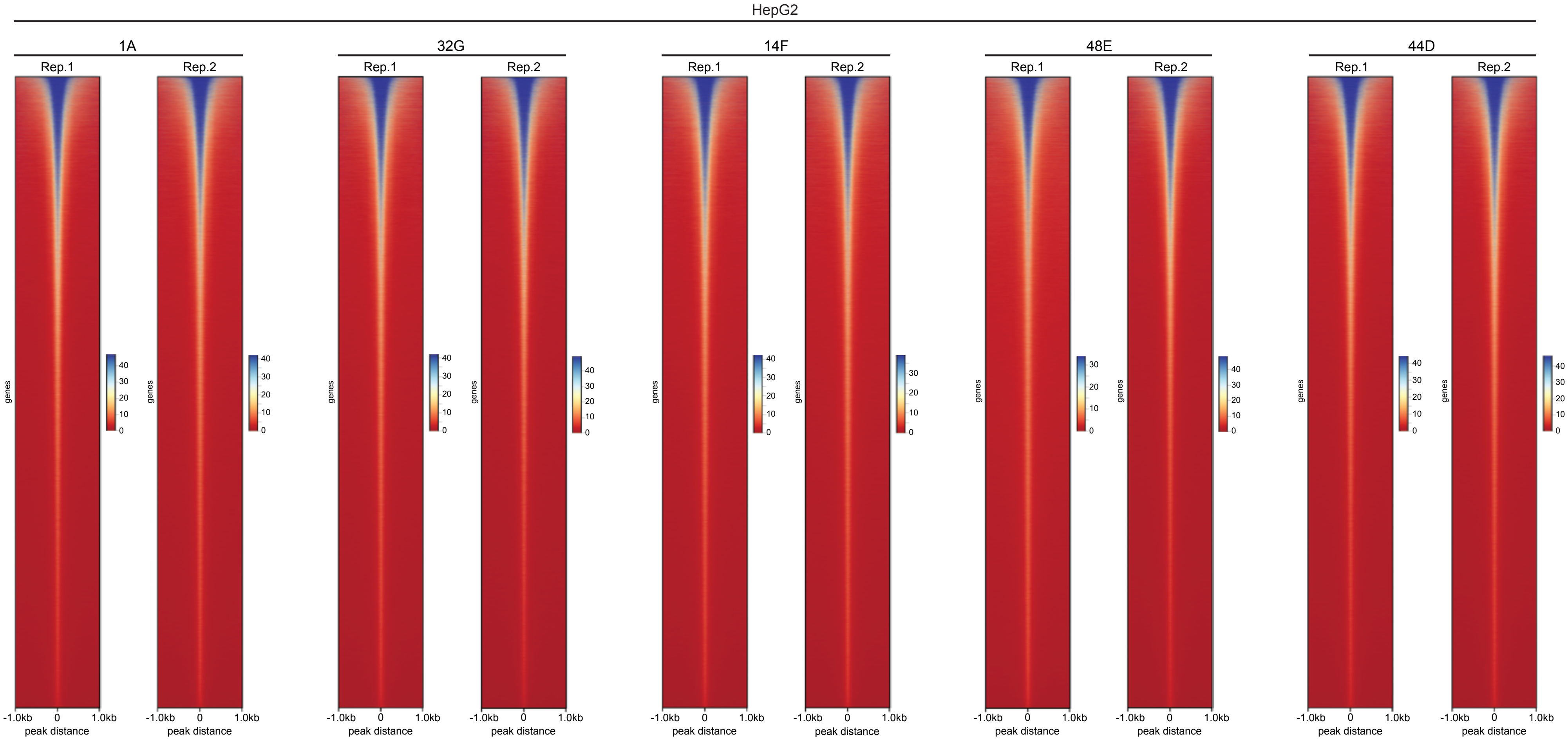
HepG2
1A
32G
14F
48E
44D
Rep.1
Rep.2
genes
peak distance
-1.0kb
0
1.0kb
0
10
20
30
40

**Supplementary Fig. 11: No major differences are observed in the accessible chromatin landscape between clonal cell populations diverging at the transcriptome level.** ATAC-Seq was performed on clonal populations from the HA1ER (F12, 1F8, 1C9, 2A6 and 1D12), HA1E (10E, A7A, A9H, B2F and B5F) and HepG2 (1A, 14F, 44D, 48E and 32G) systems. Two biological replicates per clone were analysed. Heatmaps displaying the identified peaks as determined by MACS using FDR<0.05 for each replicate is depicted. Peaks distance ± 1Kb. Scale bar is depicted.

a

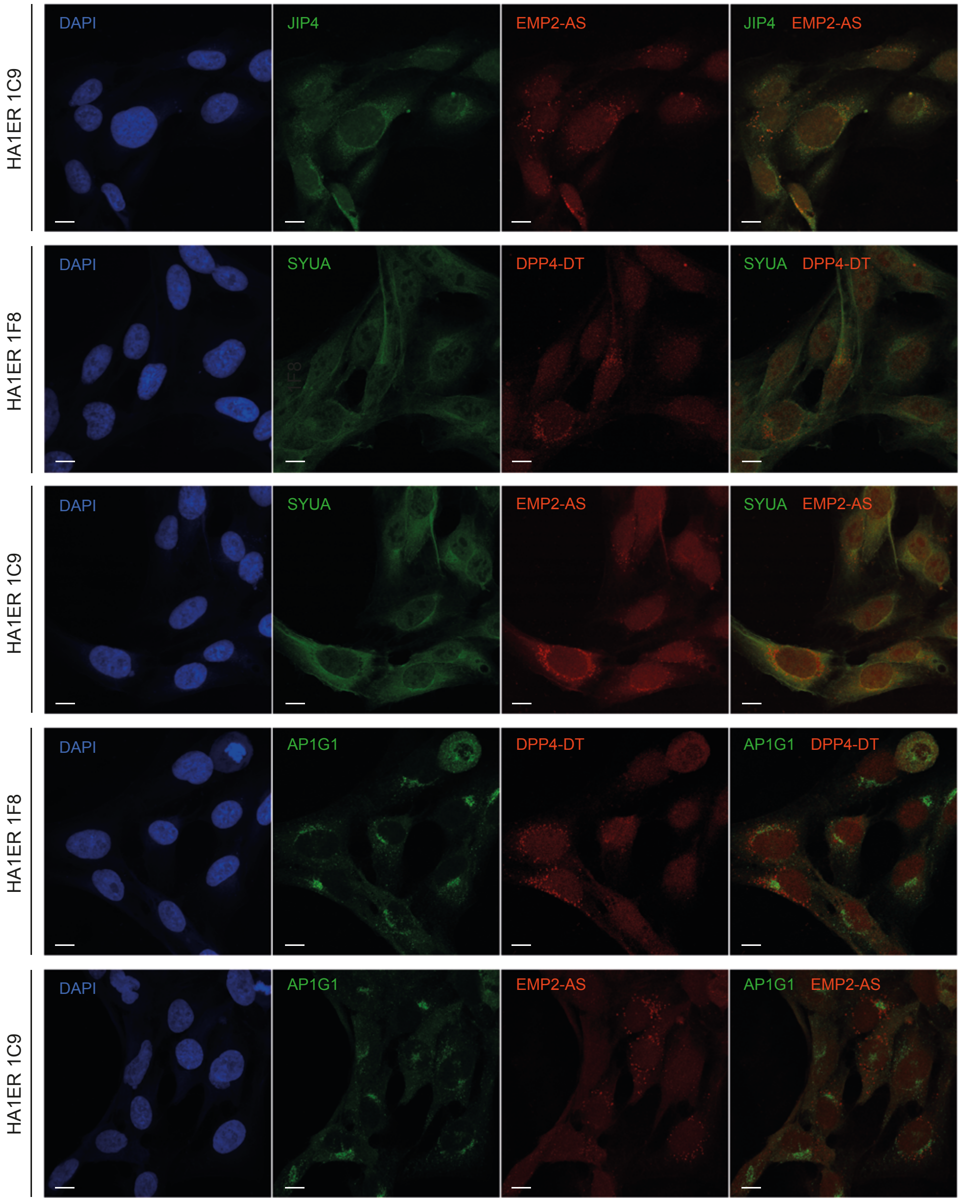
HA1ER 1C9
HA1ER 1F8
HA1ER 1C9
HA1ER 1F8
HA1ER 1C9
DAPI
JIP4
EMP2-AS
JIP4 EMP2-AS
SYUA
DPP4-DT
SYUA DPP4-DT
SYUA EMP2-AS
AP1G1
AP1G1 DPP4-DT
AP1G1 EMP2-AS

b

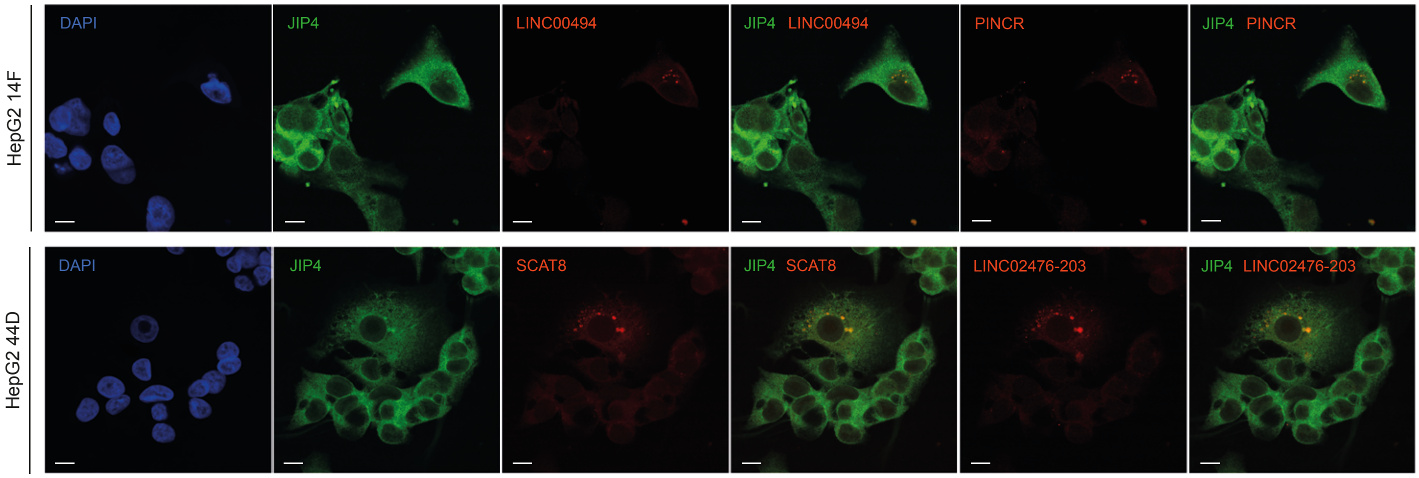
HepG2 14F
HepG2 44D
DAPI
JIP4
LINC00494
JIP4 LINC00494
PINCR
JIP4 PINCR
SCAT8
JIP4 SCAT8
LINC02476-203
JIP4 LINC02476-203

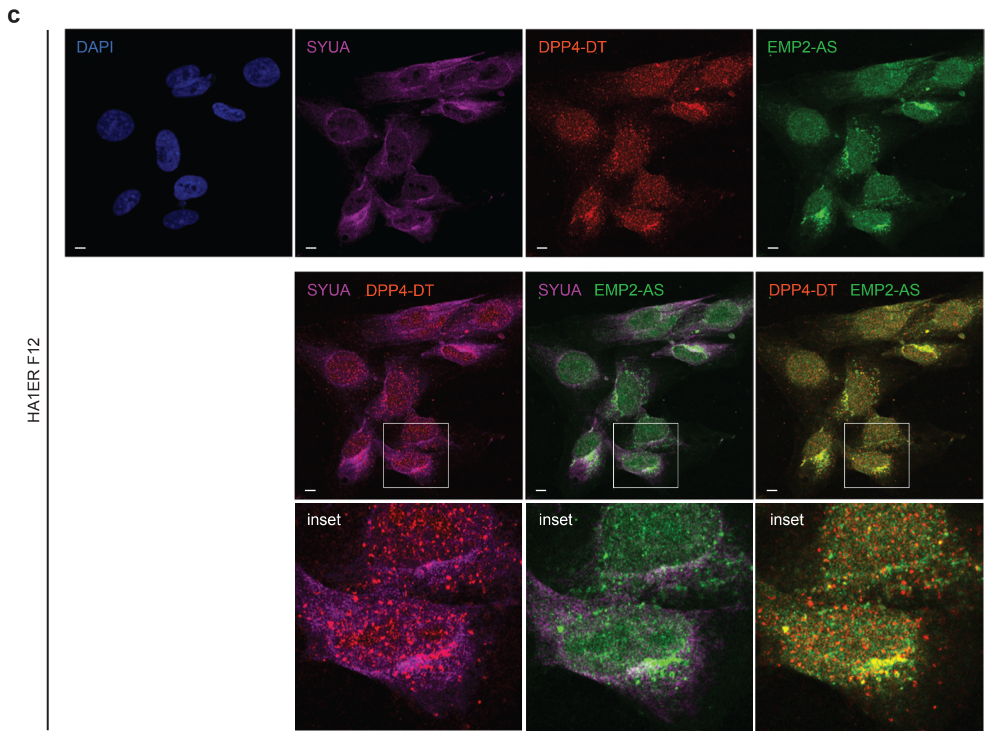


**Supplementary Fig. 12: IDR-containing proteins co-localise with clonally diverging lncRNAs. a.** Cells from the HA1ER clonal system (1F8 and 1C9) were processed for immunocytochemistry for IDR-containing proteins (JIP4, SYUA and AP1G1) followed by RNA-FISH for EMP2-AS and DPP4-DT. Evaluated proteins are pseudo-coloured in green and lncRNAs in red. Overlay images (IDR-proteins/RNAs) are depicted that are also shown in Fig. 5a. DAPI channel is shown. Bar 10 µm. **b.** Cells from the HepG2 clonal system (14F and 44D) were processed for immunocytochemistry for the IDR-containing proteins JIP4 followed by RNA-FISH for LINC00494, PINCR, SCAT8 and LINC02476-203. JIP4 is pseudo-coloured in green and lncRNAs in red. Overlay images (IDR-proteins/RNAs) are depicted. DAPI channel is shown. Bar 10 µm. **c.** Cells from the HA1ER parental clone F12 were subjected to immunocytochemistry for IDR-containing protein SYUA followed by co-labelling lncRNAs DPP4-DT and EMP2-AS as RNA-FISH specificity controls. Evaluated SYUA protein is pseudo-coloured in magenta and lncRNAs DPP4-DT and EMP2-AS in red and green, respectively. Overlay images (SYUA/lncRNAs and both lncRNAs) are depicted below, together with an inset. DAPI channel is shown. Bar 10 µm. Note that the signals from both lncRNAs evaluated do not always co-localise.

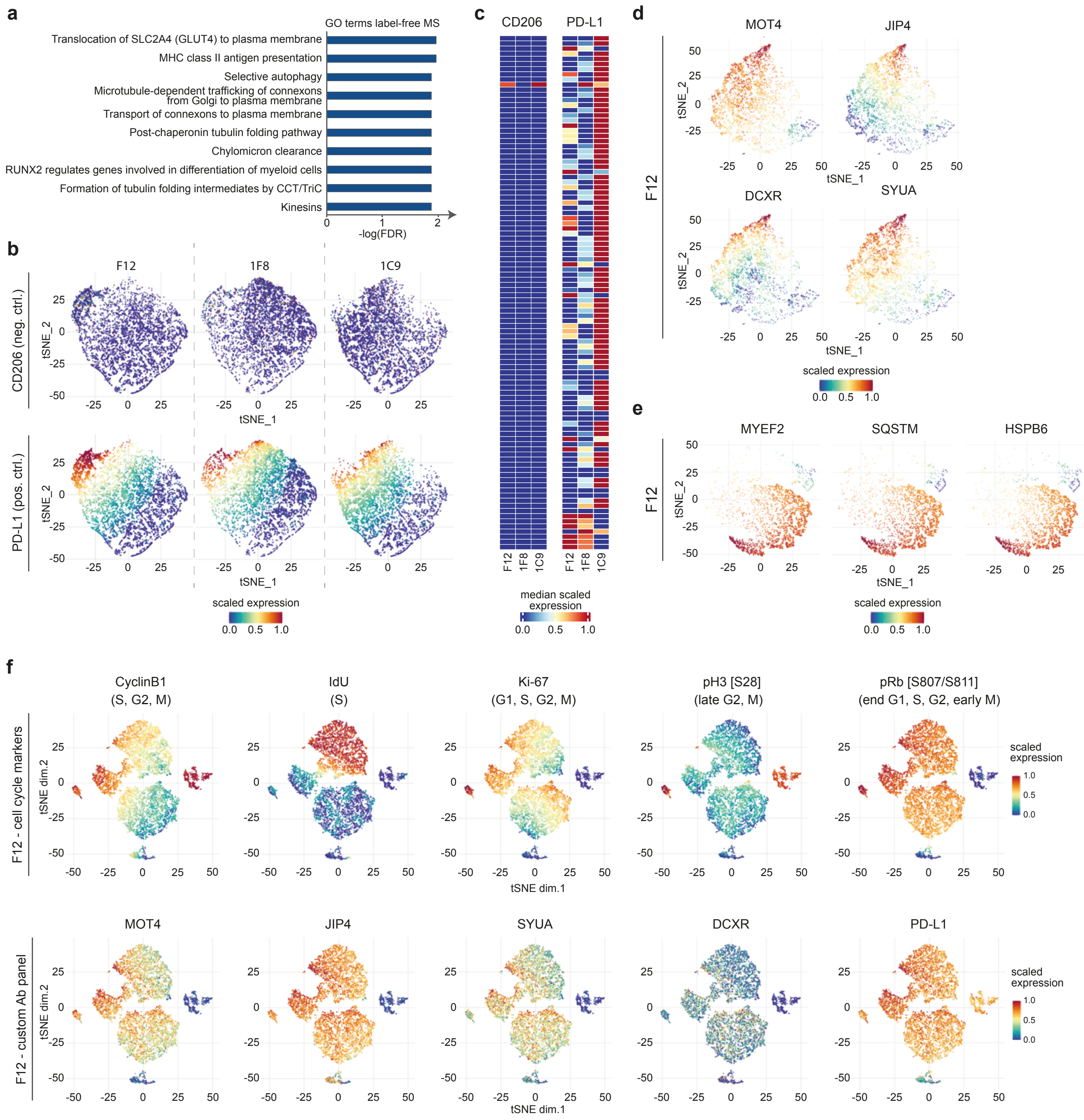


**Supplementary Fig. 13: IDR-containing proteins denote divergent subpopulations. a.** GO enrichment analysis (Reactome; FDR<0.05) of 1F8 vs 1C9 differentially expressed proteins identified by label free mass spectrometry. Top 10 pathways are shown. **b.** Mass cytometry (CyTOF) of cells from the HA1ER clonal system (F12, 1F8 and 1C9) using specific antibodies. tSNE plots represent the levels of CD206 (negative control) and PD-L1 (positive control) evaluated using specific antibodies in F12, 1F8 and 1C9. Scale bar is depicted. **c.** Heatmap represents the median scaled clustered expression of CD206 and PD-L1 antibodies in F12, 1F8 and 1C9. Scale bar is depicted. **d.** Mass cytometry (CyTOF) of cells from the HA1ER clonal system (F12) using specific antibodies for MOT4, JIP4, DCXR and SYUA. tSNE plots represent the levels of each antibody in the clonal population. Scale bar is depicted. **e.** Mass cytometry (CyTOF) of cells from the HA1ER clonal system (F12) using specific antibodies for MYEF2, SQSTM and HSPB6. tSNE plots represent the levels of each antibody in the clonal population. Scale bar is depicted. **f.** Cell cycle analysis (Cyclin B1, IdU, Ki-67, pH3 and pRb [S807/S811]) was concomitantly performed with MOT4, JIP4, SYUA (all IDR-containing) and DCXR (non-IDR) by mass cytometry on the HA1ER genetic anchor F12. tSNE plots represent the levels of each antibody in the clonal population. Scale bar is depicted.

a

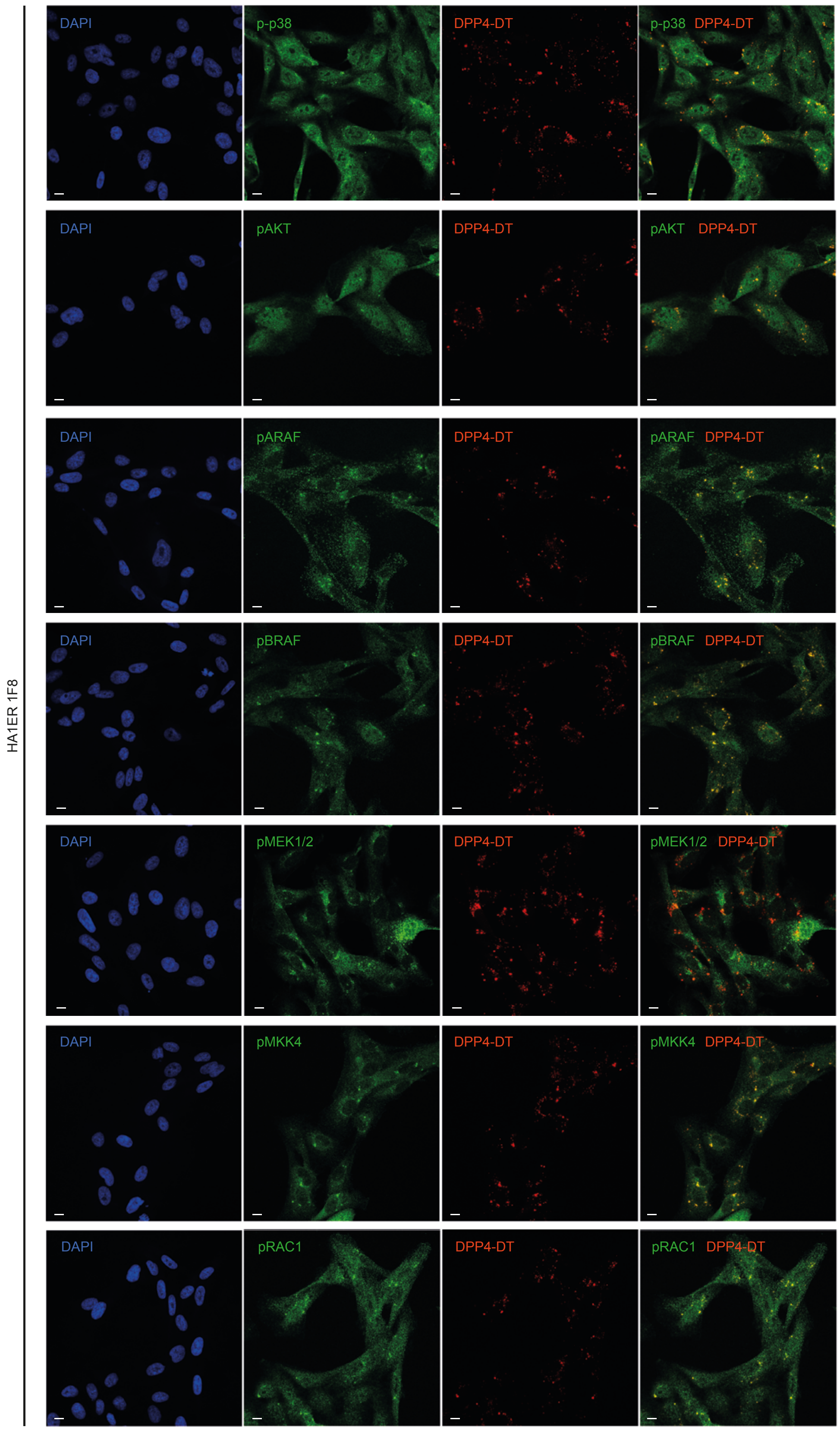

HA1ER 1F8
DAPI
p-p38
DPP4-DT
p-p38 DPP4-DT
pAKT
pAKT DPP4-DT
pARAF
pARAF DPP4-DT
pBRAF
pBRAF DPP4-DT
pMEK1/2
pMEK1/2 DPP4-DT
pMKK4
pMKK4 DPP4-DT
pRAC1
pRAC1 DPP4-DT

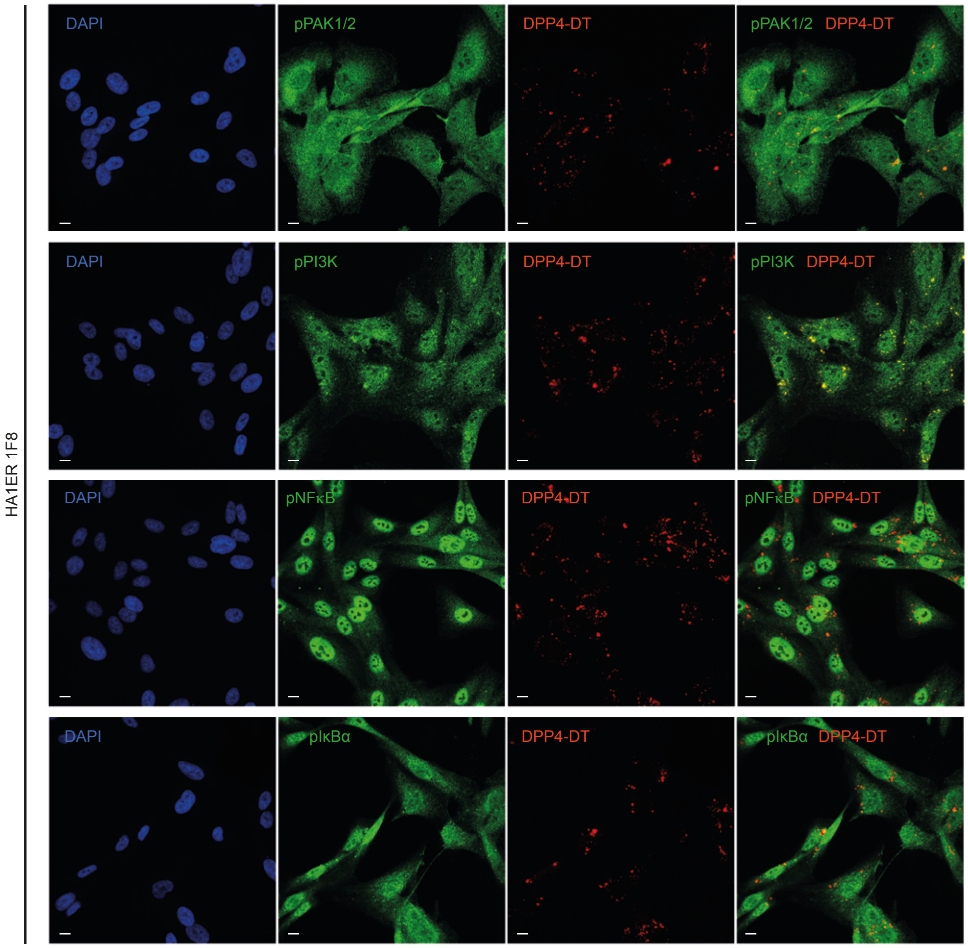
HA1ER 1F8
DAPI
pPAK1/2
DPP4-DT
pPAK1/2 DPP4-DT
DAPI
pPI3K
DPP4-DT
pPI3K DPP4-DT
DAPI
pNFκB
DPP4-DT
pNFκB DPP4-DT
DAPI
pIκBα
DPP4-DT
pIκBα DPP4-DT

b

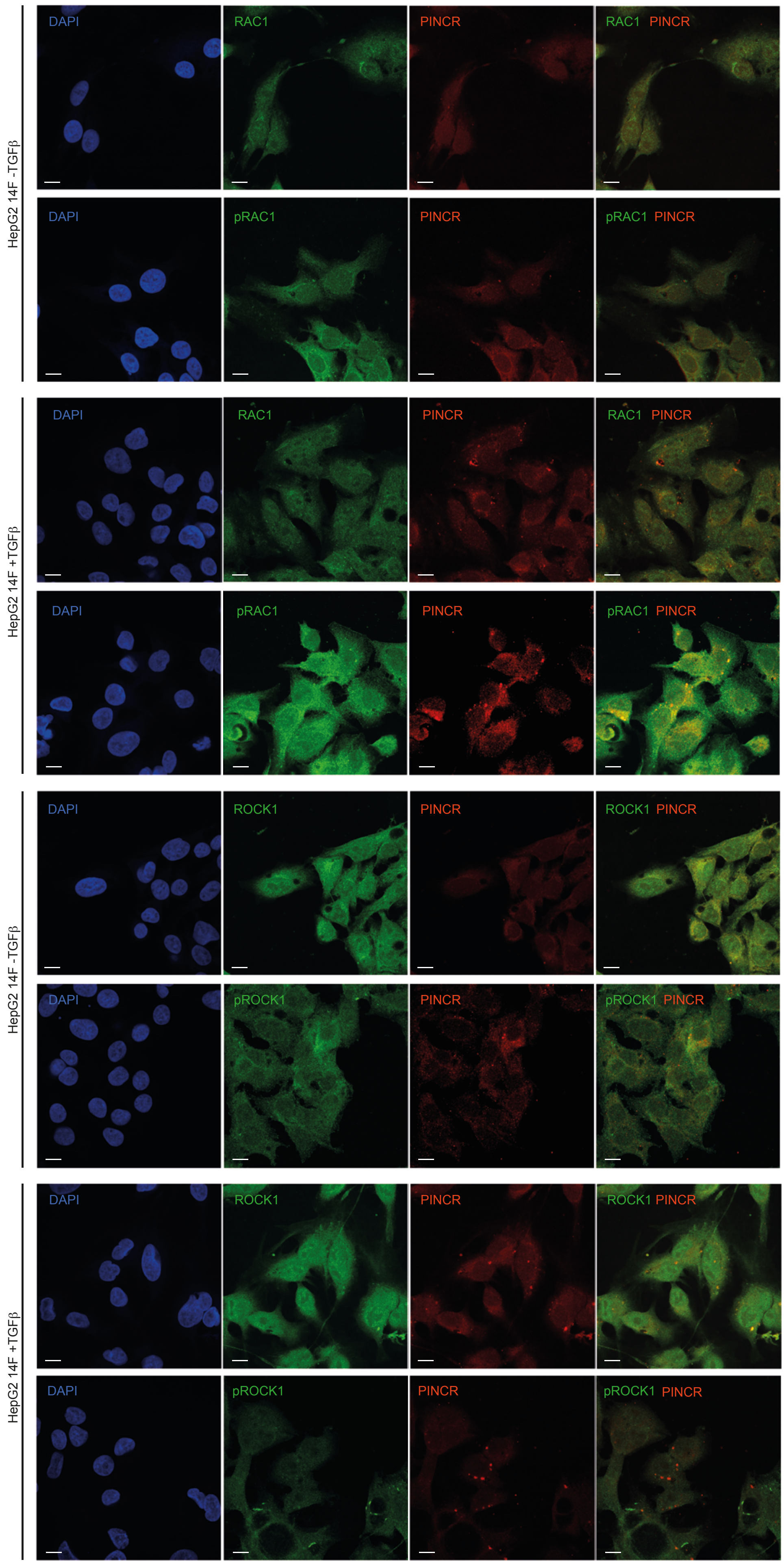
HepG2 14F -TGFβ
DAPI
RAC1
PINCR
RAC1 PINCR
pRAC1
pRAC1 PINCR
HepG2 14F +TGFβ
ROCK1
ROCK1 PINCR
pROCK1
pROCK1 PINCR

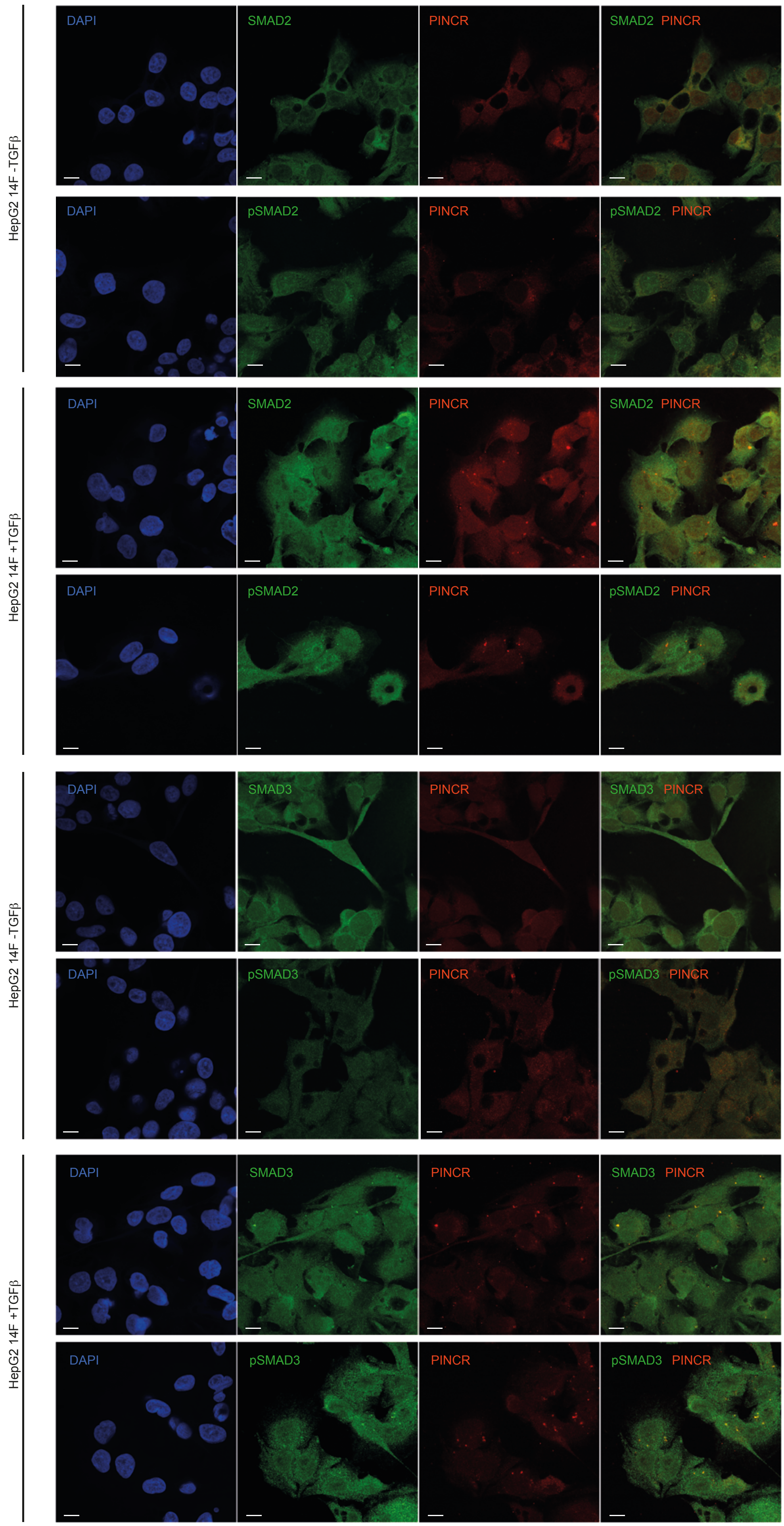
HepG2 14F -TGFβ
HepG2 14F +TGFβ
HepG2 14F -TGFβ
HepG2 14F +TGFβ
DAPI
SMAD2
PINCR
SMAD2 PINCR
pSMAD2
pSMAD2 PINCR
SMAD3
SMAD3 PINCR
pSMAD3
pSMAD3 PINCR

**Supplementary Fig. 14: Activated signalling proteins co-localise with clonally diverging lncRNAs. a.** HA1ER clone 1F8 was subjected to immunocytochemistry for activated signalling pathways (p-p38, pAKT, pARAF, pBRAF, pMEK1/2, pMKK4, pRAC1, pPAK1/2, pPI3K, pNFκB and pIκBα) coupled to RNA-FISH targeting DPP4-DT lncRNA. Panels depict individual channels (DAPI, signalling pathway protein and DPP4-DT lncRNA). Merged images are displayed (signalling protein/DPP4-DT) which are also shown in Fig. 5b. Bar 10 µm. **b.** HepG2 clone 14F was subjected to immunocytochemistry for TGFβ-dependent activated signalling pathways (RAC1, pRAC1, ROCK1, pROCK1, SMAD2, pSMAD2, SMAD3 and pSMAD3) coupled to RNA-FISH targeting PINCR lncRNA in cells either vehicle treated (HepG2 -TGFβ) or TGFβ treated (HepG2 +TGFβ). Panel depicts individual channels (DAPI, signalling pathway protein and PINCR lncRNA). Merged images are displayed (signalling protein/PINCR) which are also depicted in Fig. 5c. Bar 10 µm.

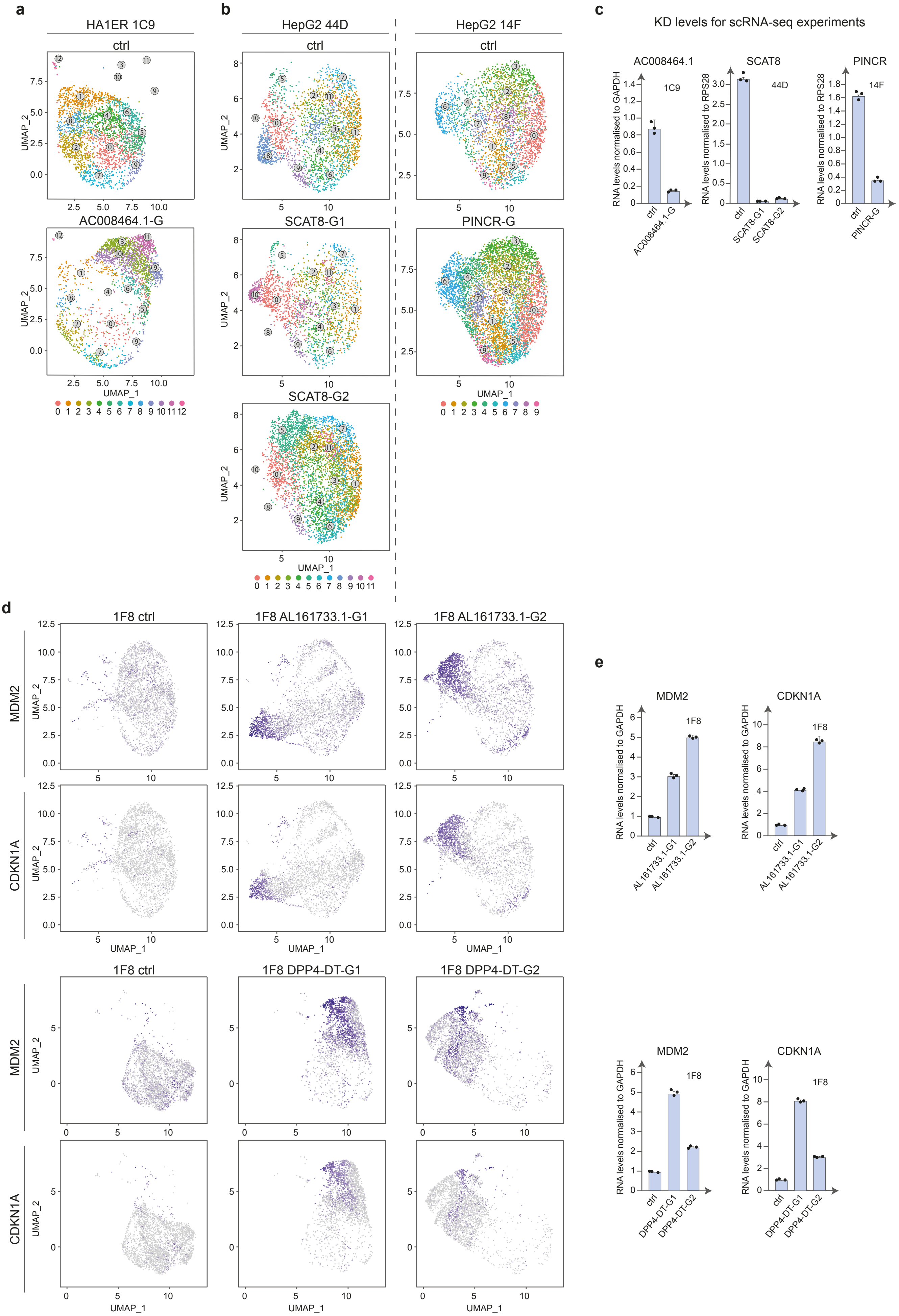
a
HA1ER 1C9
ctrl
AC008464.1-G
UMAP_1
UMAP_2
b
HepG2 44D
ctrl
SCAT8-G1
SCAT8-G2
HepG2 14F
ctrl
PINCR-G
c
KD levels for scRNA-seq experiments
AC008464.1
1C9
SCAT8
44D
PINCR
14F
RNA levels normalised to GAPDH
RNA levels normalised to RPS28
ctrl
AC008464.1-G
SCAT8-G1
SCAT8-G2
PINCR-G
d
1F8 ctrl
1F8 AL161733.1-G1
1F8 AL161733.1-G2
MDM2
CDKN1A
1F8 DPP4-DT-G1
1F8 DPP4-DT-G2
e
MDM2
CDKN1A
1F8
AL161733.1-G1
AL161733.1-G2
DPP4-DT-G1
DPP4-DT-G2

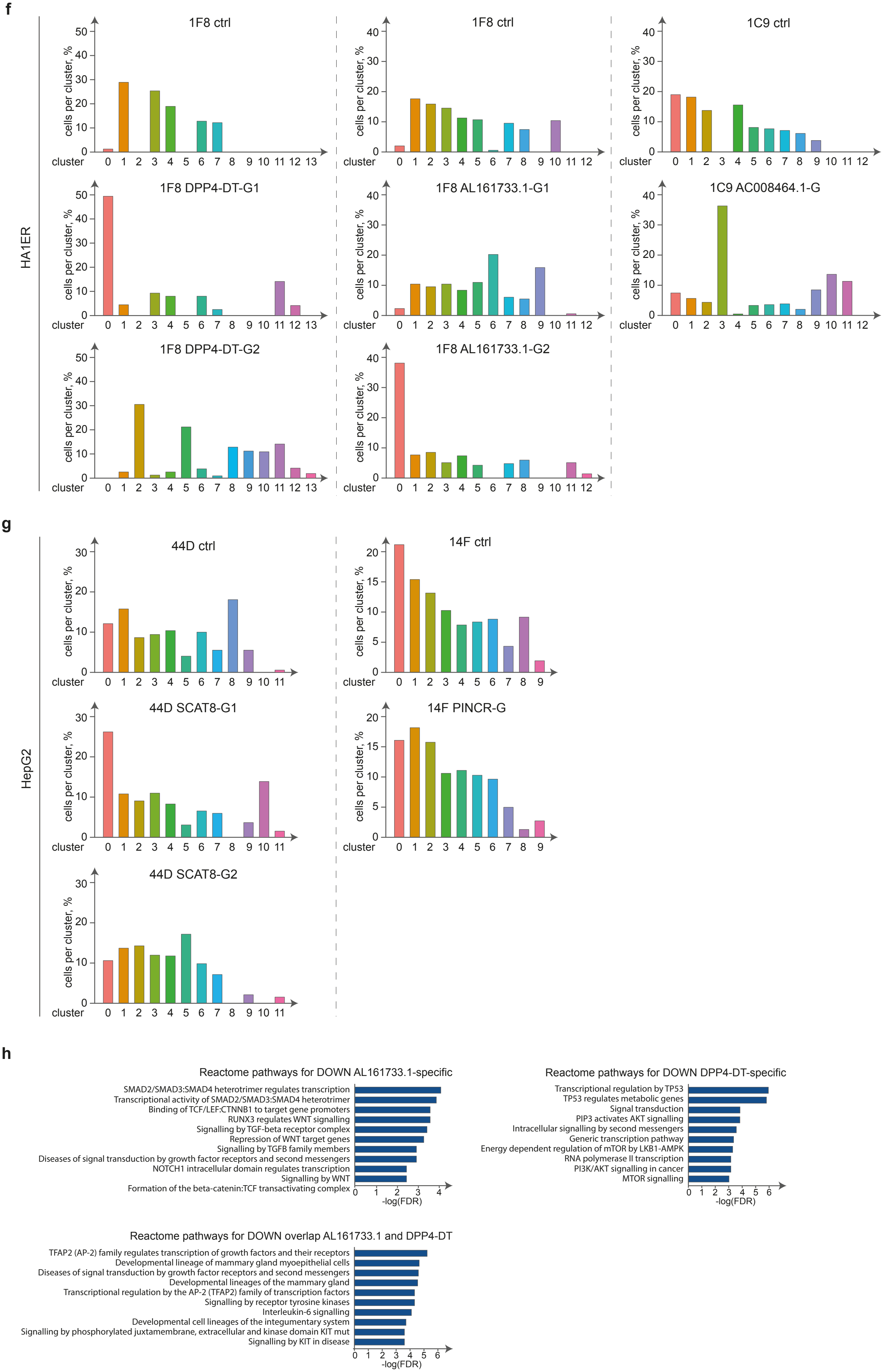
f
HA1ER
1F8 ctrl
1F8 ctrl
1C9 ctrl
1F8 DPP4-DT-G1
1F8 AL161733.1-G1
1C9 AC008464.1-G
1F8 DPP4-DT-G2
1F8 AL161733.1-G2
cells per cluster, %
cluster
g
HepG2
44D ctrl
14F ctrl
44D SCAT8-G1
14F PINCR-G
44D SCAT8-G2
cells per cluster, %
cluster
h
Reactome pathways for DOWN AL161733.1-specific
SMAD2/SMAD3:SMAD4 heterotrimer regulates transcription
Transcriptional activity of SMAD2/SMAD3:SMAD4 heterotrimer
Binding of TCF/LEF:CTNNB1 to target gene promoters
RUNX3 regulates WNT signalling
Signalling by TGF-beta receptor complex
Repression of WNT target genes
Signalling by TGFB family members
Diseases of signal transduction by growth factor receptors and second messengers
NOTCH1 intracellular domain regulates transcription
Signalling by WNT
Formation of the beta-catenin:TCF transactivating complex
-log(FDR)
Reactome pathways for DOWN DPP4-DT-specific
Transcriptional regulation by TP53
TP53 regulates metabolic genes
Signal transduction
PIP3 activates AKT signalling
Intracellular signalling by second messengers
Generic transcription pathway
Energy dependent regulation of mTOR by LKB1-AMPK
RNA polymerase II transcription
PI3K/AKT signalling in cancer
MTOR signalling
-log(FDR)
Reactome pathways for DOWN overlap AL161733.1 and DPP4-DT
TFAP2 (AP-2) family regulates transcription of growth factors and their receptors
Developmental lineage of mammary gland myoepithelial cells
Diseases of signal transduction by growth factor receptors and second messengers
Developmental lineages of the mammary gland
Transcriptional regulation by the AP-2 (TFAP2) family of transcription factors
Signalling by receptor tyrosine kinases
Interleukin-6 signalling
Developmental cell lineages of the integumentary system
Signalling by phosphorylated juxtamembrane, extracellular and kinase domain KIT mut
Signalling by KIT in disease
-log(FDR)

**Supplementary Fig. 15: Gapmers targeting diverging lncRNAs promote transcriptome remodelling and rearrangement of signalling pathways. a.** HA1ER clone 1C9 was subjected to antisense-mediated knock-down (Gapmer) for AC008464.1 lncRNA and further processed for scRNA-Seq. UMAP plots depict transcriptome states in control (non-targeting control gapmer) and upon AC008464.1-G knockdown. **b.** HepG2 clones 44D and 14F were subjected to antisense-mediated knock-down (Gapmer) for SCAT8 and PINCR lncRNAs respectively and further processed for scRNA-Seq. UMAP plots depict transcriptome states in control (non-targeting control gapmer) and upon SCAT8 or PINCR knockdown. Two alternative antisense molecules targeting different regions of the SCAT8 transcript (SCAT8-G1 and SCAT8-G2) were used and one for PINCR. **c.** lncRNA knockdown was evaluated for cells processed for scRNA-Seq shown in **a** and **b** (AC008464.1, HA1ER clone 1C9; SCAT8, HepG2 clone 44D; PINCR, HepG2 clone 14F). Transcript levels were determined by reverse transcription followed by qPCR (RT-qPCR) and were normalised to GAPDH (HA1ER) or RPS28 (HepG2). Histograms represent the mean ± propagated error of 3 technical replicates. Individual data points are depicted. **d.** UMAP plots depicting the convergent expression of mRNAs (MDM2 and CDKN1A) resulting from the knockdown of AL161733.1 or DPP4-DT lncRNAs in HA1ER 1F8 clone. Two alternative antisense molecules targeting different regions of the AL161733.1 lncRNA (AL161733.1-G1 and AL161733.1-G2) and two for DPP4-DT lncRNA (DPP4-DT-G1 and DPP4-DT-G2) were used. **e.** The induction of the upregulated transcripts identified upon DPP4-DT and AL161733.1 knockdown in the HA1ER clone 1F8 (MDM2 and CDKN1A) was evaluated by RT-qPCR prior to the RPPA experiment presented in Fig. 6b. Validation of efficient lncRNA knockdowns are shown in Fig. 6e. Histograms represent the mean expression (normalised to GAPDH) ± propagated error of 3 technical replicates. **f.** Cells from HA1ER system were subjected to lncRNA knockdown (clone 1F8 - DPP4-DT and AL161733.1; clone 1C9 - AC008464.1) or control treatment (non-targeting control gapmer, ctrl) and processed for scRNA-Seq. Histograms represent percentage of cells in each cluster for each sample. Antisense gapmers are denoted as G, G1 or G2. **g.** Cells from HepG2 system were subjected to lncRNA knockdown (clone 44D – SCAT8; clone 14F - PINCR) and processed for scRNA-Seq. Histograms represents percentage of cells in each cluster for each clone. Antisense gapmers are denoted as G, G1 or G2. **h.** Pathway analysis (Reactome) on HA1ER clone 1F8 upon DPP4-DT or AL161733.1 lncRNAs knockdown (FDR<0.01). Top 10 downregulated pathways are shown for antibody signals specifically downregulated (≥20% decrease to control) upon DPP4-DT (DPP4-DT-specific) or AL161733.1 knock-down (AL161733.1-specific), as well as for downregulated signalling pathway members detected in both knock-downs (overlap).

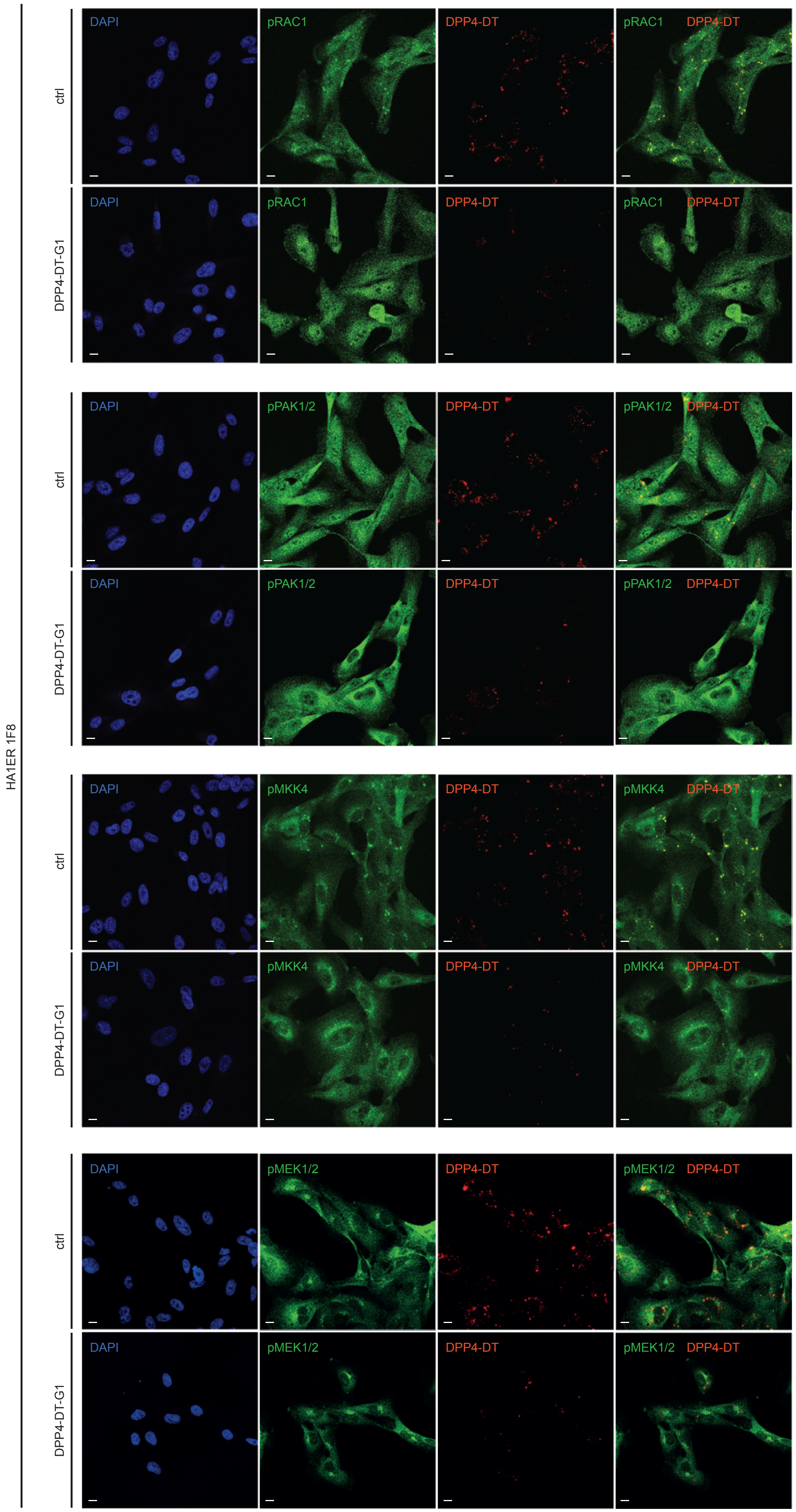
HA1ER 1F8
ctrl
DPP4-DT-G1
DAPI
pRAC1
DPP4-DT
pPAK1/2
pMKK4
pMEK1/2

**Supplementary Fig. 16: Gapmers targeting diverging lncRNA induce intracellular relocalisation of activated signalling pathways.** HA1ER clone 1F8 was subjected to antisense-mediated knock-down for DPP4-DT lncRNA (gapmer DPP4-DT-G1) or non-targeting control gapmer (ctrl) and further processed for immunocytochemistry for activated signalling pathways (pRAC1, pPAK1/2, pMKK4, pMEK1/2) coupled to RNA-FISH targeting DPP4-DT lncRNA. Panels depict individual channels (DAPI, signalling pathway protein (green) and DPP4-DT lncRNA (red)). Merged images are displayed (signalling protein/DPP4-DT) which are also shown in Fig. 7a. Bar 10 µm.
